# A precisely adjustable, variation-suppressed eukaryotic transcriptional controller to enable genetic discovery

**DOI:** 10.1101/2019.12.12.874461

**Authors:** Asli Azizoğlu, Roger Brent, Fabian Rudolf

## Abstract

Methods to express genes conditionally into phenotype remain central to biological experimentation and biotechnology. Current methods enable either on/off or imprecisely controlled graded gene expression. We developed a “well-tempered” controller, WTC_846_, for precisely adjustable, graded and growth condition independent conditional expression of genes in *Saccharomyces cerevisiae*. In WTC_846_ strains, the controlled genes are expressed from a strong, native promoter engineered to be repressed by the prokaryotic TetR protein and induced by tetracycline and analogues. A second instance of this promoter drives TetR itself. This autorepression loop exhibits low cell-to-cell variation in gene expression and allows precise adjustment of the steady state abundance of any protein with inducer. A second, constitutively expressed zeroing repressor abolishes basal expression in the absence of inducer. WTC_846_-controlled, stable (Cdc42, Tpi1) and unstable (Ipl1) proteins recapitulated known knockout and overexpression phenotypes. *WTC_846_::CDC20* strains enabled inducer regulated cell cycle synchronization. WTC_846_ alleles of *CDC28*, *TOR1*, *PBR1* and *PMA1* exhibited expected gene dosage-dependent growth rates and morphological phenotypes, and *WTC_846_::WHI5* strains exhibited inducer controlled differences in cell volume. WTC_846_ controlled genes comprise a new kind of “expression clamped” allele, for which variation in expression is minimized and gene dosage can be set by the experimenter across the range of cellular protein abundances. In yeast, we expect WTC_846_ alleles to find use in assessment of phenotypes now incompletely penetrant due to variable dosage of the causative protein, and in genome-wide epistasis screens. Implementation in higher cells should enable experiments now impossible due to cell-to-cell variation and imprecise control.

## Introduction

Since the spectacular demonstration of suppression of nonsense mutations and its application to T4 development,^1^ means to express genes conditionally into phenotype have remained central to biological experimentation and discovery. During the 20th century, workhorse methods to ensure the presence or absence of gene products have included use of ts and cs mutations within genes, for example to give insight into ordinality of cell biological events.^2^ After the advent of recombinant DNA methods, conditional expression of genes into proteins, for example by derepression of lac promoter derivatives,^3^ also found application in biotechnology for production of therapeutics and industrial products.^4^ In 2020, contemporaneous approaches to conditional expression in wide use include construction of transgenes activated by chimeric activators controlled by promoters whose expression is temporally and spatially restricted to different cell lineages,^5^ hundreds of approaches based on production of DNA rearrangements by phage-derived site specific recombination,^6^ and triggered induction of engineered genes by chimeric transcription regulators with DNA binding moieties based on derivatives of TetR from Tn10.^7, 8^ Most of these approaches are all-or-none, in the sense that they are not intended to bring about expression of intermediate levels of protein; and the observations they enable are often qualitative.

But it has long been recognized that adjustment of protein dosage can provide additional insight into function that cannot be gained from all-or-none expression. For example, controlled expression of the bacteriophage λ cI and cro gene products was key to understanding how changes in the level of those proteins regulated the phage’s decision to undergo lytic or lysogenic growth.^9–11^ In *S. cerevisiae*, contemporaneous means to tune dosage include metabolite induced promoters, such as *P_GAL1_*, *P_MET3_*, *P_CUP1_*,^12^ in which expression is controlled by control of growth media composition, and small molecule induced systems, such as the *β*-estradiol induced LexA-hER-B112 system.^13^ Many of these systems depend on fusions between eukaryotic and viral activator domains and prokaryotic proteins^8, 13–15^ that bind sites on engineered promoters.^16^ These methods suffer from a number of drawbacks, including basal expression when not induced,,^8, 13, 17^ deleterious effects on cell growth due to sequestration of cellular components by the activation domain^18^ induction of genes in addition to the controlled gene,^14^ and high cell-to-cell variability in expression of the controlled genes.^13, 19, 20^

These inducible systems rely on “activation by recruitment”,^21^ the activator binds a site on DNA upstream of a yeast gene and recruits general transcription factors and regulators of the Pre-Initiation Complex (PIC). These assemble downstream at the “core promoter”, and recruit RNA polymerase II to induce transcription.^22^ An alternative to inducible activation would be to engineer reversible repression of yeast transcription by prokaryotic repressors.^23–25^ For TATA containing promoters, binding of prokaryotic proteins such as LexA and the lac repressor near the TATA sequence can repress transcription,^23, 26, 27^ presumably by interference with the formation of the PIC, transcription initiation, or early elongation. It has long been recognized^28^ that prokaryotic repressors likely work through different mechanisms than mechanisms used by repressors native to eukaryotes.^29, 30^

We envisioned that an ideal conditional expression system to support genetic and quantitative experimentation would: 1) function in all growth media, 2) be inducible by an exogenous small molecule with minimal other effects on the cell, 3) manifest no basal expression of the controlled gene in absence of inducer, allowing generation of null phenotypes, 4) enable a very large range of precisely adjustable expression 5) drive very high maximum expression, allowing generation of overexpression phenotypes. Moreover, since differences in global ability to express genes into proteins^31^ lead to differences in allelic penetrance and expressivity,^32^ the ideal controller should 6) exhibit low cell-to-cell variability at any set output, facilitating detection of phenotypes that depend on thresholds of protein dosage, and other inferences of single cell behaviors from population responses.

Here we describe the development of a prokaryotic repressor-based transcriptional controller of gene expression, Well-tempered Controller_846_ (WTC_846_), that fulfils the criteria outlined above. This development had three main stages. We first engineered a powerful eukaryotic promoter that is repressed by the prokaryotic repressor TetR and induced by the chemical tetracycline and its analogue anhydrotetracycline (aTc), to use as the promoter of the controlled gene. Next, we used instances of this promoter to construct a configuration of genetic elements that show low cell-to-cell variation in expression of the controlled gene, by creating an autorepression loop in which TetR repressed its own synthesis. Third and last, we abolished basal expression of the controlled gene in the absence of the inducer, by engineering a weakly expressed “zeroing” repressor, a chimera between TetR and an active yeast repressor Tup1. With WTC_846_, adjusting the extracellular concentration of aTc can precisely set the expression level of the controlled gene in different growth media, over time and over cell cycle stage. The gene is then “expression clamped” with low cell-to cell variability at a certain protein dosage, which can range from undetectable to greater abundance than wild type. We showed that strains carrying WTC_846_ allelic forms of essential genes recapitulated known knockout and overexpression phenotypes. We constructed strains bearing WTC_846_ alleles of genes involved in size control, growth rate, and cell cycle state and showed that these allowed precise experimental control of these fundamental aspects of cell physiology. We expect that WTC_846_ alleles will find use in biological engineering and in discovery research, in assessment of phenotypes now incompletely penetrant due to cell-to-cell variability of the causative gene, in hypothesis-directed cell biological research, and in genome-wide studies such as gene by gene epistasis screens.

## Results

### Construction of a repressible *P_TDH3_* promoter

Our goal was to engineer efficient repression of eukaryotic transcription by a bacterial repressor. We started with a strong,^33^ well characterized, constitutive, and endogenous yeast promoter. This promoter, *P_TDH3_*, has three key Transcription Factor (TF) binding sites, one for Rap1 and two for Gcr1^34^ in its Upstream Activating Region (UAS), and a TATA sequence at which Pre-Initiation Complex (PIC) assembles on the core promoter (Figure 1A). Based on earlier work, we knew that binding of prokaryotic repressors to sites flanking the TATA sequence of *P_TDH3_* repressed activity of this promoter,^27^ presumably by interfering with PIC formation, transcription initiation, or early elongation. We therefore placed well characterized, 15bp long TetR binding sites (*tetO_1_*)^35^ immediately upstream and downstream of the *P_TDH3_* TATA sequence to create *P_2tet_*. To determine whether repressor binding could also block function in the UAS, we placed a single *tetO_1_* directly upstream of each Rap1 and Gcr1 binding site to create *P_3tet_*. We also combined the operators in these constructs to generate *P_5tet_* (Figure 2B). We integrated a single copy^36^ of constructs bearing these promoters directing the synthesis of the fluorescent protein Citrine into the LEU2 locus.^37^

**Figure 1:**
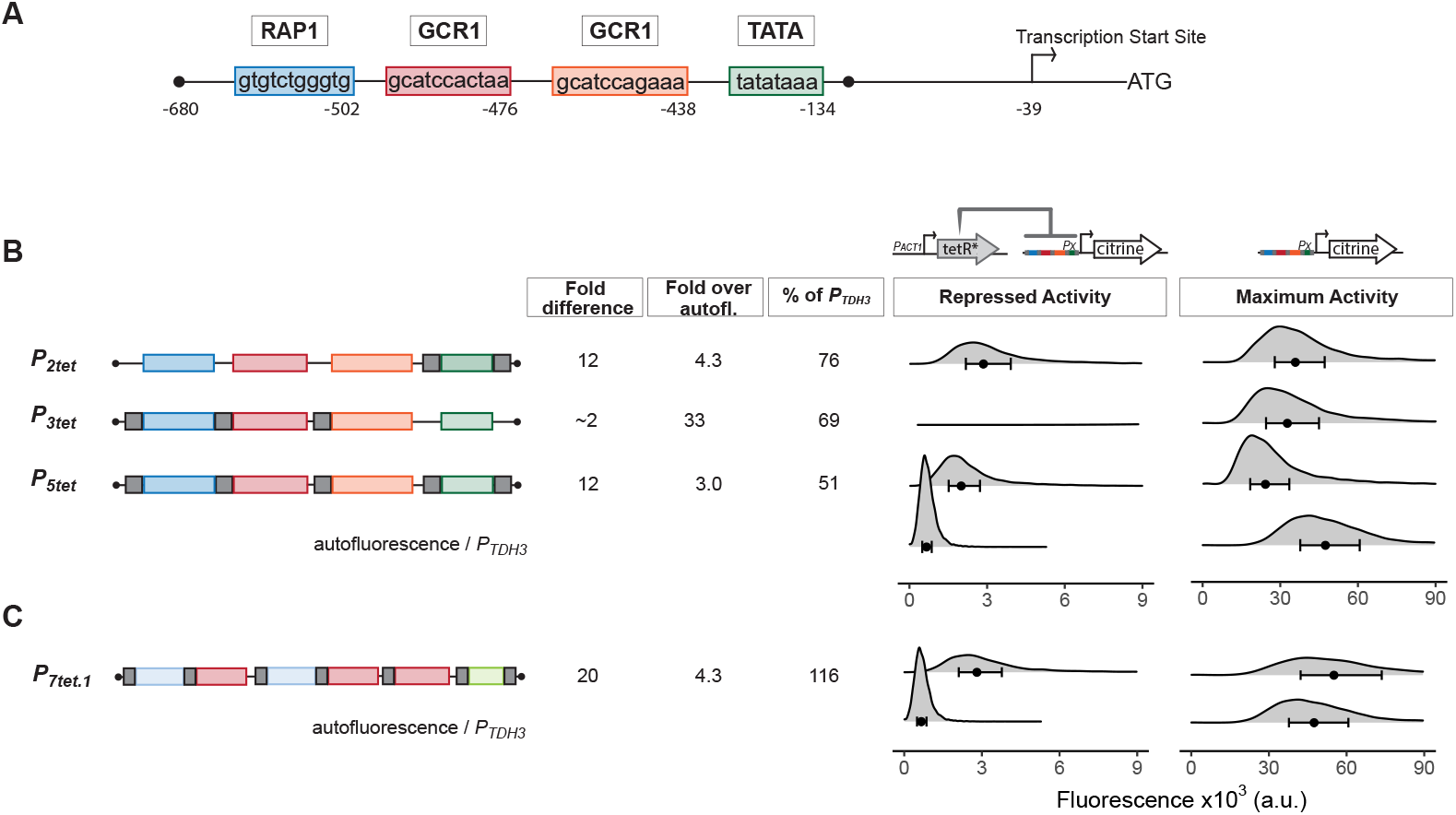
Repression of engineered *P_TDH3_* derivatives by TetR. **A) Structure of the starting promoter, *P_TDH3_***. Diagram shows the nucleotide positions of the binding sites for the endogenous transcription factors Rap1 and Gcr1, the TATA-sequence, and the transcription start site relative to the start codon of the *TDH3* gene. **B) Repression and maximum activity of engineered *P_TDH3_* derivatives**. Diagrams above the plots display the genetic elements of strains used in B&C. Left diagram depicts strains used to test repressed activity, right diagram maximum activity. Px denotes any TetR repressible promoter. The * in TetR indicates a SV40 Nuclear Localization Sequence. In all strains, the *P_TDH3_* derivative promoters diagrammed on the left directed the synthesis of Citrine integrated into the LEU2 locus. Grey boxes inside the diagrams denote *tetO_1_* TetR binding sites. For measurement of repressed activity, an additional *P_ACT1_* directed TetR was integrated into the HIS3 locus. Citrine fluorescent signal was detected by flow cytometry. Fold difference refers to the median of the maximum activity divided by the median of the repressed activity signal. Fold over autofluorescence refers to median repressed activity signal divided by the median autofluorescent background signal. Maximum promoter activity is quantified as percentage of *P_TDH3_* signal using the medians. x axis shows intensity of fluorescence signal. Plots are density distributions of the whole population, such that the area under the curve equals 1 and the y axis indicates the proportion of cells at each fluorescence value. The circles inside each density plot show the median, and the upper and lower bounds of the bar correspond to the first and third quartiles of the distribution. **C) Repression and maximum activity of the optimized *P_7tet.1_***. Diagrams and plots as in (B). *P_7tet.1_* contained additional binding sites for Rap1 and Gcr1 selected for higher activity, as well as an alternative TATA sequence as described in the Supplementary Information. It shows the highest fold difference, maximum activity comparable to *P_TDH3_*, and low repressed activity.

**Figure 2:**
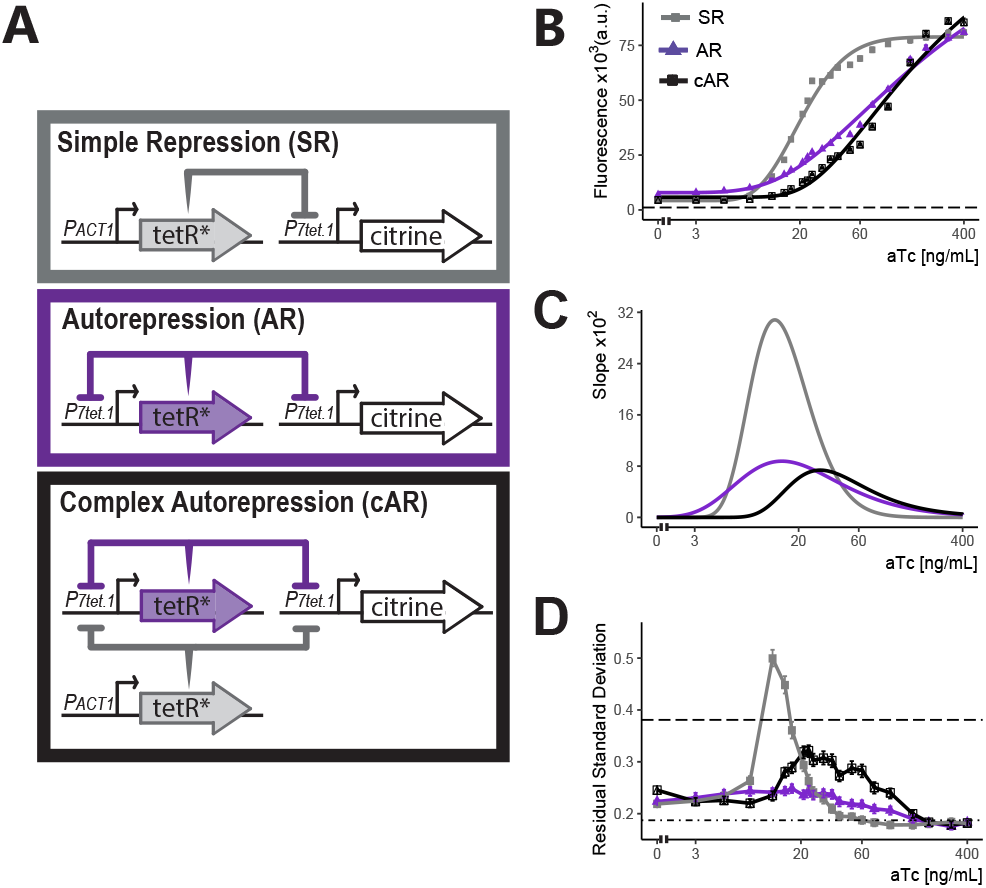
Comparison of the three controller architectures. **A) Genetic elements of the different controller architectures used in these experiments**. The * next to TetR indicates SV40 Nuclear Localization Sequence and flat headed arrows indicate repression. In all cases, *P_7tet.1_* drives Citrine expression integrated at the LEU2 locus. In SR, the repressor of *P_7tet.1_*, TetR, is integrated at the HIS3 locus and is constitutively expressed. In AR, tetR is again integrated at the HIS3 locus, but is now expressed by *P_7tet.1_*. cAR has the same constructs as AR and an additional, constitutively expressed zeroing repressor integrated at the URA3 locus. **B) aTc dose response curves of Citrine expression for the three different architectures**. Citrine fluorescence from strains bearing these architectures was measured at steady state using flow cytometry after 7 hours of induction with different concentrations of aTc. Circles indicate the median fluorescence at each dose. Lines are fitted using a 5-parameter log logistic function as explained in Materials&Methods. Dashed line indicates autofluorescence signal measured from the parental strain without Citrine. **C) Slopes of the dose response curves in (B)**. The x axis range with non-zero slopes define the useful input dynamic range. **D) Cell-to-cell variation of expression of these three architectures**. We measured and calculated single-reporter CCV as described. Higher RSD values mean more cell-to-cell variation in the population. Dot-dash line indicates the CCV of the strain where Citrine is constitutively expressed from *P_TDH3_* and dashed line indicates CCV of autofluorescence in the parent strain without Citrine. Error bars indicate 95% confidence interval calculated using bootstrapping (n=1000) as described in Materials&Methods.

We compared the Citrine fluorescence signal (measured by flow cytometry at wavelengths 515-545nm) from these promoters to quantify their activity. We compared the strains FRY2551, FRY2564, and FRY2566 with an otherwise-isogenic strain in which Citrine was expressed from native *P_TDH3_* (FRY2683). This fluorescence signal measures Citrine expression, but also includes autofluorescent background from the yeast cells. We quantified this background by using the otherwise-isogenic parent strain FRY70. Measured in this way, *P_2tet_* had 76%, *P_3tet_* 69%, and *P_5tet_* 51% of *P_TDH3_* activity (Figure 1B). To assess repressibility of these promoters, we compared Citrine expression in these strains with expression in otherwise-isogenic strains in which a genomically integrated *P_ACT1_* promoter drove constitutive expression of TetR (FRY2562, FRY2573, FRY2577). By this measure, TetR repressed *P_2tet_* by a factor of 12, *P_3tet_* by a factor of 1.5, and *P_5tet_* by a factor of 12 (Figure 1B & Figure S1). Absolute repressed signal from these promoters was 4.3, 33 and 3 times the autofluorescence background. Because our aim was to create a promoter with no expression when repressed, we viewed even small reductions in repressed expression as useful and therefore decided to use *P_5tet_* as a basis for further constructions.

Insertion of *tetO_1_* sites in *P_TDH3_* to create *P_5tet_* had reduced promoter maximum activity considerably. In order to regain the lost activity, we tested numerous constructs to find optimal placement for the *tetO_1_* sites, optimized Rap1, Gcr1 and TATA sequences, and increased the number of Rap1 and Gcr1 sequences (see Supplementary Text & Figure S21). This work resulted in *P_7tet.1_*, which carried two Rap1 and three Gcr1 sites, sequence optimized to generate higher promoter activity, and an alternative TATA sequence to that of *P_TDH3_*. By the assays described above, the new promoter *P_7tet.1_* (FRY2661) showed comparable maximum expression to *P_TDH3_*, 20-fold repression of Citrine signal, and absolute repressed activity of 4.3 fold over background (FRY2663 & Figure 1C). We chose *P_7tet.1_* as the promoter to develop our controller with.

### Complex autorepressing (cAR) controller architecture expands the input dynamic range and reduces cell-to-cell variation

We set out to optimize control of genes by *P_7tet.1_*. To do so, we tested the ability of different constructions that directed the synthesis of TetR to regulate *P_7tet.1_*-Citrine fluorescence signal. Figure 2A shows the three different architectures. In Simple Repression (SR), the *P_7tet.1_* controlled gene was repressed by TetR expressed from a constitutive promoter. In Autorepression (AR), the *P_7tet.1_* controlled gene was repressed by TetR expressed from a second instance of *P_7tet.1_*, therefore creating a negative feedback loop. In Complex Autorepression (cAR), a second TetR gene expressed from a constitutive promoter was added to the AR architecture.

We compared the input-output relationship (i.e. dose response) for the three architectures. To do so, we constructed otherwise-isogenic strains with these architectures in which *P_7tet.1_* directed Citrine expression (FRY2663, FRY2674, and FRY2741). We used flow cytometry to quantify Citrine fluorescence signal from all strains seven hours after addition of different concentrations of anhydrotetracycline (aTc) and fitted a log logistic model to the median fluorescence (See Materials & Methods) (Figure 2B&C).

Compared to the SR architecture, the AR architecture showed a more gradual dose response curve and a larger input dynamic range (the range of input doses for which the slope of the dose response curve was non-zero), from 3-400ng/mL vs. 5-80 ng/mL aTc. This same flattening of the response curve and increased input dynamic range in autorepressing, synthetic TetR based eukaryotic systems has been described,^38^ and we believe it operates in evolved prokaryotic systems including Tn10 and the *E. coli* SOS regulon, in which the TetR and LexA repressors repress their own synthesis (see discussion). A broader input dynamic range allows more precise adjustment of protein levels, since small differences in inducer concentration (due for example to experimental errors, or differences in aTc uptake among cells) have smaller effects.

In these experiments, we also measured cell-to-cell variation (CCV) in the expression of the controlled gene. Many existing inducible gene expression systems show considerable variation in expression of the controlled gene, making it difficult to achieve homogenous phenotypes at the population level.^13, 19, 20^ In *S. cerevisiae* and *C. elegans*, comparison of signals from strains with different constellations of reporter genes allows quantification of different sources of variation in protein dosage.^31, 39, 40^ Here, we quantified overall variation in protein dosage by measuring the Coefficient of Variation (CoV) in fluorescent output from a single reporter (Figure S2), and we developed a second measure (explained in SI) that normalized variation in dosage with respect to a key confounding variable, cell volume, to correct for its effect on protein concentration. In this, we measured the Residual Standard Deviation (RSD) in signal from cells after normalization of output for volume estimated by a vector of forward and side scatter signals (Figure 2D & Figure S3). By both measures, strains carrying the SR architecture showed high variation throughout the input dynamic range, with a peak around the mid-point (12ng/mL aTc). Strains bearing the AR architecture showed low overall CCV, and no peak at intermediate aTc concentrations. This diminution of CCV in synthetic, autorepressing TetR based eukaryotic systems has previously been described.^38, 41^ In the SR architecture, variations in the amount of TetR in different cells cannot be buffered. In the AR architecture, such variations in repressor concentration are corrected for (see discussion) and variation in expression of the controlled gene is at or around the same level as seen for constitutive expression driven by a number of native promoters (see Figure S5 for variability of commonly used promoters). This reduced cell-to-cell variation is useful for inferring single cell behaviors by observing population level responses (see discussion).

Compared with cells bearing the SR architecture, otherwise-isogenic cells bearing the AR architecture showed increased basal expression (6.3 vs. 4.1-fold over autofluorescence background). The increased basal expression was a consequence of the fact that in the AR architecture *P_7tet.1_* directs the synthesis of both the controlled Citrine gene and of TetR itself, and the steady state abundance of TetR in the cell is lower than in cells in which synthesis of TetR is driven by *P_ACT1_*. To further lower basal expression, we constructed strains with a third architecture, cAR, in which an additional constitutive promoter drove expression of a second TetR gene. Compared to otherwise-isogenic AR strains, strains expressing Citrine controlled by the cAR architecture showed reduced basal expression (4.1-fold over autofluorescence), but retained the reduced CCV and the more gradual dose response (Figure 2C&D and S4). We therefore picked this cAR architecture for our controller.

### Hybrid repressor abolishes basal expression of *P_7tet.1_*

To further decrease basal expression in the cAR architecture, we set out to create a more effective TetR derivative. Initially we followed an approach that increased the size and nuclear concentration of TetR by fusing it to other inert bacterial proteins and nuclear localization sequences, but this approach was not enough to abolish all basal expression (see Supplementary Text).

*P_3tet_* bears *tetO_1_* sites only in its UAS. The fact that *P_3tet_* SR strains only showed weak repression (1.5 fold) suggested that TetR, and other inert derivatives described in the SI, exerted their effects on *P_7tet.1_* mostly by their action at the *tetO_1_* sites flanking the TATA sequence. We thus hypothesized that TetR derivatives that carried native, active yeast repressors might more effectively repress from sites in the UAS. The yeast repressor Tup1 complexes with Ssn6 (also called Cyc8) with a ratio of 4:1, forming a complex of 420kDa,^42^ and this complex represses transcription through a number of mechanisms. These include repositioning and stabilizing nucleosomes to form an inacessible chromatin structure.^43–45^ Tup1 also blocks chromatin remodeling, masks activation domains, and excludes TBP.^44, 46, 47^ LexA-Tup1 fusion proteins repress transcription when bound upstream of the Cyc1 promoter,^48^ and TetR-Tup1 fusions reduce uninduced expression in a dual TetR activator-repressor controller.^17^ For *P_7tet.1_*, we imagined that as many as seven TetR-Tup1 dimers might bind to the promoter, potentially recruiting two additional Tup1 and one Ssn6 molecules per *tetO_1_* site. The resulting ~3mDa of protein complexes might block activation by one or more of the above mechanisms. We therefore measured the ability of a TetR-nls-Tup1 fusion to repress *P_7tet.1_*-driven Citrine signal in SR strains. When its expression was directed from *P_ACT1_* (FRY2669), TetR-nls-Tup1 decreased uninduced fluorescence signal to background levels (Figure 3A). Because fusion of TetR to a mammalian repressor domain in mammalian cells had shown very slow induction kinetics,^49^ we checked whether the TetR-nls-Tup1 fusion showed increased induction time compared to TetR alone but found no such effect (Figure S6). Additionally, TetR-nsl-Tup1 abolished uninduced expression driven by *P_3tet_* (Figure S7) (77-fold repression), compared to repression of otherwise isogenic strains by TetR, which showed basal expression reduced by 1.5 fold (Figure S1). By contrast, TetR-nls-Tup1 fusion repressed *P_2tet_*, where *tetO_1_* flank only the TATA sequence, more strongly than TetR alone, but still showed basal expression. Our data thus suggested that the TetR-nls-Tup1 suppressed basal expression mainly by its effects in the UAS (see Discussion).

**Figure 3:**
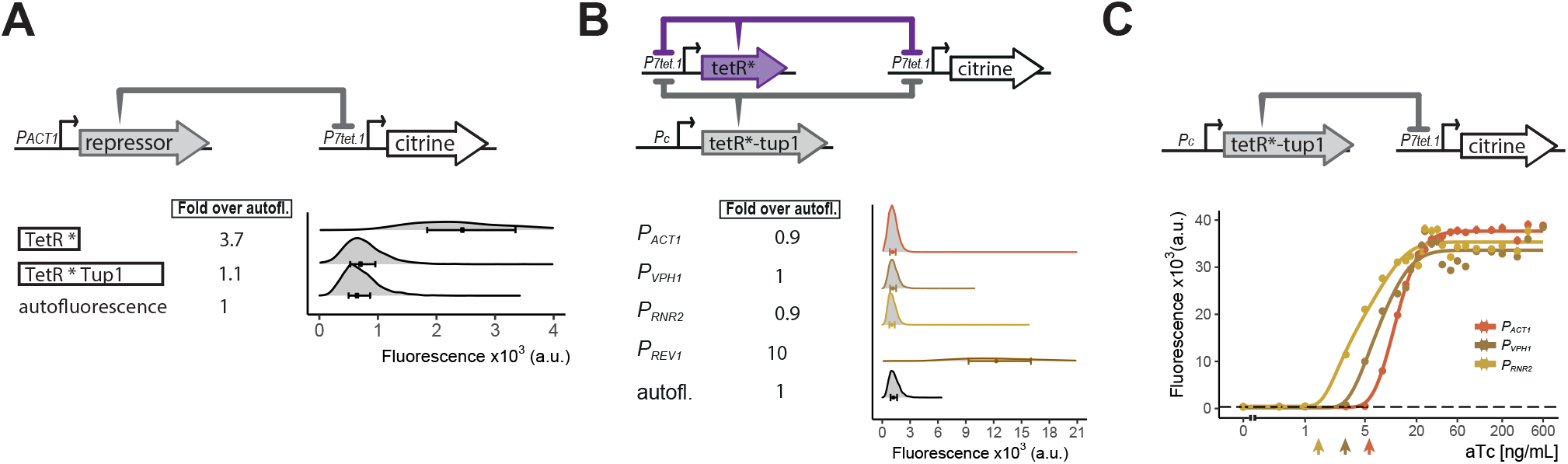
Repressor optimization to abolish *P_7tet.1_* basal expression. **A) Testing repression by the TetR-Tup1 fusion**. The top diagram indicates the genetic elements of the SR architecture used to test the ability of the TetR-Tup1 fusion to abolish basal expression from *P_7tet.1_*. Diagrams to the left of the plot show the different repressors used. Each * indicates one SV40 Nuclear Localization Sequence. For both (A) and (B), Citrine fluorescence from *P_7tet.1_* repressed by the repressors indicated was measured using flow cytometry. Plots as in Figure 1. The circles inside each density plot show the median and the upper and lower bounds of the bar correspond to the first and third quartiles of the distribution. Numbers to the left of the plot indicate fold expression over autofluorescence, i.e. the median of the Citrine fluorescence detected divided by the median of the autofluorescence signal. **B) Finding the lowest expression level of the zeroing repressor TetR-nls-Tup1 that abolishes basal expression in *P_7tet.1_* cAR**. The top diagram shows the genetic elements of the cAR architecture in the strains tested. Pc indicates a constitutive promoter. Promoters driving TetR-nls-Tup1 expression are indicated to the left of the plot. Numbers to the left of the plot as in (A).**C) Reducing expression of TetR-nls-Tup1 lowers induction threshold**. The top diagram shows genetic elements of SR architecture in which synthesis of TetR-nls-Tup1 was directed by different promoters. The plot shows Citrine fluorescence measured using flow cytometry at steady state, 7 hours after induction with different aTc concentrations. Arrows indicate induction thresholds, defined as the lowest aTc dose where an increase in fluorescence signal was detected. Dashed line indicates autofluorescence control (parent strain without Citrine), circles indicate the median of the experimentally measured population, lines are fitted. Error bars indicate 95% confidence interval calculated using bootstrapping (n=1000) as explained in Materials&Methods.

In the cAR architecture, the induction threshold, i.e. the smallest concentration of inducer that can induce expression, is determined by the number of molecules of the repressors present before induction. We sought to lower the induction threshold in order to maximize the input dynamic range. Therefore, we constructed cAR controllers using TetR and TetR-nls-Tup1, to determine the lowest level of TetR-nls-Tup1 that could still abolish uninduced expression from *P_7tet.1_*. TetR-nls-Tup1 was driven by constitutive promoters of genes whose products were of decreasing abundance^33^ (*P_ACT1_*, *P_VPH1_*, *P_RNR2_*,*P_REV1_*) (FRY2673, FRY2684, FRY2749, and FRY2715). The *P_ACT1_*, *P_VPH1_* and *P_RNR2_* strains showed no uninduced expression, while the *P_REV1_* strain did (Figure S8). Out of the three, Rrn2 protein is present at lower abundance, and the *P_RNR2_* driven TetR-nls-Tup1 has the lowest induction threshold in a dose response experiment with strains bearing SR architectures (FRY2669, 2676, 2717) (Figure 3C).

We therefore chose as our final controller the cAR controller in which *P_7tet.1_* directed the expression of both TetR and of the controlled gene, while *P_RNR2_* directed the synthesis of TetR-nls-Tup1. We constructed plasmids such that the tetR and tetR-nls-tup1 components are encoded on a single integrative plasmid, and a separate plasmid can be used to generate PCR fragments bearing *P_7tet.1_* for homologous recombination directed replacement of the promoter of any yeast gene. Due to its ability to give precisely regulated expression over a wide range of inducer concentrations, we called this cAR construct a “Well Tempered Controller” and gave it the number of Bach’s first Prelude and Fugue^1^(Figure 4A).

**Figure 4:**
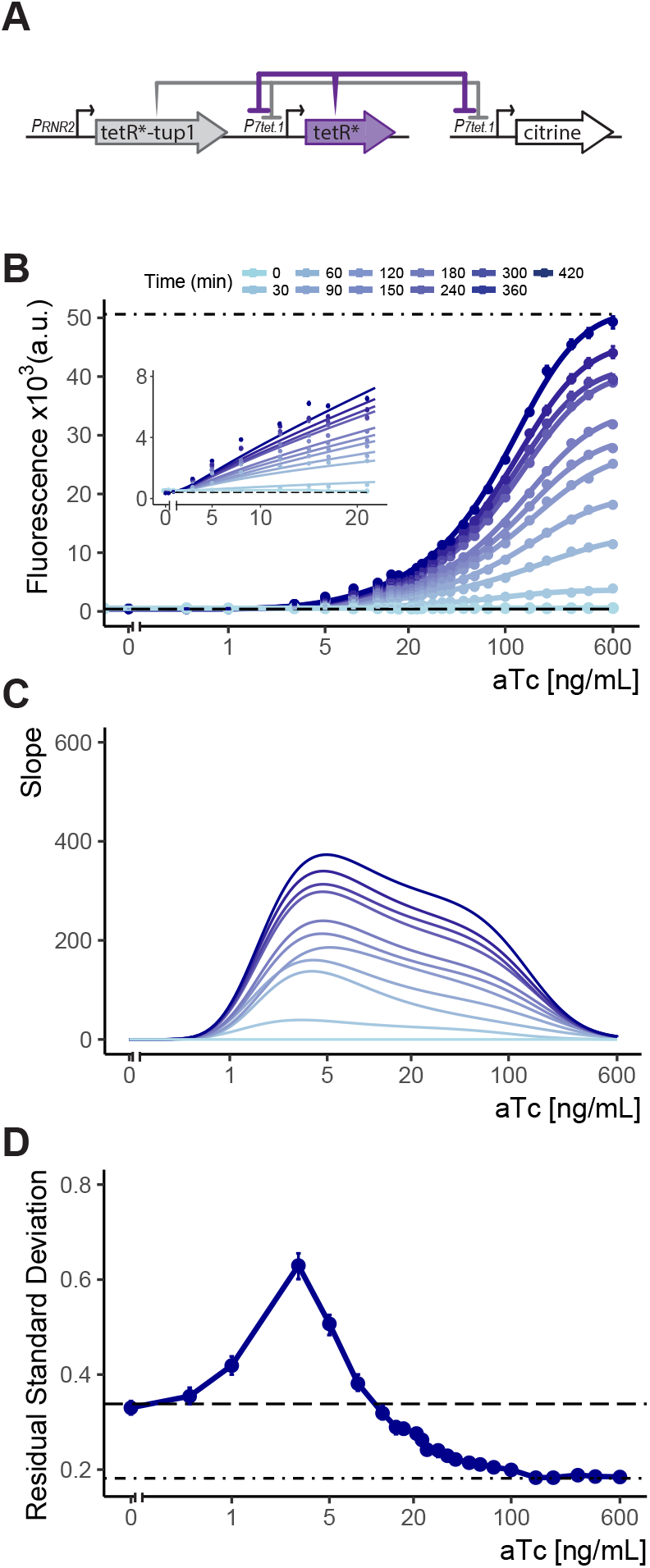
Controlled gene expression from WTC_846_. **A) Architecture of WTC_846_**. The final WTC_846_ system is composed of a single integrative plasmid bearing TetR and TetR-Tup1 driven by the promoters indicated. This plasmid was integrated at the URA3 locus. *P_7tet.1_* driven Citrine was integrated at the LEU2 locus. * indicates SV40 Nuclear Localization Sequence. Repression of promoters is indicated by flat headed arrows. **B) Time dependent dose response of WTC_846_-controlled expression**. Citrine fluorescence was measured using flow cytometry at 30 min intervals after induction with different concentrations of aTc (ng/mL). Dashed line indicates median autofluorescence (parent strain without Citrine) and dot dashed line fluorescent signal from wild type *P_TDH3_* (FRY2683). Circles show the median of the experimentally measured population, and the lines were fitted as explained in Figure 2B. The inset shows response at low input aTc doses. **C) The slopes of the dose response curves in (A)**, as a visual representation of the input dynamic range, defined as the range of doses where the slope of the dose response curve is non-zero. **D) Cell-to-cell variation of WTC_846_-controlled expression**. Single reporter CCV at 7h calculated as in 2D. Dashed line shows CCV of autofluorescence, dot-dashed line CCV of *P_TDH3_*-driven Citrine signal. Where present, error bars indicate 95% confidence interval calculated using bootstrapping (n=1000) as described in Materials&Methods.

### Well-tempered Controller_846_ (WTC_846_) fulfills the criteria of an ideal transcriptional controller

We measured the time dependent dose response of fluorescent signal in FRY2759, the *WTC_846_::Citrine* strain during exponential growth using flow cytometry (Figure 4B&C). Without aTc, there was no signal above background. After induction, signal appeared within 30 minutes, and reached steady state within 7 hours. Steady state expression was adjustable over aTc concentrations from 0.5ng/mL to 600ng/mL, a 1200-fold input dynamic range. Maximum expression was similar to that for the *P_TDH3_*-Citrine strain FRY2683. Direct observation of Citrine and TetR expression by Western blotting showed no expression of Citrine in absence of aTc, adjustable Citrine levels over the same input dynamic range and TetR expression synchronized with Citrine (Figure S9). In all 8 growth media tested, *WTC_846_::Citrine* expression in FRY2759 was precisely adjustable (Figure S10), and even very high induction of the WTC_846_ system in a strain where only the control plasmid bearing tetR and tetR-nls-tup1 was integrated (FRY2761) had no significant effect on growth rates (Figure S11).

We quantified the cell-to-cell variability in Citrine expression in the *WTC_846_::Citrine* strain (FRY2759) grown in YPD (Figure 4D, Figure S12&S14). At increasing concentrations of aTc, CCV initially rose to 0.63 at 8ng/mL, similar to the CCV measured for Citrine expression repressed by *P_RNR2_*-driven TetR-nls-Tup1 in an SR strain (FRY2717, RSD of 0.67, Figure S13). At higher aTc inputs, CCV rapidly dropped below that seen in FRY70, an otherwise-isogenic autofluorescence control strain, and reached the same low level (0.18) observed for Citrine whose expression was driven by *P_TDH3_* (FRY2683). Because the autofluorescence varied so greatly, absolute CCV for cells grown in different media could not be directly compared. However, under all growth conditions (Figure S15), CCV was highest at the similarly low concentrations of aTc and decreased at higher concentrations to the levels shown by the *P_TDH3_*-Citrine strain (Figure S10). We interpret the peak of CCV in the input dynamic range as arising from the fact that the cAR architecture combines Simple Repression and Autorepression of the *P_7tet.1_*-controlled gene (here, Citrine). At low concentrations of inducer, in the SR regime, most repression of *P_7tet.1_* was due to the constitutively expressed TetR-nls-Tup1, and the peak CCV was similar to that found for the strain where *P_7tet.1_* was repressed by constitutively expressed TetR-nls-Tup1 (Figure S13 and see previous Results section). At higher concentrations of aTc, in the AR regime, *P_7tet.1_* is derepressed, the concentration of TetR and the ratio of TetR to TetR-nls-Tup1 is large. At these inducer concentrations, TetR controls its own synthesis and variability is suppressed by this negative feedback. Taken together, these results indicated that WTC_846_ fulfilled our initially stated criteria for an ideal conditional expression system.

### WTC_846_ alleles allow precise control over protein dosage and cellular physiology

We then assessed the ability of WTC_846_ to direct conditional expression of endogenous genes. We selected (i) genes that are essential for growth, but for which previously generated transcriptionally controlled alleles still formed colonies on solid medium (*CDC42*, *TOR2*, *PBR1*, *CDC20*) or continued to grow in liquid medium (*PMA1*) under uninduced conditions, (ii) essential genes for which existing transcriptionally controlled alleles did not show the expected overexpression phenotype (*IPL1*), or (iii) essential genes for which conditional expression alleles did not exist (*CDC28*).^50–52^ These genes encoded proteins with a variety of functions: stable (Cdc28) and unstable (Cdc20 and Cdc42) cell cycle regulators, a spindle assembly checkpoint kinase (Ipl1), a metabolic regulator (Tor2), a putative oxidoreducatase (Pbr1), and a high abundance membrane proton pump (Pma1). The encoded proteins spanned a range of abundance from ~1000 (Tor2 and Ipl1) to ¿50,000 (Pma1) molecules per cell.^33^

We constructed strains in which WTC_846_ controlled the expression of these genes (Figure 5A and Table 1, strains labelled *WTC_846-Kx_::gene_name)*. To make these, we integrated a single plasmid-borne TetR-nls-Tup1 and autorepressing TetR construct into the LEU2 locus in a BY4741 background, and replaced sequences upstream of the ATG of the essential gene with a ~1940bp casette carrying an antibiotic selection marker and *P_7tet.1_*, without altering the sequence of the upstream gene or its terminator. In most cases we removed between 20-200bp of the endogenous gene promoter. The cassette carried one of three different 15bp translation initiation sequences (extended Kozak sequences; K1, K2, K3) as the last 15 bases before the ATG. These were designed according to,^53^ to enable different levels of translation of the gene’s mRNA. The predicted efficiency of the sequences was K1¿ K2¿ K3. If cells of a strain carrying a WTC_846_-controlled essential gene formed colonies on solid medium without aTc, we constructed an otherwise-isogenic strain with a lower efficiency Kozak sequence (data not shown).

**Figure 5:**
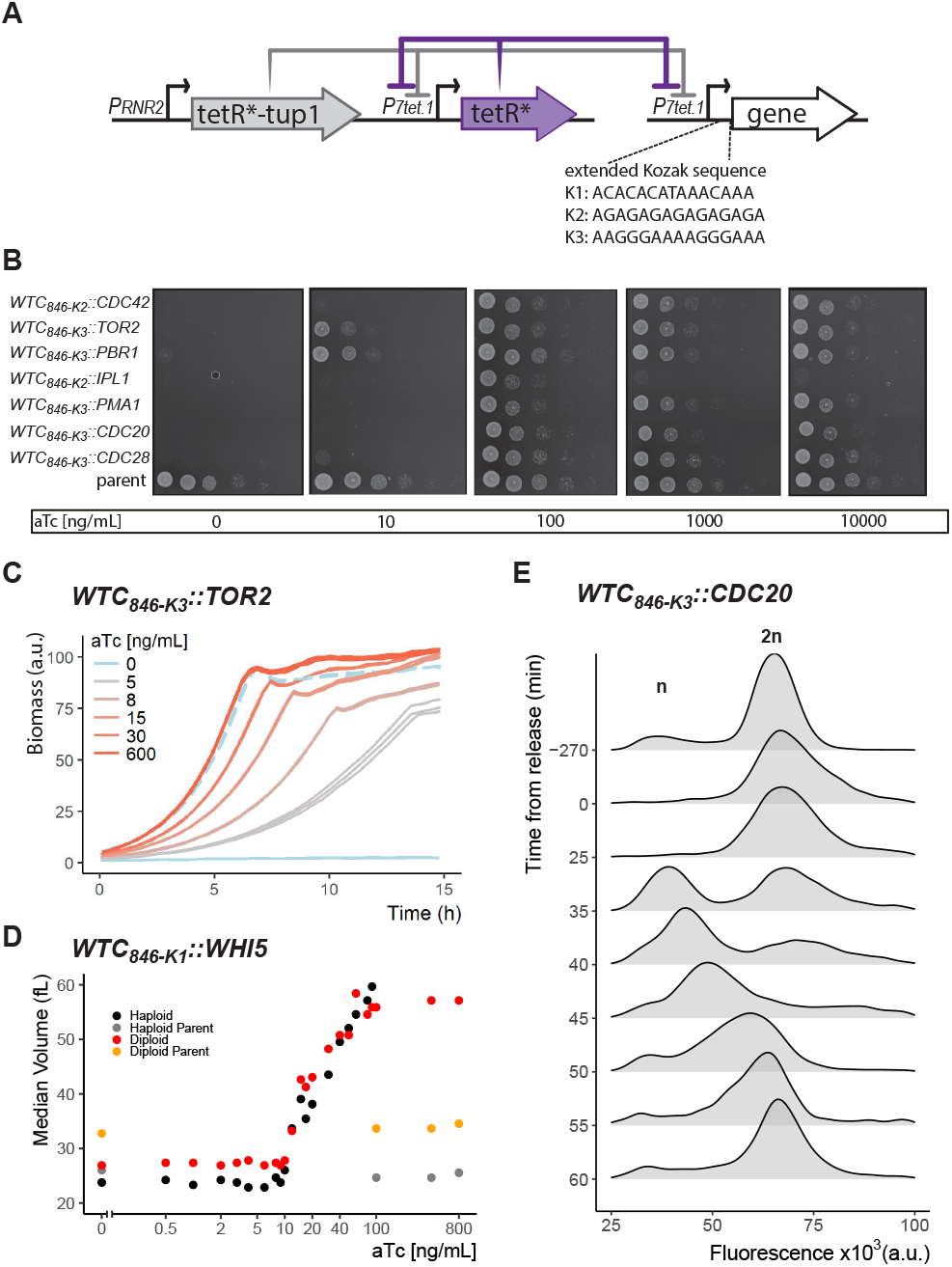
Controlled protein dosage of WTC_846_-driven yeast genes. **A) The WTC_846_ architecture used**, as in Figure 4A. Figure also shows the 3 extended Kozak sequences used to control translation efficiency. **B) WTC_846_ alleles of essential yeast genes show null and quantitative expression phenotypes**. The genes whose expression is controlled by WTC_846_ are indicated on the left. Cells growing in liquid medium were spotted onto different YPD plates, such that the leftmost circle on each plate had 2.25×10^6^ cells and each subsequent column is a 1:10 dilution. aTc concentration in each plate is indicated below each image. Parent refers to the strain where all components of WTC_846_ except the *P_7tet.1_* that directs expression of the controlled gene was present (FRY2769). **C) Precise control of growth rate by adjusting Tor2 protein dosage**. Growth of the *WTC_846-K3_::TOR2* strain was measured by scattered light intensity using a growth reader. Cells were grown in liquid YPD, three replicate wells per aTc concentration were measured. Dashed line indicates the growth curve of the parent strain, where Tor2 was under endogenous control. The y-axis was normalized to a range between 0-100 and indicates culture density. **D) Precise control of cell volume by titrating dosage of Whi5**. Haploid and Diploid refer to *WTC_846-K1_::WHI5* alleles grown in S Ethanol with varying concentrations of aTc. Haploid and diploid parent indicates strains where Whi5 was under endogenous control. Median cell volume was measured using a Coulter Counter. **E) Batch culture cell cycle synchronization**. A batch culture of *WTC_846-K3_::CDC20* strain growing in 20ng/mL aTc was arrested and synchronized by aTc withdrawal. Cells were released from the cell cycle block by addition of aTc at time 0. Cells were stained with Sytox and analyzed with flow cytometry. 10000 cells per time point were recorded. The plots are density distributions of the Sytox fluorescent signal of the whole population, such that the area under the curve equals 1. The peaks corresponding to one and 2 sets of chromosomes are labeled. These indicate the cells that are in G1 and G2/M phases of the cell cycle, respectively.

**Table 1:** Main strains used in this work and their relevant genotype. A detailed table including all strains used in the supplementary figures can be found in the Supplement.

| FRY | Name | Relevant Genotype |
| --- | --- | --- |
| 70 | autofluorescence | BY4743 derivative, haploid, MATa *his3Δ1 leu2Δ0 met15Δ0 ura3Δ0 lys2Δ0* |
| 2683 | *P<sub>TDH3</sub>*-const | *leu2Δ::P<sub>TDH3</sub>-citrine-LEU2* |
| 2551 | *P<sub>2tet</sub>*-const | *leu2Δ::P<sub>2tet</sub>-citrine-LEU2* |
| 2564 | *P<sub>3tet</sub>*-const | *leu2Δ::P<sub>3tet</sub>-citrine-LEU2* |
| 2566 | *P<sub>5tet</sub>*-const | *leu2Δ::P<sub>5tet</sub>-citrine-LEU2* |
| 2562 | *P<sub>2tet</sub>*-SR | *leu2Δ::P<sub>2tet</sub>-citrine-LEU2 his3Δ::P<sub>ACT1</sub>-tetR-NLS-HIS3* |
| 2573 | *P<sub>3tet</sub>*-SR | *leu2Δ::P<sub>3tet</sub>-citrine-LEU2 his3Δ::P<sub>ACT1</sub>-tetR-NLS-HIS3* |
| 2577 | *P<sub>5tet</sub>*-SR | *leu2Δ::P<sub>5tet</sub>-citrine-LEU2 his3Δ::P<sub>ACT1</sub>-tetR-NLS-HIS3* |
| 2659 | *P<sub>5tet.1</sub>*-const | *leu2Δ::P<sub>5tet.1</sub>-citrine-LEU2* |
| 2656 | *P<sub>5tet.1</sub>*-SR | *leu2Δ::P<sub>5tet.1</sub>-citrine-LEU2 his3Δ::P<sub>ACT1</sub>-tetR-NLS-HIS3* |
| 2661 | *P<sub>7tet.1</sub>*-const | *leu2Δ::P<sub>7tet.1</sub>-citrine-LEU2* |
| 2663 | *P<sub>7tet.1</sub>*-SR | *leu2Δ::P<sub>7tet.1</sub>-citrine-LEU2 his3Δ::P<sub>ACT1</sub>-tetR-NLS-HIS3* |
| 2674 | *P<sub>7tet.1</sub>*-AR | *leu2Δ::P<sub>7tet.1</sub>-citrine-LEU2 met15Δ::P<sub>7tet.1</sub>-tetR-NLS-MET15* |
| 2741 | *P<sub>7tet.1</sub>*-cAR | *leu2Δ::P<sub>7tet.1</sub>-citrine-LEU2 met15Δ::P<sub>7tet.1</sub>-tetR-NLS-MET15 his3Δ::P<sub>ACT1</sub>-tetR-NLS-HIS3* |
| 2673 | *P<sub>7tet.1</sub>*-cAR(*P<sub>ACT1</sub>*-TUP1) | *leu2Δ::P<sub>7tet.1</sub>-citrine-LEU2 his3Δ::P<sub>ACT1</sub>-tetR-NLS-tup1-HIS3 met15Δ::P<sub>7tet.1</sub>-tetR-NLS-MET15* |
| 2684 | *P<sub>7tet.1</sub>*-cAR(*P<sub>VPH1</sub>*-TUP1) | *leu2Δ::P<sub>7tet.1</sub>-citrine-LEU2 his3Δ::P<sub>VPH1</sub>-tetR-NLS-tup1-HIS3 met15Δ::P<sub>7tet.1</sub>-tetR-NLS-MET15* |
| 2749 | *P<sub>7tet.1</sub>*-cAR(*P<sub>RNR2</sub>*-TUP1) | *leu2Δ::P<sub>7tet.1</sub>-citrine-LEU2 his3Δ::P<sub>RNR2</sub>-tetR-NLS-tup1-HIS3 met15Δ::P<sub>7tet.1</sub>-tetR-NLS-MET15* |
| 2715 | *P<sub>7tet.1</sub>*-cAR(*P<sub>PREV1</sub>*-TUP1) | *leu2Δ::P<sub>7tet.1</sub>-citrine-LEU2 his3Δ::P<sub>PREV1</sub>-tetR-NLS-tup1-HIS3 met15Δ::P<sub>7tet.1</sub>-tetR-NLS-MET15* |
| 2669 | *P<sub>7tet.1</sub>*-SR(*P<sub>ACT1</sub>*-TUP1) | *leu2Δ::P<sub>7tet.1</sub>-citrine-LEU2 his3Δ::P<sub>ACT1</sub>-tetR-NLS-tup1-HIS3* |
| 2676 | *P<sub>7tet.1</sub>*-SR(*P<sub>VPH1</sub>*-TUP1) | *leu2Δ::P<sub>7tet.1</sub>-citrine-LEU2 his3Δ::P<sub>VPH1</sub>-tetR-NLS-tup1-HIS3* |
| 2717 | *P<sub>7tet.1</sub>*-SR(*P<sub>RNR2</sub>*-TUP1) | *leu2Δ::P<sub>7tet.1</sub>-citrine-LEU2 his3Δ::P<sub>RNR2</sub>-tetR-NLS-tup1-HIS3* |
| 2759 | *WTC<sub>846</sub>::Citrine* | *leu2Δ::P<sub>7tet.1</sub>-citrine-LEU2 ura3Δ::P<sub>RNR2</sub>-tetR-NLS-tup1-*P<sub>7tet.1</sub>*-tetR-NLS-URA3* |
| 2761 | *WTC<sub>846</sub>::* | *ura3Δ::P<sub>RNR2</sub>-tetR-NLS-tup1-*P<sub>7tet.1</sub>*-tetR-NLS-URA3* |
| 2769 | parent | *whi5Δ::WHI5-mKOkappa-HIS3, myo1Δ::MYO1-mKate(3x)-KanMX, leu2Δ::P<sub>RNR2</sub>-tetR-NLS-tup1-*P<sub>7tet.1</sub>*-tetR-NLS-LEU2* |
| 2772 | *WTC<sub>846-K1</sub>::TOR2* | *whi5Δ::WHI5-mKOkappa-HIS3, myo1Δ::MYO1-mKate(3x)-KanMX, leu2Δ::P<sub>RNR2</sub>-tetR-NLS-tup1-*P<sub>7tet.1</sub>*-tetR-NLS-LEU2 P<sub>TOR2</sub>::P<sub>7tet.1</sub>-K1-HygMX* |
| 2775 | *WTC<sub>846-K2</sub>::CDC28* | *whi5Δ::WHI5-mKOkappa-HIS3, myo1Δ::MYO1-mKate(3x)-KanMX, leu2Δ::P<sub>RNR2</sub>-tetR-NLS-tup1-*P<sub>7tet.1</sub>*-tetR-NLS-LEU2 P<sub>CDC28</sub>::P<sub>7tet.1</sub>-K2-NatMX* |
| 2837 | *WTC<sub>846-K3</sub>::CDC20* | *whi5Δ::WHI5-mKOkappa-HIS3, myo1Δ::MYO1-mKate(3x)-KanMX, leu2Δ::P<sub>RNR2</sub>-tetR-NLS-tup1-*P<sub>7tet.1</sub>*-tetR-NLS-LEU2 P<sub>CDC20</sub>::P<sub>7tet.1</sub>-K3-NatMX* |
| 2788 | *WTC<sub>846-K2</sub>::CDC42* | *whi5Δ::WHI5-mKOkappa-HIS3, myo1Δ::MYO1-mKate(3x)-KanMX, leu2Δ::P<sub>RNR2</sub>-tetR-NLS-tup1-*P<sub>7tet.1</sub>*-tetR-NLS-LEU2 P<sub>CDC42</sub>::P<sub>7tet.1</sub>-K2-NatMX* |
| 2789 | *WTC<sub>846-K2</sub>::IPL1* | *whi5Δ::WHI5-mKOkappa-HIS3, myo1Δ::MYO1-mKate(3x)-KanMX, leu2Δ::P<sub>RNR2</sub>-tetR-NLS-tup1-*P<sub>7tet.1</sub>*-tetR-NLS-LEU2 P<sub>IPL1</sub>::P<sub>7tet.1</sub>-K2-NatMX* |
| 2828 | *WTC<sub>846-K3</sub>::PMA1* | *whi5Δ::WHI5-mKOkappa-HIS3, myo1Δ::MYO1-mKate(3x)-KanMX, leu2Δ::P<sub>RNR2</sub>-tetR-NLS-tup1-*P<sub>7tet.1</sub>*-tetR-NLS-LEU2 P<sub>PMA1</sub>::P<sub>7tet.1</sub>-K3-NatMX* |
| 2773 | *WTC<sub>846-K3</sub>::TOR2* | *whi5Δ::WHI5-mKOkappa-HIS3, myo1Δ::MYO1-mKate(3x)-KanMX, leu2Δ::P<sub>RNR2</sub>-tetR-NLS-tup1-*P<sub>7tet.1</sub>*-tetR-NLS-LEU2 P<sub>TOR2</sub>::P<sub>7tet.1</sub>-K3-HygMX* |
| 2827 | *WTC<sub>846-K3</sub>::CDC28* | *whi5Δ::WHI5-mKOkappa-HIS3, myo1Δ::MYO1-mKate(3x)-KanMX, leu2Δ::P<sub>RNR2</sub>-tetR-NLS-tup1-*P<sub>7tet.1</sub>*-tetR-NLS-LEU2 P<sub>CDC28</sub>::P<sub>7tet.1</sub>-K3-NatMX* |
| 2830 | *WTC<sub>846-K3</sub>::PBR1* | *whi5Δ::WHI5-mKOkappa-HIS3, myo1Δ::MYO1-mKate(3x)-KanMX, leu2Δ::P<sub>RNR2</sub>-tetR-NLS-tup1-*P<sub>7tet.1</sub>*-tetR-NLS-LEU2 P<sub>PBR1</sub>::P<sub>7tet.1</sub>-K3-NatMX* |
| 2849 | *WTC<sub>846-K3</sub>::TPI1* | *leu2Δ::P<sub>RNR2</sub>-tetR-NLS-tup1-*P<sub>7tet.1</sub>*-tetR-NLS-LEU2 P<sub>TPI1</sub>::P<sub>7tet.1</sub>-K3-NatMX* |
| 2791 | *WTC<sub>846-K1</sub>::WHI5* | *whi5Δ::WHI5-mKOkappa-HIS3, myo1Δ::MYO1-mKate(3x)-KanMX, leu2Δ::P<sub>RNR2</sub>-tetR-NLS-tup1-*P<sub>7tet.1</sub>*-tetR-NLS-LEU2 P<sub>WHI5</sub>::P<sub>7tet.1</sub>-K1-NatMX* |
| 2929 | *WTC<sub>846-K1</sub>::WHI5(diploid)* | BY4743, *whi5Δ::WHI5-mKOkappa-HIS3/WHI5 myo1Δ::MYO1-mKate(3x)-KanMX/MYO1 leu2Δ::P<sub>RNR2</sub>-tetR-NLS-tup1-*P<sub>7tet.1</sub>*-tetR-NLS-LEU2/leu2Δ0 ura3Δ::P<sub>RNR2</sub>-tetR-NLS-tup1-*P<sub>7tet.1</sub>*-tetR-NLS-URA3/ura3Δ0 P<sub>WHI5</sub>::P<sub>7tet.1</sub>-K1-HygMX/P<sub>WHI5</sub>::P<sub>7tet.1</sub>-K1-NatMX* |

We spotted serial dilutions of cultures of the final seven strains on YPD, YPE, SD, S Glycerol and SD Proline plates with and without inducer (Figure 5B and Figure S16). On all these media, no strain formed colonies without aTc and at intermediate concentrations of aTc all strains did. This result showed that WTC_846_ alleles can produce null phenotypes.

The *WTC_846-K2_::IPL1* strain formed colonies with lower plating efficiency than the parent strain at a high aTc concentration. Ipl1 is a component of the kinetochore, required for correct sister chromatid separation during mitosis. In mouse embryonic fibroblasts, overexpression of the orthologous Aurora B kinase causes aberrant chromosome segregation and increases duration of mitosis by activating the Spindle Assembly Checkpoint, which stops mitosis until correct spindle attachments to sister chromatids can be formed.^54^ In *S. cervisiae*, however, *P_GAL1-10_*-driven overexpression of Ipl1 does not decrease plating efficiency, does not cause accumulation of cells with 2n DNA content unable to complete mitosis, and does not cause aberrant chromosome segregation as assessed by microscopy, unless simultaneously overexpressed with another kinetochore component (Sli15).^55^ We asked whether WTC_846_-driven Ipl1 overexpression alone could show these phenotypes in *S. cerevisiae*. We cultured *WTC_846-K2_::IPL1* cells for 18 hours in YPD with a high concentration of aTc (400ng/mL), and measured total DNA content in flow cytometry to assess cell cycle progression. In these cultures compared to the parent with WT Ipl1, many cells were in the G2/M phase with 2n DNA content, indicative of an inability to complete mitosis, and a significant portion of the population showed aberrant chromosome numbers above 2n (Figure S17). Thus, the phenotype caused by WTC_846_-driven Ipl1 overexpression in *S. cerevisiae* resembles that of Aurora B overexpression in mammalian cells.

We tested whether adjustable expression of metabolic and essential genes could be used to titrate growth rates. We constructed strains with WTC_846_ alleles of Tor2, a low abundance, stable, essential protein necessary for nutrient signalling and actin polarization,^56^ Pma1, an abundant, essential proton pump that regulates the internal pH of the cell,^57^ and Tpi1, a highly abundant, non-essential glycolytic enzyme^58^ (FRY2773, 2828, 2849). We cultured *WTC_846_::TOR2*, *WTC_846_::PMA1*, and *WTC_846_::TPI1* cells in different liquid media over a large input dynamic range of aTc, and measured growth by scattered light intensity in a growth reader as a proxy for culture density (Biolector or GrowthProfiler) (Figure 5C for Tor2 and Figure S18 for all three proteins). All strains showed distinct growth rates at different aTc concentrations. For all strains, we identified an aTc concentration that resulted in the same growth rate of as the otherwise-isogenic strain bearing the native gene promoter. In order to assess whether the *WTC_846_::PMA1* strain showed the expected hypomorphic phenotype of defective daughter cell separation,^59^ we used flow cytometry and Sytox Green staining to quantify DNA content. At low aTc concentrations, cells showed an apparent increase in ploidy and cell size and microscopic observation showed that each mother had multiple daughters attached to it (Figure S19). Observation of *WTC_846_::TOR2* strains revealed a novel overexpression phenotype: at high aTc concentrations, cells bearing the higher translational efficiency *TOR2* allele (*WTC_846-K1_::TOR2*) grew more slowly than the otherwise-isogenic control parent strain with WT *TOR2* (Figure S18C). The strain with the less efficient *WTC_846-K3_::TOR2* allele did not show this overexpression phenotype. These results demonstrate that researchers can adjust input to WTC_846_ alleles to tune protein levels and different growth rates precisely, and that the dynamic range of phenotypic outputs can be further expanded by the ability to construct WTC_846_ alleles with alternative Kozak sequences to observe phenotypes at the two dosage extremes.

We then tested whether adjustable gene expression could precisely regulate cell size. In *S. cerevisiae*, Whi5 regulates the volume at which unbudded cells commit to a round of division and start forming buds. Δwhi5 cells are smaller, and cells expressing Whi5 under *P_GAL1_* control are larger than otherwise-isogenic cells.^60^ Whi5 controls cell volume by a complex mechanism. Whi5 mRNA and protein are expressed during S/G2/M (in haploids, at about 2500 molecules), and the Whi5 protein in the nucleus suppresses transcription of the G1 cyclins needed to commence a new round of division.^60, 61^ During G1, as cells increase in volume, the nuclear concentration of Whi5 falls due to dilution^62^ and slow nuclear export^63^ until a threshold is reached, after which Whi5 is rapidly exported from the nucleus, and cells enter START. Here, we constructed haploid and diploid *WTC_846_::WHI5* strains (FRY2791, FRY2929) in which expression of Whi5 mRNA was independent of cell cycle state. We grew these along with otherwise isogenic control strains in S Ethanol to exponential phase at different aTc concentrations, and measured cell volume using a Coulter counter. Increasing Whi5 expression resulted in increasingly larger cells (Figure 5D). Without aTc, diploid *WTC_846_::WHI5* cells were about the same volume as haploid controls (median 27fL vs 25fL), whereas haploid *WTC_846_::WHI5* cells were only slightly smaller at 24fL. At around 10 and 12 ng/mL aTc, both haploid and diploid strains had about the same volume as controls. At full induction, both *WTC_846_::WHI5* strains had a median volume of around 60fL, almost twice as large as the diploid control, yielding a more than 2-fold range of possible cell volumes attainable using WTC_846_ for both haploid and diploid cells. We also calculated the CoV of cell volume to assess cell-to-cell variation of this WTC_846_ directed phenotype. For most of the volume range, the CoV was around the same level as for the control strains with WT Whi5 (Figure S20). Cells expressing high levels of Whi5 especially in haploid cells showed increased variation in volume, perhaps a consequence of more uniform transcription of WTC_846_-driven, overexpressed Whi5 throughout the cell cycle.

Finally, we tested the ability of WTC_846_ to exert dynamic control of gene expression by constructing a *WTC_846-K3_::CDC20* strain (FRY2837) and using this allele to synchronize cells in batch culture by setting Cdc20 expression to zero and then restoring it.^64^ Cdc20 is an essential activator of Anaphase Promoting Complex C, which once bound to Cdc20, initiates the mitotic metaphase to anaphase transition,^65^ and is then degraded during anaphase. Upon depletion of Cdc20, for example by shift of *ts* strains to the restrictive temperature, or transcriptionally controlled alleles to non-inducing medium, cells arrest in metaphase with large buds and 2n DNA content. When Cdc20 is restored by switching to the permissive condition, cells enter the next cell cycle simultaneously.^66, 67^ Use of *WTC_846_::CDC20* to synchronize the cells in a culture would require complete transcriptional shut-off, followed by rapid and complete re-expression of Cdc20 in all the cells in a population. We diluted exponentially growing *WTC_846-K3_::CDC20* cells into YPD medium without aTc (0.5 million cells/mL) and took samples for Sytox staining and flow cytometry analysis for DNA content at fixed intervals. Within 480 minutes, the entire culture arrested at the G2/M phase with 2n DNA content (Figure 5E). We used microscopic inspection to verify that cells had arrested with large buds, as is expected upon a G2/M arrest. We then added 600ng/mL of aTc. As assayed by Sytox staining and flow cytometry and confirmed by microscopy, cells then re-entered the cell cycle within 35 minutes and went through one cell cycle completely synchronously.

## Discussion

Conditional expression of genes into phenotypes remains important for biological discovery. Many methods used historically, such as suppression of nonsense mutations, or conditional inactivation of temperature sensitive mutations, do not facilitate titration of graded or intermediate doses of protein. More current methods for graded expression do not allow experimenters to adjust and set protein levels and show high cell-to-cell variation of protein expression in cell populations, limiting their utility for elucidating protein-dosage dependent phenotypes. Moreover, most such methods also have secondary consequences including slowing of cell growth. In order to overcome these limitations, we developed for use in *S. cerevisiae* a “Well-tempered Controller”. This controller, WTC_846_, is an autorepression-based transcriptional controller of gene expression. It can set protein levels across a large input and output dynamic range. WTC_846_ alleles display no uninduced basal expression, high maximum expression, low cell-to-cell variation, and operate in different media conditions without adverse effects on cell physiology.

The central component of WTC_846_ is an engineered TATA containing promoter, *P_7tet.1_*. We and others had shown that prokaryotic repressors including LexA,^23^ TetR,^26^ λcI^27^ and LacI^24, 68^ can block transcription from engineered TATA-containing eukaryotic promoters, when those promoters contain binding sites between the Upstream Activating Region (UAS) (or, for vertebrate cells, the enhancer) and the TATA^23^ or downstream of or flanking the TATA. To develop *P_7tet.1_*, we placed seven *tetO_1_* TetR binding sites in the promoter of the strongly expressed yeast gene *TDH3*. Two of the sites flank the TATA sequence, the other five abut binding sites for an engineered UAS that binds the transcription activators Rap1 and Gcr1. In WTC_846_, one instance of *P_7tet.1_* drives expression of the controlled gene, while a second instance of *P_7tet.1_* drives expression of the TetR repressor, which thus represses its own synthesis.

We believe that repression of *P_7tet.1_* by TetR is due mainly to its action at the two *tetO_1_* sites flanking the TATA sequence, because TetR represses a precursor promoter that only carries such sites to the same extent. The mechanism(s) by which binding of repressors near the TATA might interfere with Pre-Initiation Complex (PIC) formation, transcription initiation, or early elongation remain unknown, as well as why binding of larger presumably transcriptionally inert TetR fusion proteins results in stronger repression. However, examination of the Cryo-EM structure of TBP and TFIID bound to mammalian TATA promoters^69^ suggests that binding of TetR and larger derivatives of it to these sites might simply block PIC assembly. Studies of repression of native *Drosophila melanogaster* promoters by the *en* and *eve* homeobox proteins show that a similar, steric occlusion based mechanism can block eukaryotic transcription by binding of the repressors to sites close to the TATA sequence.^70, 71^

In WTC_846_, when inducer is absent, basal expression of the controlled gene is abolished by a second TetR derivative, a fusion bearing an active repressor protein native to yeast. Because the same TetR-nls-Tup1 fusion protein fully represses a precursor promoter that only carries TetR operators in the UAS, we believe that the main zeroing effect of TetR-nls-Tup1 is manifested through binding the *tetO_1_* sites in the UAS. Native Tup1 repressor complexes with Ssn6 (also called Cyc8) to form a 420kDa protein complex,^42^ and TetR binds DNA as a dimer. In Δgcr1 cells, in which transcription from *P_TDH3_* is severely diminished, the native *P_TDH3_* promoter has two nucleosomes positioned between the UAS and the transcription start site.^72^ It is thus possible that in the UAS as many as five very large dimeric TetR-nls-Tup1 complexes block binding of Gcr1 and Rap1, mask their activating domains, or some combination of these, resulting in similar placement of two nucleosomes in *P_7tet.1_*. One of these nucleosomes could then be positioned at the 294nt stretch between the UAS and the TATA sequence. It also seems possible that binding of the Tet-Rnls-Tup1 repressor might shift the position of the second nucleosome further downstream, so that it obscures the transcription start site.

Both the increased input dynamic range and the lower cell-to-cell variation in expression from WTC_846_ arise from the fact that the TetR protein that represses the controlled gene also represses its own synthesis. This autorepression architecture is common in prokaryotic regulons^73, 74^ including Tn10, the source of the TetR gene used here, and it has been engineered into eukaryotic systems.^38, 41^ In self-repressing TetR systems, the input (here, anhydrotetracycline (aTc)) and TetR output together function as a comparator-adjustor.^75^ In such systems, aTc diffuses into the cell. Intracellular aTc concentration is limited by entry. Inside the cell, aTc and TetR^free^ concentrations are continuously compared by their binding interaction. If TetR^free^ is in excess, it represses TetR expression, and total intracellular TetR concentration is reduced by dilution, cell division, and active degradation of DNA-bound TetR until an equilibrium determined by the intracellular aTc concentration is again reached. The consequence of this autorepression is that the WTC_846_ requires more aTc to reach a given level of controlled gene expression than strains in which TetR is expressed only constitutively. Autorepression flattens the dose response curve, increases the range of aTc doses where a change in promoter activity can be observed, and buffers the effects of stochastic cell-to-cell variations in TetR concentration, thereby reducing cell-to-cell variation (CCV) in expression of the controlled gene throughout the input dynamic range.

We further extended the output dynamic range of WTC_846_-controlled genes by developing three different Kozak sequences, K1, K2, and K3,^53^ to allow controlled genes to be translated at different levels. We used these sequences to construct strains bearing conditional alleles of the essential genes *CDC28*, *IPL1*, *TOR2*, *CDC20*, *CDC42*, *PMA1*, and *PBR1*. These strains all showed graded expression of growth and other phenotypes, from lethality at zero expression to penetrant expression of previously observed phenotypes at higher protein dosage. We used controlled expression in the *WTC_846_::CDC20* strain to bring about G2/M arrest followed by synchronous release, with fast induction kinetics and low cell-to-cell variation in induction timing, thus demonstrating that WTC_846_ can be used in experimental approaches that require dynamic control of gene expression. We showed in *WTC_846_::IPL1* strains that high level expression of this spindle assembly checkpoint kinase arrests cells at G2/M with 2n or higher DNA content. This phenotype, thought to be due to disruption of kinetichore microtubule attachments, is displayed in mammalian cells when the homologous Aurora B is overexpressed,^54^ but had not been observed previously in *S. cerevisiae* when Ipl1 was overexpressed from a *PGal1-10* fusion promoter.^55^ We also showed that in *WTC_846_::WHI5* strains, different levels of controlled expression of Whi5 can constrain cell sizes within different limits.

Cell-cell variation in WTC_846_-driven expression is highest at low aTc levels, because control in this regime depends mostly on the higher variability Simple Repression (SR) architecture. Notably, transcription of WTC_846_-controlled genes is synchronized to that of the autorepressing TetR protein, so that mRNA expression of WTC_846_-driven genes (such as *WHI5*) should be steady throughout the cell cycle (although protein abundance may not be). However, despite these sources of residual variation, this autorepressing circuitry operationally defines WTC_846_ as a first example of an “expression clamp”, a device for adjusting and setting gene expression at desired levels, and maintaining it with low cell-to-cell variation, and so allowing protein concentration/expressed protein dosage in individual cells to closely track the population average.

Taken together, our results show that WTC_846_-controlled genes define a new type of conditional allele that allows precise control of gene dosage. We anticipate that WTC_846_ alleles will find use in cell biological experimentation, for example in assessment of phenotypes now incompletely penetrant due to variable dosage of the causative gene products,^76^ and for sharpening the thresholds at which dosage dependent phenotypes manifest. We also expect them to find use in genome-wide gene-by-gene and gene-by-chemical epistasis screens to detect protein dosage independent interactions, and in engineering applications such affinity maturation of antibodies expressed by yeast surface display, where precise ability to lower surface concentration can aid selection for progressively higher affinity binders. Implementation of the WTC_846_ control logic in mammalian cells and in engineered multicellular organisms should allow similar experimentation now impossible due to cell-to-cell variation and imprecise control.

## Materials&Methods

### Plasmids

Information on plasmids, and promoter and protein sequences used in this study can be found in Tables S2 and S4. Plasmids with auxotrophic markers were constructed based on the pRG shuttle vector series^36^ using either restriction enzyme cloning or isothermal assembly.^77^ Inserts were generated either by PCR on existing plasmids or custom DNA synthesis (GeneArt, UK). Oligos for cloning and for strain construction were synthesized by Thermofisher, UK. Plasmids used to generate linear PCR products for tagging transformations were based on the pFA6 backbone.^78^ Plasmids necessary to construct WTC_846_ strains are available through Addgene. Plasmid structures and a detailed protocol for strain construction can be found in the Supplement.

pRG shuttle vector series backbones used for integrative transformations have T7 and T3 promoters flanking the insert.^36^ During cloning, the insert of plasmids bearing TetR were cloned such that the insert promoter was closer to the T7 promoter and the terminator was near the T3 promoter of the backbone. In plasmids bearing Citrine, the insert was flipped onto the opposite strand, such that the insert promoter was near the T3 promoter, and the terminator near the T7 promoter. This inversion was done to avoid homologous recombination during subsequent integration of these plasmids into the same strain, since in many strains TetR and Citrine were flanked by the same promoter and the same terminator.

### Strains

Strains used in this study can be found in Table S1. Strains used for fluorescent measurements and the *WTC_846-K3_::TPI1* strain are based on a BY4743 derivative haploid background (MATa Δ*his3* Δ*leu2* Δ*met15* Δ*ura3* Δ*lys2*). Strains where *P_7tet.1_* replaced endogenous promoters were based on the haploid BY4741 background with the modifications *whi5*Δ*::WHI5-mKOkappa-HIS3*, *myo1*Δ*::MYO1-mKate(3x)-KanMX* and so were resistant to kanamycin. The oligos used to replace the promoters of the different endogenous genes with WTC_846_-controlled *P_7tet.1_* can be found in Table S3. Correct replacement of the endogenous promoter with *P_7tet.1_* was checked using colony PCR with the protocol from the Blackburn lab (also detailed in^36^), and subsequent sequencing (Microsynth, Switzerland). For colony PCR we used a standard forward oligo annealing to *P_7tet.1_*, and gene specific reverse oligos annealing within the tagged gene. Oligo sequences for colony PCR can be found in Table S3.

### Chemicals & Media

YPD/YPE was prepared with 1% yeast extract (Thermofisher, 212720), 2% bacto-peptone (Thermofisher, 211820) and 2% glucose (Sigma, G8270) / ethanol (Honeywell, 02860). Synthetic (S) media except SD Proline contained 0.17% yeast nitrogen base (without amino acids and ammonium sulfate) (BD Difco, 233520) with 0.5% ammonium sulfate (Sigma, 31119) as nitrogen source, complete complement of amino acids and adenine and uracil, except for SD min which contained only the necessary amino acid complements to cover auxotrophies. SD Proline media contained 0.17% yeast nitrogen (without amino acids and ammonium sulfate), only the amino acids necessary to cover auxotrophies and 1mg/mL proline as the sole nitrogen source. The carbon source was 2% glucose for SD and SD Proline, 2% ethanol for S Ethanol, 3% glycerol for S Glycerol (Applichem, A2957), 2% fructose for S Fructose and 2% Raffinose for S Raffinose. Experiments were performed in YPD media unless otherwise specified. Solid medium plates were poured by adding 2% agar (BD Sciences, 214040) to the media described above.

aTc was purchased from Cayman Chemicals (10009542) and prepared as a 4628.8ng/mL (10mM) stock in ethanol for long term storage at −20°C and diluted in water for experiments as necessary.

When constructing strains where *P_7tet.1_* replaces endogenous promoters, a PCR fragment containing *P_7tet.1_* and an antibiotic marker (either Nourseothricin (Werner BioAgents, clonNAT) or Hygromycin (ThermoFisher,10687010)) was transformed for homologous recombination directed replacement of the endogenous promoter. Cells were plated on YPD + antibiotic plates for selection. Whenever the promoter of an essential gene was being replaced, transformations were plated on multiple plates with YPD + antibiotic and 10/50/100/500 ng/mL aTc.

### Spotting Assay

For spotting assays of cell growth and viability, cells were precultured in YPD media with 20ng/mL aTc (except for *WTC_846-K2_::IPL1* strain which was precultured in 10ng/mL aTc) and the necessary antibiotic to stationary phase, and diluted into YPD + antibiotic without aTc at a concentration of 0.8×10^6^ cells/mL. 6 hours later, cells were spun down and resuspended in YPD. Cells were spotted onto plates containing different media and aTc concentrations prepared as described above such that the most concentrated spot has 2.25×10^6^ cells, and each column is a 1:10 dilution. Pictures were taken after 24 hours for the YPD and SD plates, and 42 hours for SD Proline, S Glycerol and YPE plates.

### Flow Cytometry

Cells were diluted 1:200 from dense precultures and cultured to early exponential phase (2-5 × 10^6^ cells/mL) in 96 deep-well plates at 30°C before induction with aTc if necessary. For aTc dose responses, samples were taken at times indicated. For experiments where no dose response was necessary, cells were measured at least 4 hours after dilution of precultures, but always before stationary phase. Samples were diluted in PBS and measured using a LSR-Fortessa LSRII equipped with a high-throughput sampler. PMT voltages for the forward and side scatter measurements were set up such that the height of the signal was not saturated. Citrine fluorescence was quantified using a 488 nm excitation laser and a 530/30 nm emission filter. PMT voltage for this channel was set up such that the signal from *P_TDH3_* expressed Citrine did not saturate the measurement device, except for basal level measurements in Figure 3B and Figure S8, where PMT voltage for the Citrine channel was increased to maximum. Side scatter was measured using the 488 nm excitation laser and 488/10 nm emission filter.

### Western Blots

Cells were grown to stationary phase with the indicated aTc concentration. 5mL of cell culture was centrifuged and resuspended in 1mL 70% ethanol. Fixed cells were again centrifuged, and resuspended in 200uL Trupage LDS loading buffer (Merck, PCG3009) supplemented with 8M urea. Cells were broken using glass beads and a bead beater, and boiled at 95°C for 30 minutes. Proteins were separated using SDS-Page with Trupage precast 10% gels (Merck, PCG2009-10EA) and the associated commercial buffer, and transferred onto a nitrocellulose membrane (GE Healthcare Life Sciences, 10600008).

We used mouse monoclonal primary antibodies for detecting TetR (Takara, Clone 9G9), and Citrine (Merck, G6539), both diluted 1:2000 in Odyssey Blocking buffer (PBS) (LI-COR Biosciences) + 0.2% Tween 20. The secondary antibody was the nearinfrared fluorescent IRDye 800CW Goat anti-Mouse IgG Secondary Antibody from Li-Cor (926-32210), diluted 1:5000 in the same manner. We used Chameleon Duo pre-stained Protein Ladder as our molecular weight marker (928-60000). We used the SNAP i.d. 2.0 system which uses vacuum to drive reagents through the membrane, and the Odyssey CLx (LI-COR) detector for imaging. Images were processed using the Fiji software to obtain black and white images with high contrast.^79^

### Growth curves

Cells were precultured in YPD (with aTc in the case of strains where WTC_846_ controlled essential genes) to stationary phase, then diluted into fresh media at a concentration of 50.000 cells per mL and induced with the necessary aTc concentrations, except for YP Ethanol and S Ethanol media where the concentration was 500.000 cells per mL. The Growth Profiler 960 (EnzyScreen) with 96 well plates and 250uL volume per well, or Biolector (m2p-labs) with 48 well plates and 1mL volume per plate was used to measure growth curves. These are commercial devices that quantify culture density by detecting the light that is reflected back by the liquid culture.

### Arrest & Release Assay and DNA Staining

*WTC_846-K3_::CDC20* and the appropriate control strains were precultured in YPD (pH 4) with 20ng/mL aTc to a concentration of 2×10^6^ cells/mL, then centrifuged and diluted 1:3 into YPD (pH 4) without aTc. We found that low pH (pH4) of the media was necessary for efficient mother-daughter separation upon completion of cytokinesis, potentially due to the low pH optimum of the chitinase CTS1,^80^ which plays a role in separation. To prevent the culture from becoming too dense, 25% of the media was filtered and returned to the culture after 4 hours of growth without aTc, which removed 1/4th of the cells. After 8 hours of arrest, 600ng/mL aTc was added to the culture. Samples were taken every 90 minutes before, and every 5 minutes after aTc was added to the culture, and fixed with 70% ethanol. To aid mother-daughter separation, the samples were sonicated for 1 minute in a water bath before fixation.

Samples for DNA staining were digested with 5mg/mL proteinase K for 50 minutes at 50°C, followed by 2 hours of RNase A (Applichem, A2760,0500) treatment at 30°C. Samples were stained for DNA content using SYTOX Green (Thermofisher, S7020) diluted 1:5000 in PBS, and were sonicated in a water bath for 25 seconds before flow cytometry. Fluorescence was detected using a 488 nm excitation laser and a 525/15 nm emission filter. The PMT voltage was set up such that the sample with the highest expected ploidy did not saturate the signal.

### Data analysis

All analysis was performed using R,^81^ and the packages Bioconductor,^82^ dplyr,^83^ drc,^84^ MASS,^85^ mixtools,^86^ and ggplot2.^87^ All raw data and analysis scripts are available upon request.

Flow cytometry data was not gated except when necessary to remove debris. For aTc dose response experiments, median fluorescence of the entire population was used to fit a five-parameter dose response curve with the drm() command and the fplogistic formula 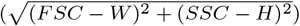 from the drc package. Parameters p_1_ and p_2_ were fixed individually for each curve, the rest of the parameters were estimated by the drm command. Parameter values can be found in Table S5. The cytometry cell volume proxy was always calculated as the magnitude of the vector of the FSC-W and SSC-H signals 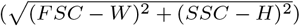, since forward and side scatter signals provide information about cell volume and budding state. The forward scatter width and side scatter height were chosen because this combination (as opposed to other combinations involving FSC-H/SSC-W or area of the signals) showed the most separation between measured signal peaks corresponding to spherical calibration beads of known diameter.

For single-reporter quantification of cell-to-cell variation, we calculated the residual standard deviation (RSD) of a linear model describing the relationship between the cytometry cell volume proxy and fluorescence of the population. To do this, the rlm() command from the MASS package was used to generate the linear model, and the residual standard deviation given by the same rlm() command was used as our measure of cell-to-cell variation. See Supplementary Text for a detailed explanation of the method.

Where shown, error bars for median fluorescence and the RSD were calculated using bootstrapping. The original set of data points was sampled with replacement and median fluorescence or RSD was calculated. 95% confidence intervals were calculated based on 1000 repetitions of this sampling process and plotted as error bars.

To generate a linear model describing the relationship between the volume proxy measurements done by flow cytometry and volume measurements by Coulter Counter, first the two data sets were sampled with replacement 5000 times. Then these samples were ordered by increasing volume proxy or volume and merged. The lm() command in the R package stats was used to fit the linear model. Sub-populations from Gates 4,6, and 9 were not included in the fitting, as these medians were deemed suboptimal representations of the bimodal distributions. The resulting linear fit had a slope of 471 and an intercept of 62032.

## Acknowledgements

We thank Jörg Stelling for conceptual discussions, Kohtaro Tanaka for useful discussions, Hans Michael Kaltenbach for discussions on the cell-to-cell variation measure, Kristina Elfström for initial construction of *P_2tet_* constructs and Robert Gnügge for the initial characterization of operator placements around the TATA sequence using LacI. This work was supported by the Swiss National Science Foundation as part of the NCCR Molecular Systems Engineering and by the grant R21CA223901 from NCI to RB.

## Supplementary Information

### Supplementary Text

#### Optimization of *tetO_1_* placement and endogenous Transcription Factor binding sites

When creating *P_7tet.1_*, the final promoter used in Well-tempered Controller_846_ (WTC_846_), we optimized the placement of the *tetO_1_* sequences, the sequences of the endogenous Gcr1 and Rap1 binding sites, the number of these sites present in the promoter, and the TATA sequence. Our goal was to increase maximum expression from the promoter, because *P_TDH3_* derivatives we had constructed with *tetO_1_* sites had shown reduced expression. For these optimizations we used as a starting promoter *P_4tet_*. This promoter is a variant of *P_5tet_* (Figure 1B) from which the *tetO_1_* sequence immediately upstream of the TATA was removed. *P_4tet_* showed a higher basal expression level than *P_5tet_*, allowing us to better observe subtle differences in basal expression (Figure S21B). We tested repression of *P_4tet_* and derivatives in the Simple Repression (SR) architecture. We constructed the derivatives as follows. We extended the Rap1 site and the upstream Gcr1 site by one base pair (*P_4tet.1_* and *P_4tet.2_*), to account for the possibility that we initially truncated the endogenous binding sites,^1^ replaced the downstream Gcr1 binding site with the same extended Gcr1 site (*P_4tet.3_*), since the upstream Gcr1 binding site was closer to the reported consensus sequence,^2^ and tested alternative TATA sequences (*P_4tet.4_* and *P_4tet.5_*).^3^ Four of these optimizations resulted in increased maximum activity (Figure S21B): the Rap1 site extension (*P_4tet.1_*), replacement of the downstream Gcr1 site (*P_4tet.3_*), and the TATA sequence optimizations (with *P_4tet.5_* driving expression more strongly than *P_4tet.4_*). The extended *P_4tet.2_* with the extended Gcr1 site showed reduced maximum expression.

We therefore chose the following modifications: the single base pair extension of the upstream Rap1 site, the replacement of the downstream Gcr1 site with the original upstream Gcr1 sequence, and the TATA sequence TATAAATA. We implemented these modifications to *P_5tet_* to generate *P_5tet.1_*. By the assays described in the main text for testing the other promoter derivatives, compared to *P_5tet_*, the new promoter *P_5tet.1_* (FRY2659) showed 95% of maximum expression driven by *P_TDH3_*, and increased repression (15 fold vs. 12 fold) with only a slight increase in basal activity when fully repressed (FRY2656, Figure 1B and Figure S21C). In order to increase maximum expression even further, we took advantage of previous work showing that increasing the number of transcription factor binding sites in a promoter could increase its strength.^4^ We investigated whether adding additional Rap1 and Gcr1 sites to *P_5tet.1_* would increase promoter activity. We created *P_7tet_* by duplicating the Rap1 site, and *P_7tet.1_* by duplicating both the Rap1 and one of the two Gcr1 sites in *P_5tet.1_*, while keeping the same *tetO_1_* placements at these duplicated sites (Figure S21C). *P_7tet.1_* had a higher maximum activity (116% vs 99% of *P_TDH3_* activity) and fold repression than *P_7tet_* (20 fold vs 18 fold), with only minimal increase in absolute repressed activity (4.3-fold vs 4 fold above autofluorescence). We therefore chose *P_7tet.1_* as the promoter for further use.

#### Single reporter quantification of cell-to-cell variation (CCV) in fluorescent protein expression by flow cytometry

In yeast and *C. elegans*, comparison of signals from strains with different combinations of different reporter genes allows the different contributions to variation to be independently quantified.^5, 6^ One of these contributions, individual differences in general ability to express genes into proteins, contributes to phenotypic variation in genetic penetrance and expressivity.^7^ Quantification of this and other sources of variation benefits from ability to measure output of single cells over time^5^ and, in flow cytometry, requires measurement of outputs of multiple reporters.^8^ Here, however, we were interested in the overall variability rather than specific sources of variability. Therefore, we only had a single reporter protein (Citrine). However single reporter studies have a major, confounding contribution to measured variation in gene expression that multi-reporter studies don’t: Fluorescent proteins in yeast are degraded very slowly unless they have degradation tags attached^9^ and therefore, if constitutively expressed, their abundance increases over time.^10^ Thus, in cycling populations of budding yeast that continually express fluorescent proteins, a major source of cell-to-cell variation in fluorescent signal is that small new-born cells have not had time to accumulate much fluorescent protein, while larger cells have. This source of variability normally affects all reporter proteins in the cell in a similar fashion, and therefore does not require correcting in multi-reporter studies. On the other hand in single reporter studies with flow cytometry in yeast, as for higher cells, this volume related variation in fluorescent protein expression is generally corrected for by gating; i.e. filtering the data to select only a narrow subset of cells with similar forward and side scatter, and thus volume, which increases with cell cycle progression. Such gating disregards data from the majority of the cells whose values fall outside the gated range. Here, in order to avoid discarding data, we established a single-reporter measure of cell-to-cell variation that corrects for variation due to fluorescent protein accumulation without gating.

We first established that forward and side scatter signals can be used to distinguish smaller cells from larger ones. We sorted cells on a BD FACS Aria III flow cytometer. We set different gates on the FSC and SSC signals (shown in Figure S22) to collect 10 sorted sub-populations, each containing about 100,000 exponentially growing FRY2683 cells. We then immediately measured a) FSC and SSC from the collected subpopulations on a different instrument, the LSRII Fortessa LSR used for the flow cytometric measurements in this work, and b) volume in fL with a Coulter Counter (Figure S23). The raw data acquired by the two methods can be seen in Figure S23A and S23B. For the flow cytometry data, we used the width of the FSC and height of the SSC to calculate a volume proxy using the formula 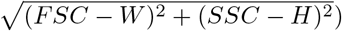 as explained in Materials&Methods. Figure 23C shows a linear relationship between the medians of the sub-populations as measured by the two methods, i.e., that the flow cytometric measurement is a proxy for volume, and that two volume measurements qualitatively agree. The three sub-populations (4,6, and 9) where a slight deviation from the linear relationship is observed are all bimodally distributed, meaning the population is a mixture of large and small cells and the median is not a good representation of this sub-population. Overall, for the cell-to-cell variation calculations outlined below, this relative relationship is enough to distinguish new-born, smaller cells from larger cells that have had time to accumulate fluorescent protein.

We then used this information to calculate the CCV in fluorescent protein expression that could not be attributed to differences in cell volume/cell cycle progression. We began with FRY2683 cells, which express Citrine from wild type *P_TDH3_*. We then plotted the volume proxy vs. the fluorescence signal observed in the entire population measured by flow cytometry (Figure S3). Then we performed a robust linear fit on the cell volume proxy vs. the Citrine fluorescence signal. This linear model allowed us to correct for the differences in cell volume and calculate the Residual Standard Deviation (RSD) of the fit as explained in the Materials&Methods. This RSD value quantifies the variation in the population that is not due to differences in cell volume between the cells. While use of this measure is in principle akin to measuring variation in expression of fluorescent proteins using a very narrow gate on the measured FSC vs SSC signals of the population, it avoids the need to discard data. Moreover, it could also be used when comparing populations with different cell volume distributions.

#### Increasing the mass and nuclear concentration of TetR in order to abolish basal expression from *P_7tet.1_*

We reasoned that basal expression from *P_7tet.1_* in the Complex Autorepression (cAR) architecture might arise because a) the nuclear concentration of TetR might be too low for all of the *tetO_1_* TetR binding sites in *P_7tet.1_* to be occupied at all times, and/or b), that TetR derivatives might fully occupy all of the operators and yet not repress completely. We tested the first idea by increasing the nuclear concentration of TetR proteins by expressing derivatives that contained a second SV40 Nuclear Localization Sequence. We tested the second idea by fusing TetR to other protein moieties that might aid repression. Specifically, we added to TetR portions of prokaryotic proteins that we could presume to be inert, hoping that these bulkier TetR derivatives might repress more strongly, for example by better sterically interfering with the binding of transcription factors, or with contacts between Gcr1 and Rap1 at the Upstream Activating Region (UAS) and the transcription apparatus at the core promoter. We tested the efficacy of these new molecules by expressing them from *P_ACT1_* in the SR architecture (FRY2681, FRY2664, FRY2665, FRY2666, and FRY2667, Figure S24). For smaller repressors, addition of a second NLS decreased uninduced expression whereas for larger repressors it did not. The strain carrying the TetR-nls-MBP (Maltose Binding Protein, the *coli* malE gene product), showed the most repression, but still exhibited uninduced expression signal of 2.2-fold above autofluorescence background.

**Figure 1:**
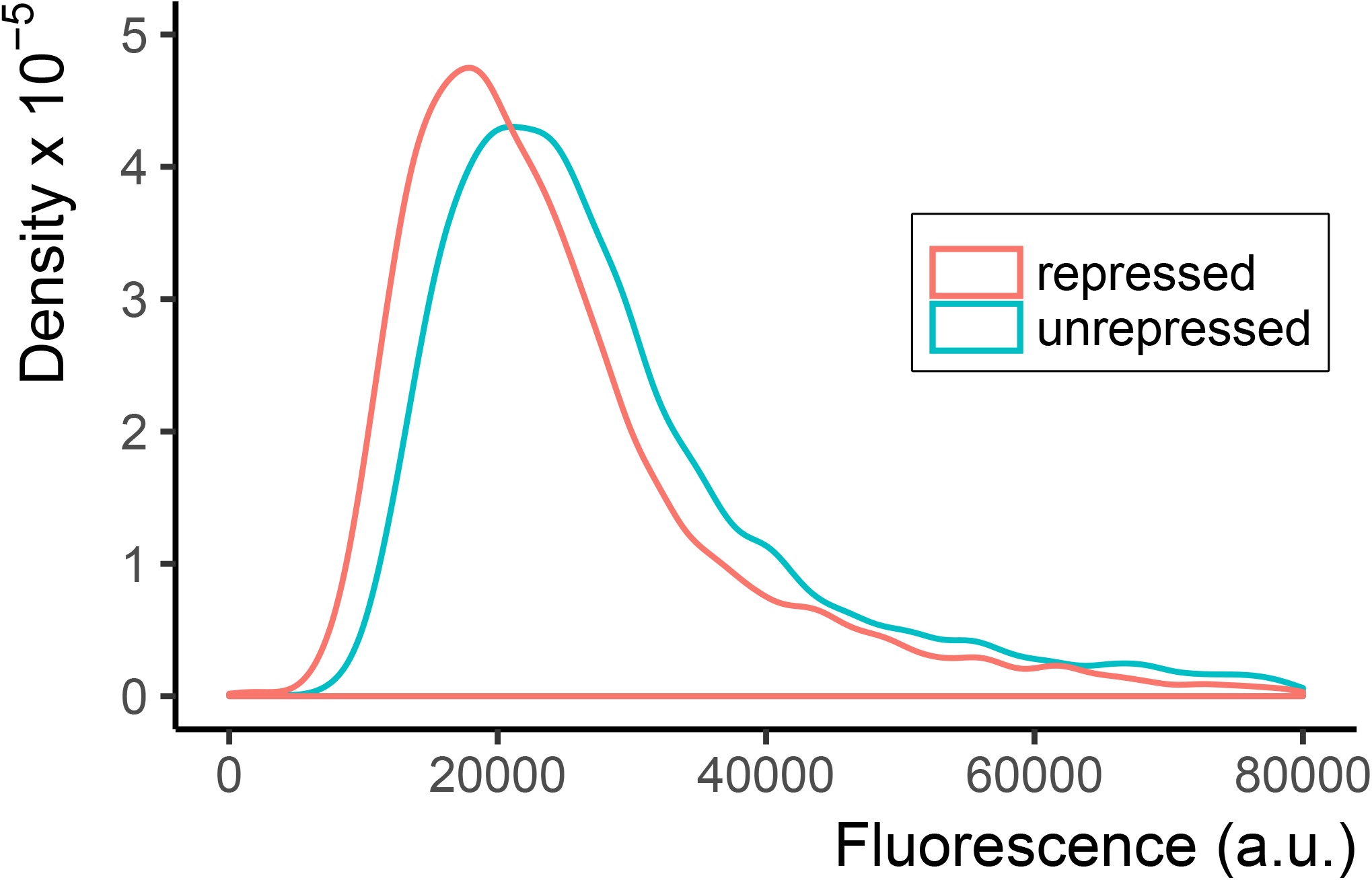
A promoter with three *tetO_1_* sequences in the UAS of *P_TDH3_* is only minimally repressed by TetR. We constructed *P_3tet_*, a *P_TDH3_* derivative that carried three *tetO_1_* TetR binding sites, adjacent to the endogenous transcription binding sites (one for Rap1, two for Gcr1) in the UAS. We used this to construct FRY2564, a strain that carried the *P_3tet_*-Citrine construct integrated into the LEU2 locus (unrepressed), and the otherwise isogenic strain FRY2573, in which a *P_ACT1_*-TetR construct was additionally integrated at the HIS3 locus (repressed). We measured Citrine fluorescence using flow cytometry. Plot shows density of cells at each fluorescence value, such that the area under the curve is 1. By comparison of medians, as described, repression by TetR was only 1.5-fold. As described in the main text, the inference from this result is either that TetR cannot efficiently prevent endogenous transcription factors Rap1 and Gcr1 from occupying their binding sites in the UAS, or that interfering with binding to UAS sites without blocking PIC assembly near the TATA-sequence is not sufficient to significantly repress transcription from this promoter.

**Figure 2:**
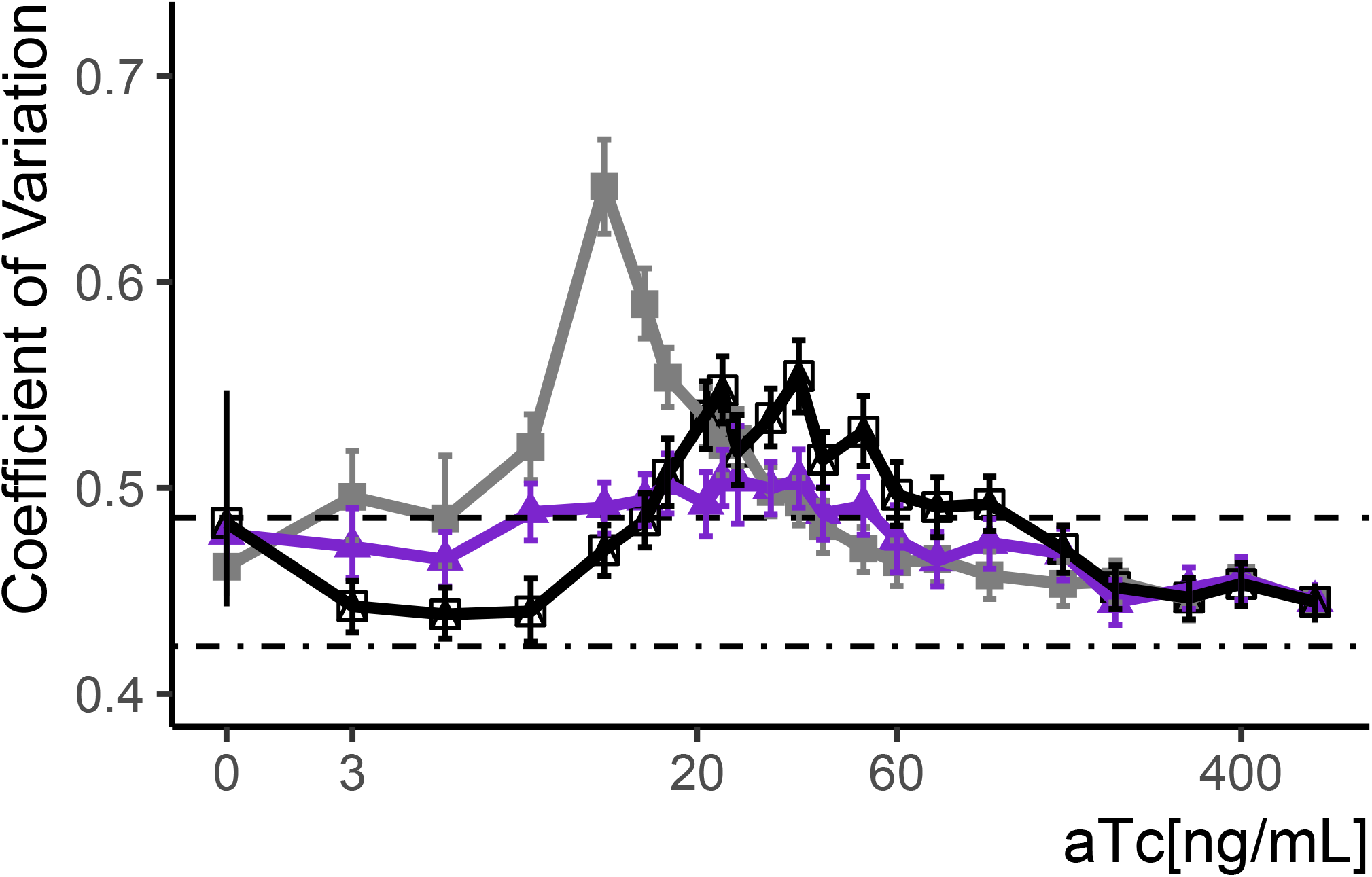
Variation in expression for the SR, AR, and cAR architectures. We calculated the CoV in fluorescence signal using the data shown in main Figure 2B, to test whether the volume-corrected RSD measure described in Supplementary Text and Figure S3 agrees with it. The plot shows observed CoV at each aTc concentration tested for the strains carrying different architectures. SR(FRY2663), grey, AR(FRY2674), purple, and cAR(FRY2741), black. Bootstrapping (n=1000) as described in Materials&Methods was used to estimate 95% confidence intervals as indicated by the error bars. Dashed line indicates the CCV of the parent cell (autofluorescence control, FRY70) and dot-dash line the strain where Citrine was constitutively expressed from *P_TDH3_* (FRY2683). The CoV provides a direct measure of the total CCV in expression, and includes variation due to larger cells having had a longer time to produce more fluorescent protein. For each architecture, both this simple CoV and the RSD in Main Figure 2B show maximum variation at the same aTc concentrations.

**Figure 3:**
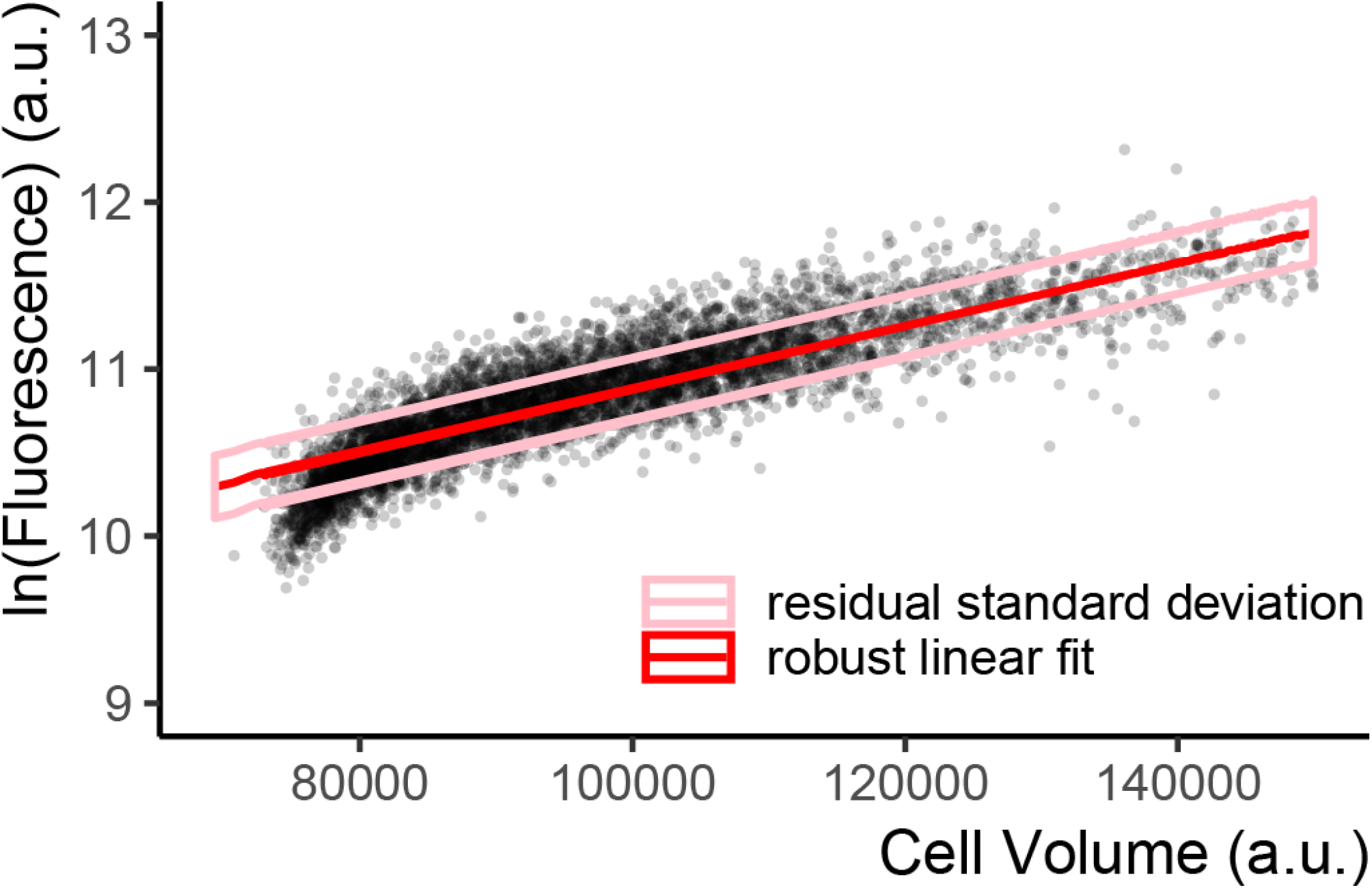
Single-reporter RSD measure of CCV in expression. Scatter plot of flow cytometry cell volume proxy vs. Citrine fluorescence in a strain where Citrine is constitutively expressed from *P_TDH3_* at the LEU2 locus (FRY2683) shows a positive correlation between the two quantities. Fluorescence and the volume proxy are positively correlated because, in cycling populations larger cells have had more time to express more fluorescent protein. Each point represents a single cell. Volume proxy was calculated as the vector of the SSC-H and FSC-W signals (Materials&Methods). The red line indicates the robust linear fit, the ideal linear correlation between volume and fluorescence in this population if there was no cell-to-cell variation. Pink lines represent +1 and −1 one residual standard deviation from this fitted correlation line. The RSD can be interpreted as the fraction of variability in expression in the population that is not explained by the cell volume proxy/inferred progress through the cell cycle.

**Figure 4:**
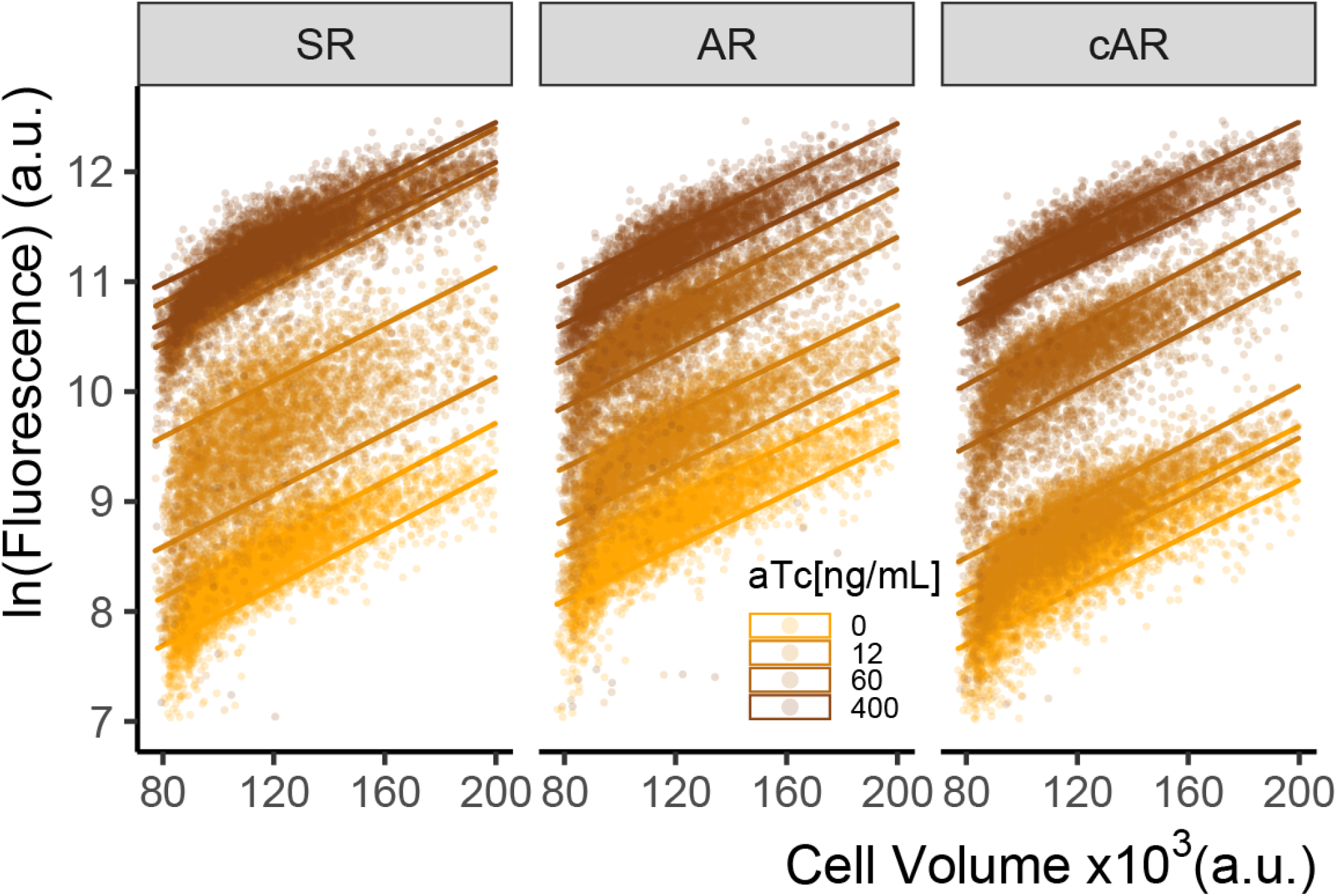
AR architecture reduces cell-to-cell variability at intermediate concentrations of aTc. Plot of flow cytometry cell volume proxy vs Citrine fluorescence in whole (ungated) populations of cells of SR,AR and cAR architectures (FRY2663,2674,2741) grown for 7 hours at different aTc concentrations. Each point indicates an individual cell. The two lines of the same color are drawn at +1 and 1− RSD for the fitted robust line of each population, same as in Figure S3. By this measure, at intermediate induction concentrations, the SR architecture shows around 2-fold increased cell-to-cell variability compared to AR and cAR architectures.

**Figure 5:**
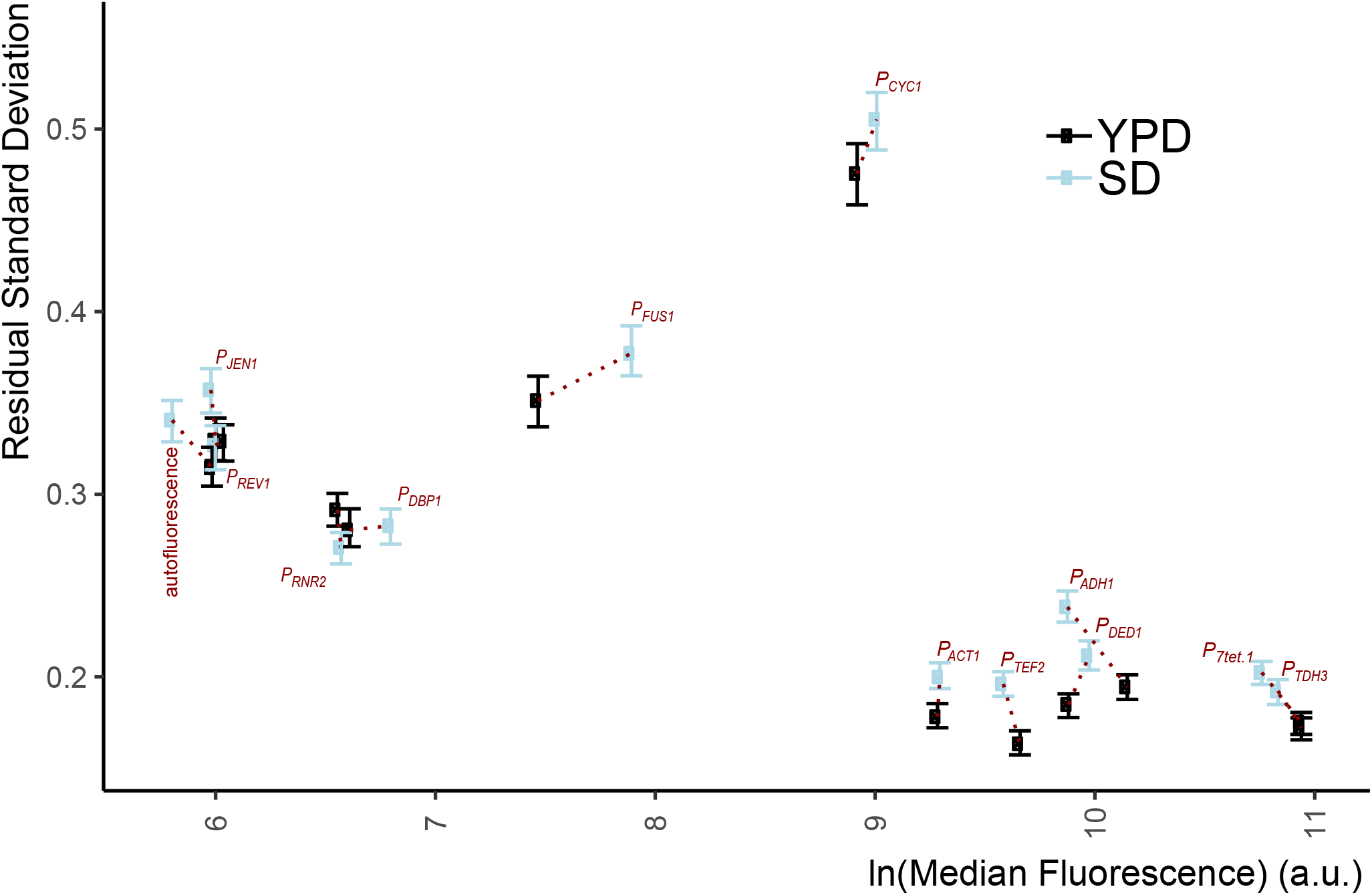
Single-reporter RSD measure of variation in expression from native yeast promoters. We measured Citrine expression and its RSD in otherwise isogenic haploid cells that carried integrated constructs at various auxotrophic marker loci, in which the indicated native yeast promoters drove Citrine expression. All strains used here are indicated in Table S1 as “promoter name-const”. Autofluorescence was strain FRY70. We grew cells to exponential phase in Rich medium (YPD) and synthetic medium (SD Full (minimal glucose media with complete amino acid complement)) and measured fluorescent signal with flow cytometry. RSD values were calculated as explained in Supplementary Text. Error bars indicate 95% confidence intervals on the RSD measurement as calculated by bootstrapping (See Materials&Methods). For most promoters, the the CCV is lower in YPD. For all strong promoters except *P_CYC1_*, CCV is around 20%. For promoters with lower endogenous activity (as inferred from expression levels of the endogenous proteins they control^11^) RSD variation was roughly 30%, similar to the RSD variation measured from the autofluorescence signal of the strain without Citrine. For *P_CYC1_*, measured RSD variation was about 50%.

**Figure 6:**
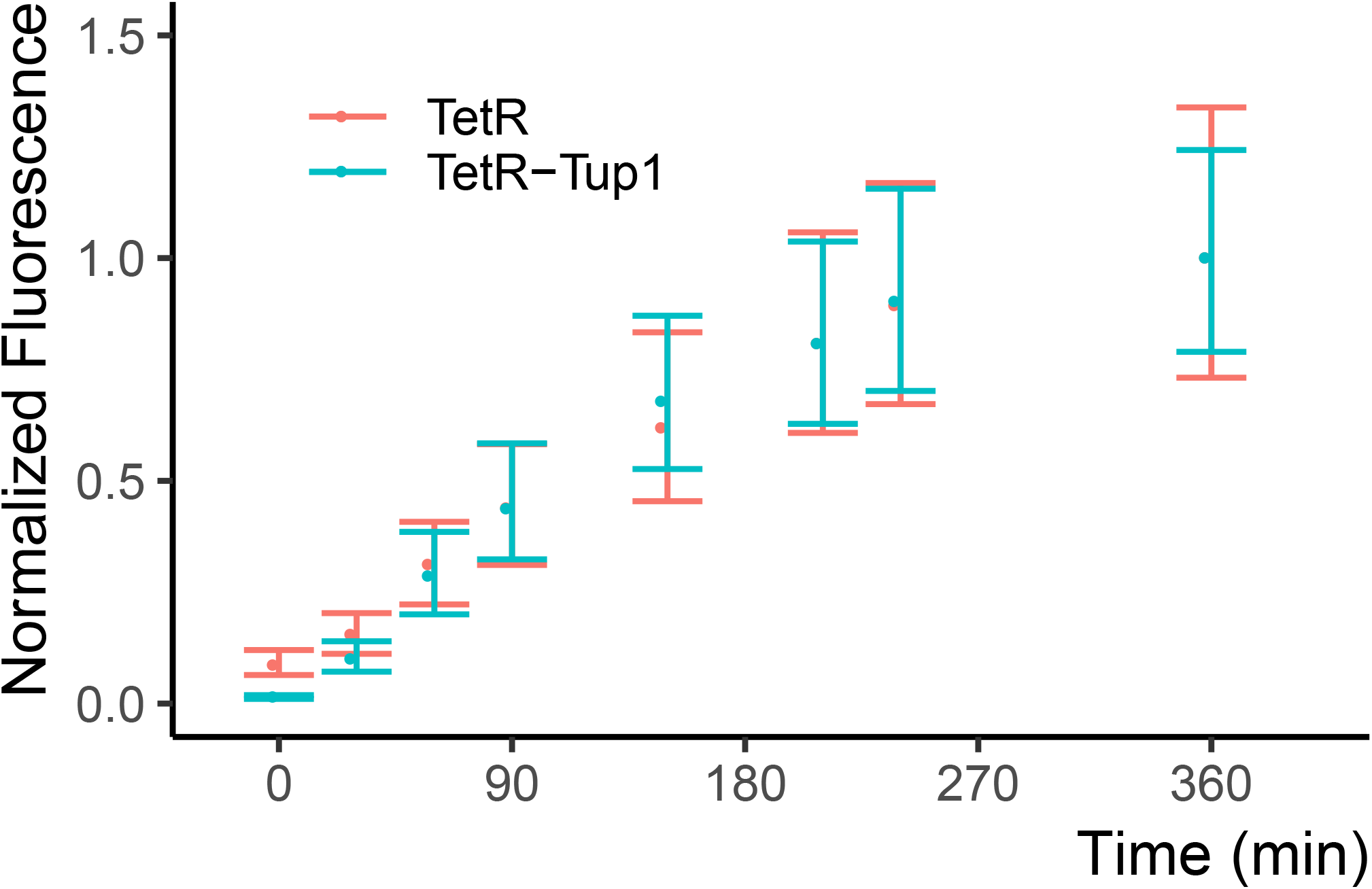
The zeroing repressor TetR-nls-Tup1 does not affect the induction speed of *P_7tet.1_*. Citrine signal after induction in *P_7tet.1_*-Citrine strains. Both strains carried a *P_7tet.1_*-Citrine integrated at the LEU2 locus, in conjunction with two different repressors. One strain (red, FRY2674) carried an autorepressing *P_7tet.1_*-TetR construct integrated at the the MET15 locus. The second strain (blue, FRY2717) was otherwise isogenic, but had no TetR. Instead it carried a constitutively expressed *P_RNR2_*-TetR-nls-Tup1 construct integrated at HIS3. We induced both strains with 600 ng/mL aTc at time zero during exponential growth phase. We quantified Citrine fluorescent signal in flow cytometry, and normalized to maximum level reached at 360min. The dots indicate the median normalized fluorescence, error bars span the range between the first and the third quartiles. The results show that TetR-nls-Tup1 does not reduce induction speed, nor the time needed to reach steady state.

**Figure 7:**
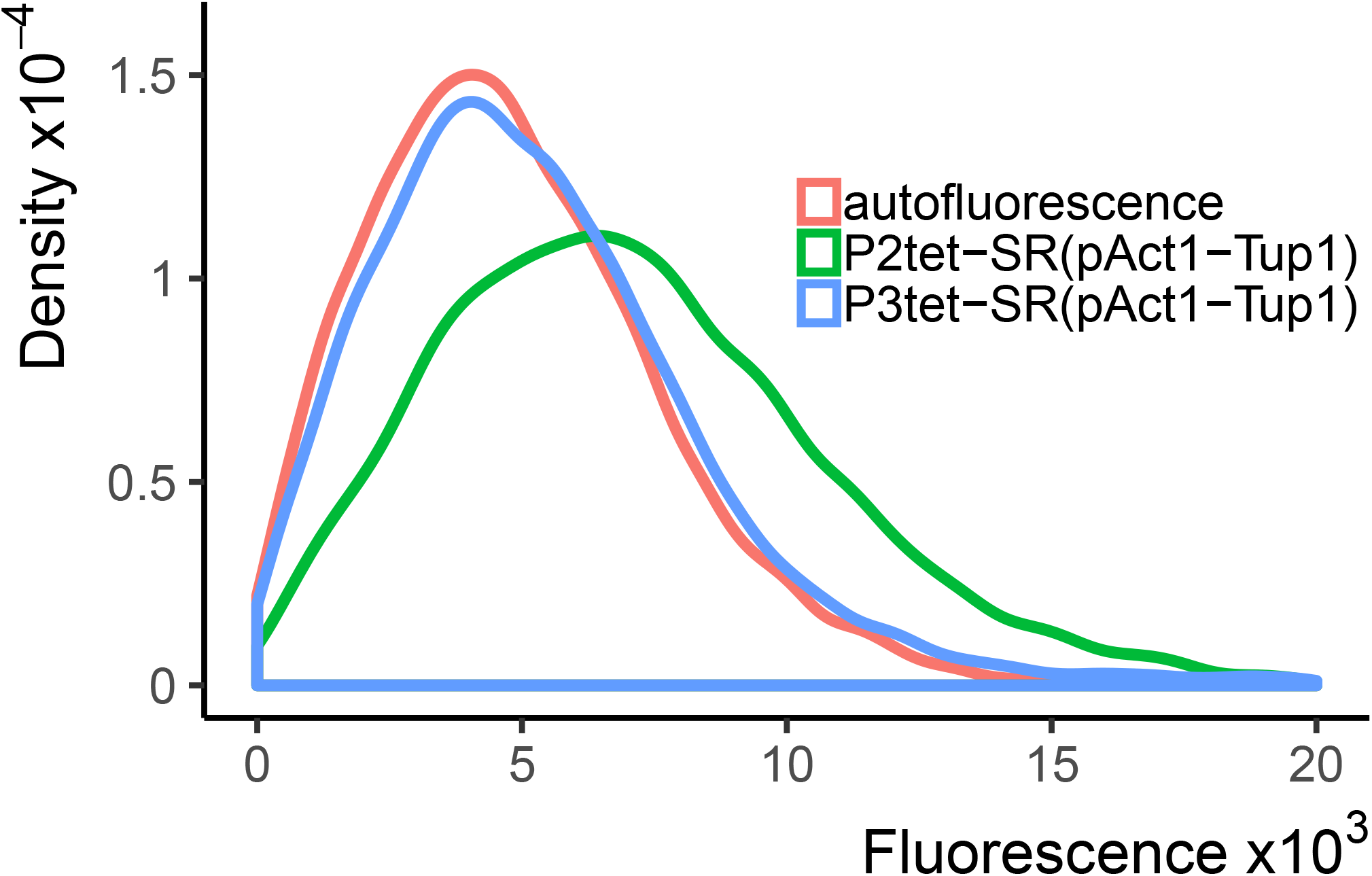
A TetR-nls-Tup1 fusion protein fully represses expression when binding only at the UAS, but not only at the TATA. Strains carried a *P_ACT1_* construct that directed the expression of TetR-nls-Tup1 integrated at HIS3 locus. FRY2702 (blue line) also carried *P_3tet_*-Citrine construct, which bears three TetR binding sites adjacent to each Gcr1 and Rap1 binding site in the UAS, integrated at LEU2. FRY2662 (green line) carried *P_2tet_*-Citrine construct, in which TetR binding sites flank the TATA-sequence, integrated at LEU2. FRY70 (red line) is a control strain with no integrated constructs. Repressed expression was measured using flow cytometry at maximum voltage, i.e. maximum sensitivity. Plot shows density of cells at each fluorescence value, such that the area under each curve is 1. TetR-nls-Tup1 completely repressed fluorescent signal driven by the *P_3tet_*, in which the TetR sites are in the UAS, but not from *P_2tet_* construct, in which the TetR binding sites flank the TATA-sequence.

**Figure 8:**
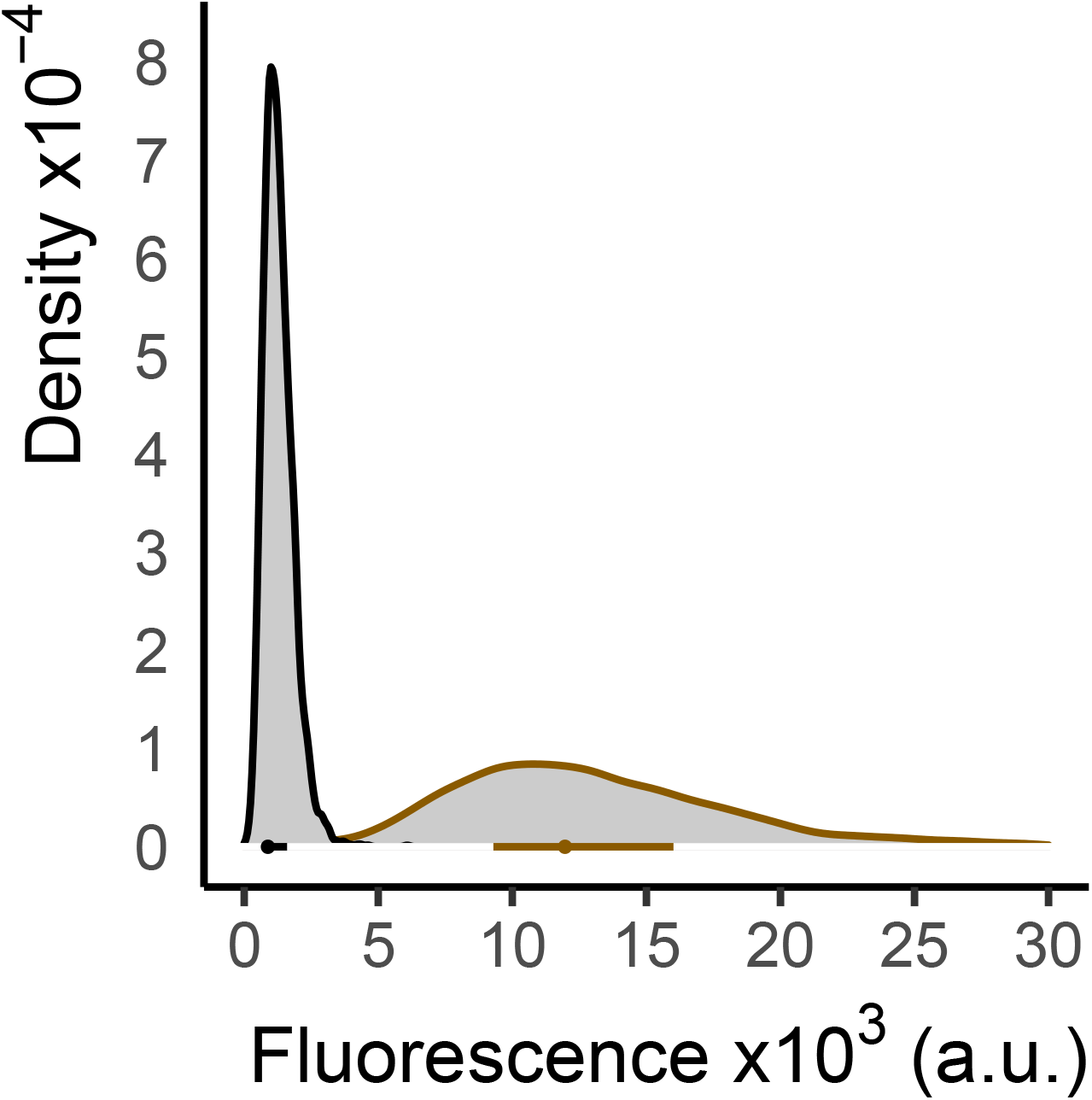
Low level TetR-nls-Tup1 expression results in incomplete repression of *P_7tet.1_*. Fluorescence signal from a strain where a *P_7tet.1_*-Citrine construct was integrated at the LEU2 locus and repressed by *P_REV1_*-driven TetR-nls-Tup1 integrated at HIS3 and *P_7tet.1_* driven TetR integrated at the MET15 locus (brown, FRY2715) was measured using flow cytometry, and autofluorescence signal (black, FRY70) was measured from an otherwise isogenic strain that lacked Citrine, TetR and TetR-nls-Tup1 expression. Plot shows density of cells at each fluorescence value, such that the area under the curve is 1. The dot inside the plot indicates the median, the bar spans the range from the first to the third quartile. TetR-nls-Tup1 can only repress *P_7tet.1_* activity to 10-fold above autofluorescence when expressed from *P_REV1_* in the cAR architecture. This result indicated that the TetR-nls-Tup1 needed to be expressed at a higher level to fully repress *P_7tet.1_*. Taken together with the results presented in Figure 3B, this result suggests that the lowest abundance of TetR-nls-Tup1 able to fully repress *P_7tet.1_* lies between that expressed from constructs driven by *P_REV1_* and constructs driven by *P_RNR2_*.

**Figure 9:**
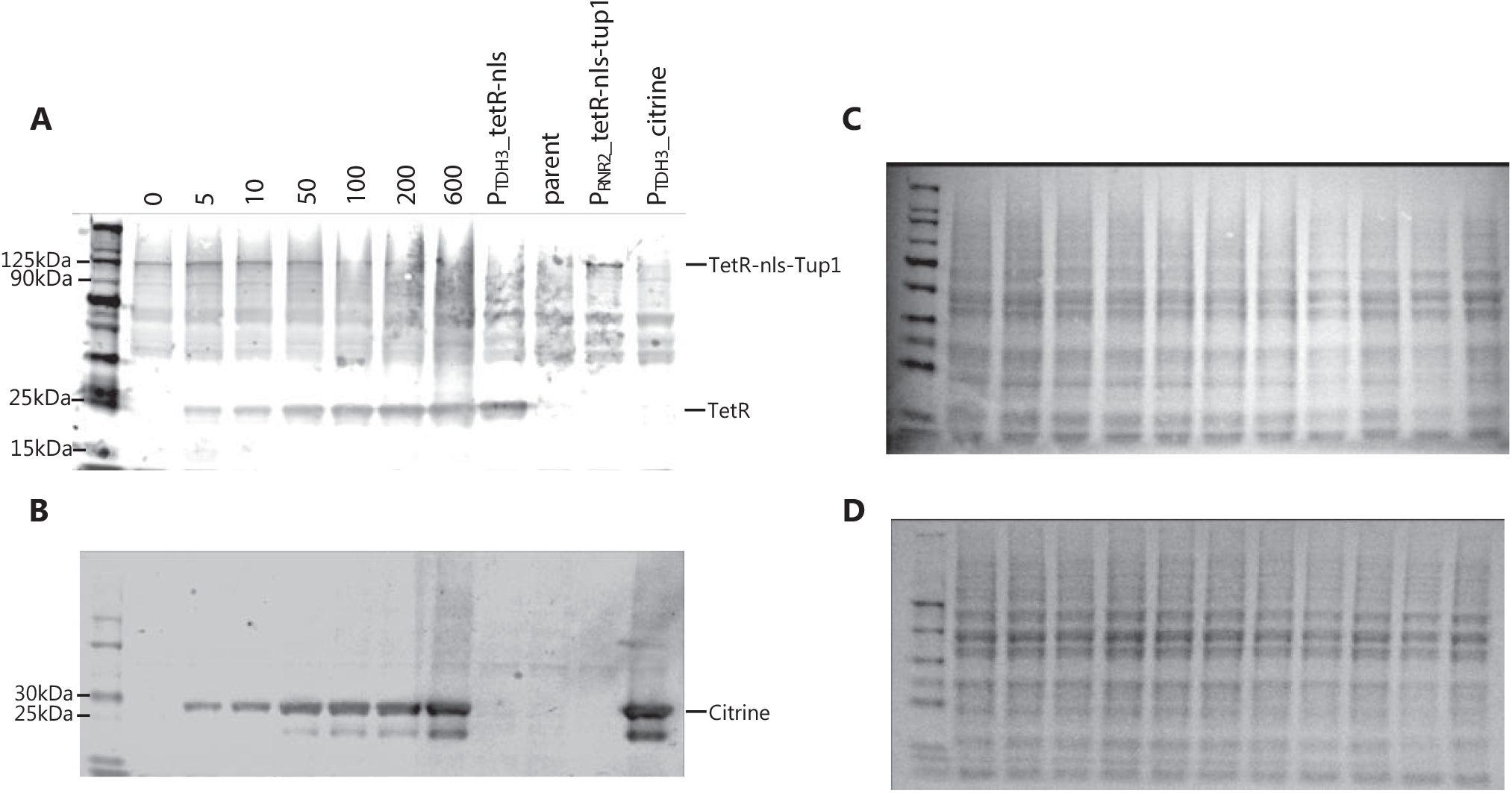
Direct observation of dose response for WTC_846_-controlled protein expression. The strain *WTC_846_::Citrine* (FRY2759), in which both TetR and Citrine were expressed from *P_7tet.1_*, and TetR-nls-Tup1 was expressed constitutively from *P_RNR2_*, was grown in YPD to stationary phase with different aTc concentrations. TetR and TetR-nls-Tup1 were integrated at the URA3, and the *P_7tet.1_*-Citrine construct at the LEU2 locus. Protein extracts from 2.5 million cells were loaded per lane for Western blotting from either these samples, or from otherwise isogenic control strains that constitutively expressed either TetR (FRY2611) or Citrine (FRY2683) under the control of *P_TDH3_*, TetR-nls-Tup1 (FRY2717) under the control of *P_RNR2_* (HIS3 locus), or no non-endogenous protein was expressed (FRY70). In this experiment, we used Ponceau staining to confirm uniform sample loading (C,D). The primary antibodies were mouse monoclonal anti-TetR and anti-GFP. The secondary antibody was IRDye 800CW Goat anti-Mouse IgG. We quantified protein level by infrared signal in the the Li-Cor reader as explained in Materials&Methods. Numbers indicate aTc concentrations in ng/ mL. In FRY2759, both the TetR(A) and Citrine(B) expression was higher at higher aTc concentrations, and neither protein was expressed in the absence of aTc. The result shows that uninduced WTC_846_ has no detectable repressed expression of either protein at the protein level, and the level of the expressed protein can be precisely adjusted.

**Figure 10:**
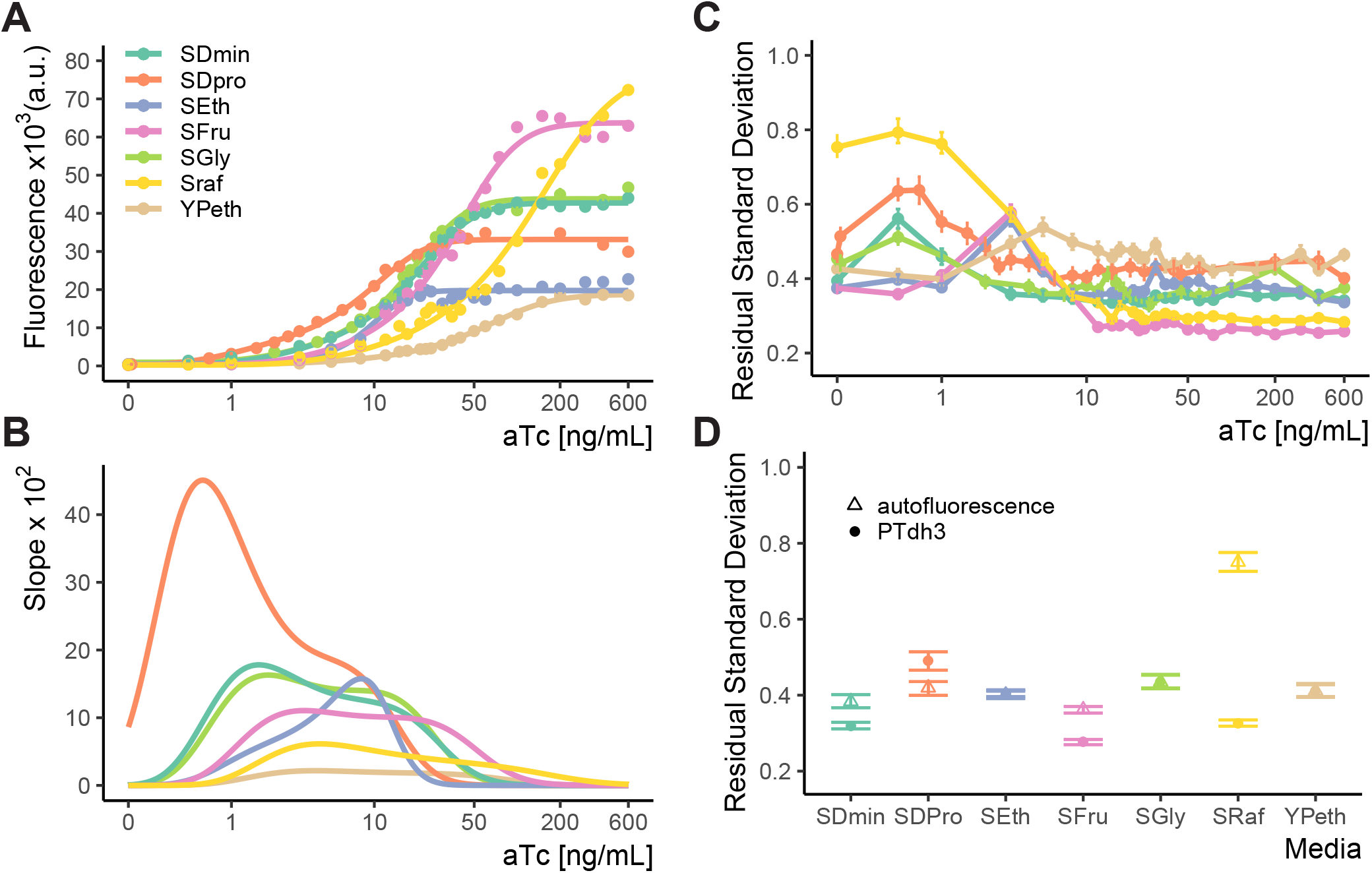
Dose response of WTC_846_-controlled expression in cells grown in different media. (A) Strain carrying *WTC_846_::Citrine* construct (FRY2759) grown in different media. Dots represent the median fluorescence of each population, and the lines were fitted using a 5 parameter log-logistic model as explained in Materials&Methods. Since dose responses were measured in separate experiments, the absolute fluorescence values of cells grown in different media are not comparable. (B) Slopes of the fit of the dose response curves depicted in (A). This plot provides a visual estimate of the input dynamic range, which is the range of aTc concentrations where the slope is non-zero. (C) Dose response of RSD measure of CCV in *WTC_846_::Citrine* expression in cells grown in different media (in A) at different aTc doses, calculated as in Materials&Methods and Supplementary Text. (D) CCV of two control strains in the same media conditions: one (FRY2683), that carried a *P_TDH3_*-Citrine construct integrated at LEU2, and a parent strain without any non-endogenous gene expression (FRY70, labelled autofluorescence). Where present, error bars indicate 95% confidence interval calculated using bootstrapping (n=1000) as explained in Materials&Methods. In all media, cell to cell variation in WTC_846_-driven Citrine expression is less than the variability measured in autofluorescence and comparable to that for constitutive expression throughout the majority of the precisely titratable input dynamic range. This shows that WTC_846_ can reliably titrate proteins under different media conditions.

**Figure 11:**
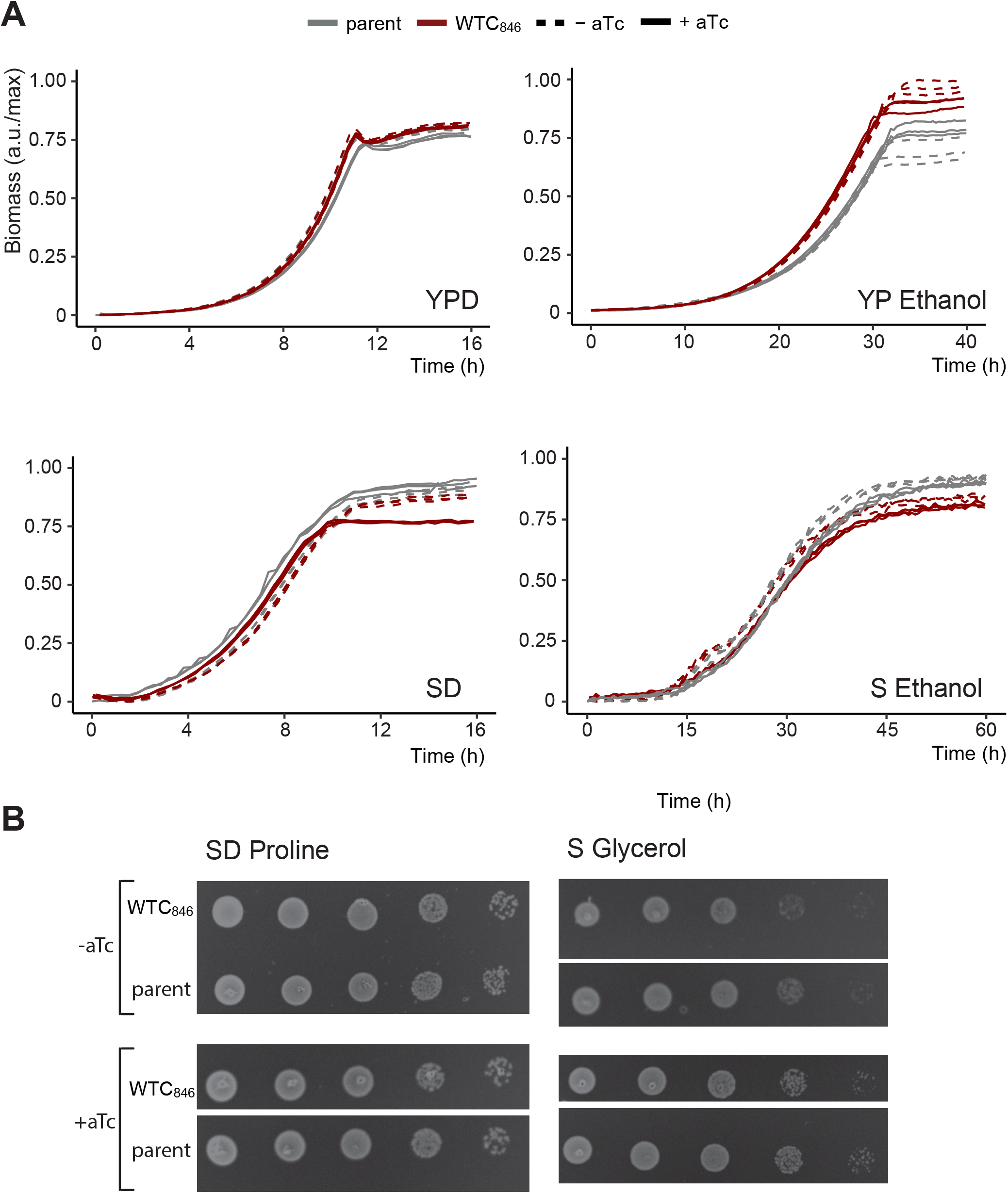
WTC_846_-directed expression does not affect gross measures of cell physiology. We measured the effect of WTC_846_ on growth rate and on cell viability measured by ability of single cells to form colonies. To do so, we used the WTC_846_:: strain FRY2761, in which a construct bearing *P_7tet.1_*-TetR and *P_RNR2_*-TetR-nls-Tup1 was genomically integrated at the URA3 locus, but no controlled gene was present. Growth compared to that for FRY70, an otherwise isogenic parent in which these components were not integrated, was used as a proxy for cellular physiology and wellbeing. (A) Growth curves were obtained using back scatter using a Biolector for YPD, SD Full and YP Ethanol with a total culture volume of 1mL, a Growth Profiler (Enzyscreen) for S Ethanol with a total culture volume of 250*μ*L. In (A), y axis shows culture density as measured by the device in arbitrary units. Values were normalized such that the highest value recorded corresponds to 1. Seeding density for liquid cultures were 50,000 cells per mL for all media except when ethanol was used as carbon source, in which case it was 500,000 cells per mL. (B) Colony formation assays for SD Proline and S Glycerol. Spotting assays were on solid SD Proline and S Glycerol media as described. For spotting assays, 500,000 cells were spotted on the left column, and each subsequent spot was from a 1:10 dilution in cell numbers. Pictures were taken after 48h of growth at 30°C. These assays showed no significant differences between the parent and the WTC_846_ strain, except in YP Ethanol, in which the WTC_846_ strain consistently reached a slightly higher density for unknown reasons.

**Figure 12:**
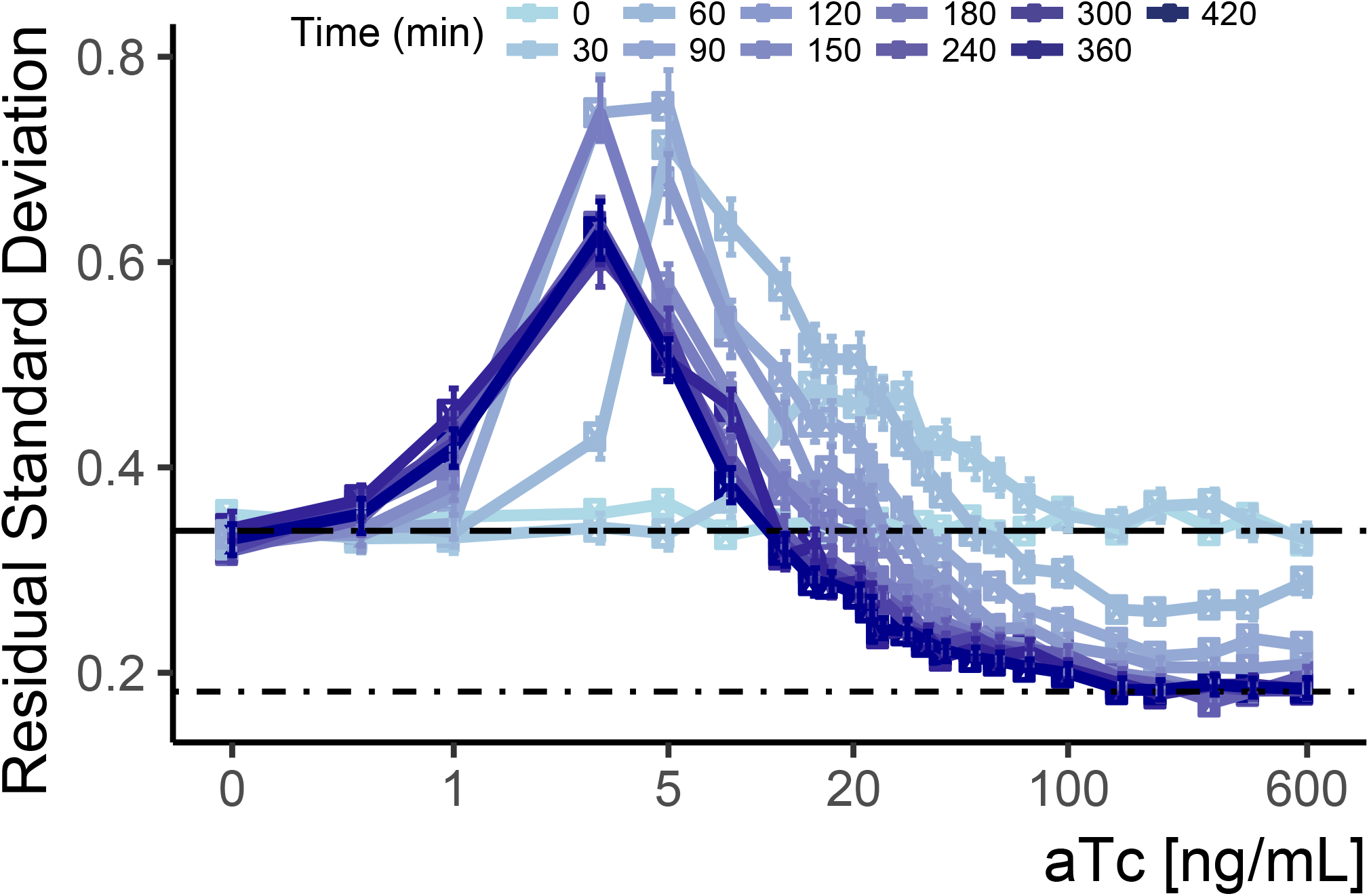
Cell-to-cell variability of WTC_846_ driven expression during induction. We used data from the Time Dependent Dose Response (TDDR) experiment shown in Figure 4B to calculate single reporter CCV during the course of induction of the *WTC_846_::Citrine* strain (FRY2759) at different doses of aTc. We compared this with CCV in autofluorescence (dashed line, FRY70) and in constitutive Citrine signal driven by a *P_TDH3_*-Citrine construct integrated at LEU2 (dot dash line, FRY2683). Error bars indicate 95% confidence interval calculated using bootstrapping (n=1000) as explained in Materials&Methods. RSD was calculated as explained in Supplementary Text. At earlier time points, the peak in CCV is observed at higher doses than steady state, but CCV stabilizes at steady state levels within 2 to 2.5 hours after induction.

**Figure 13:**
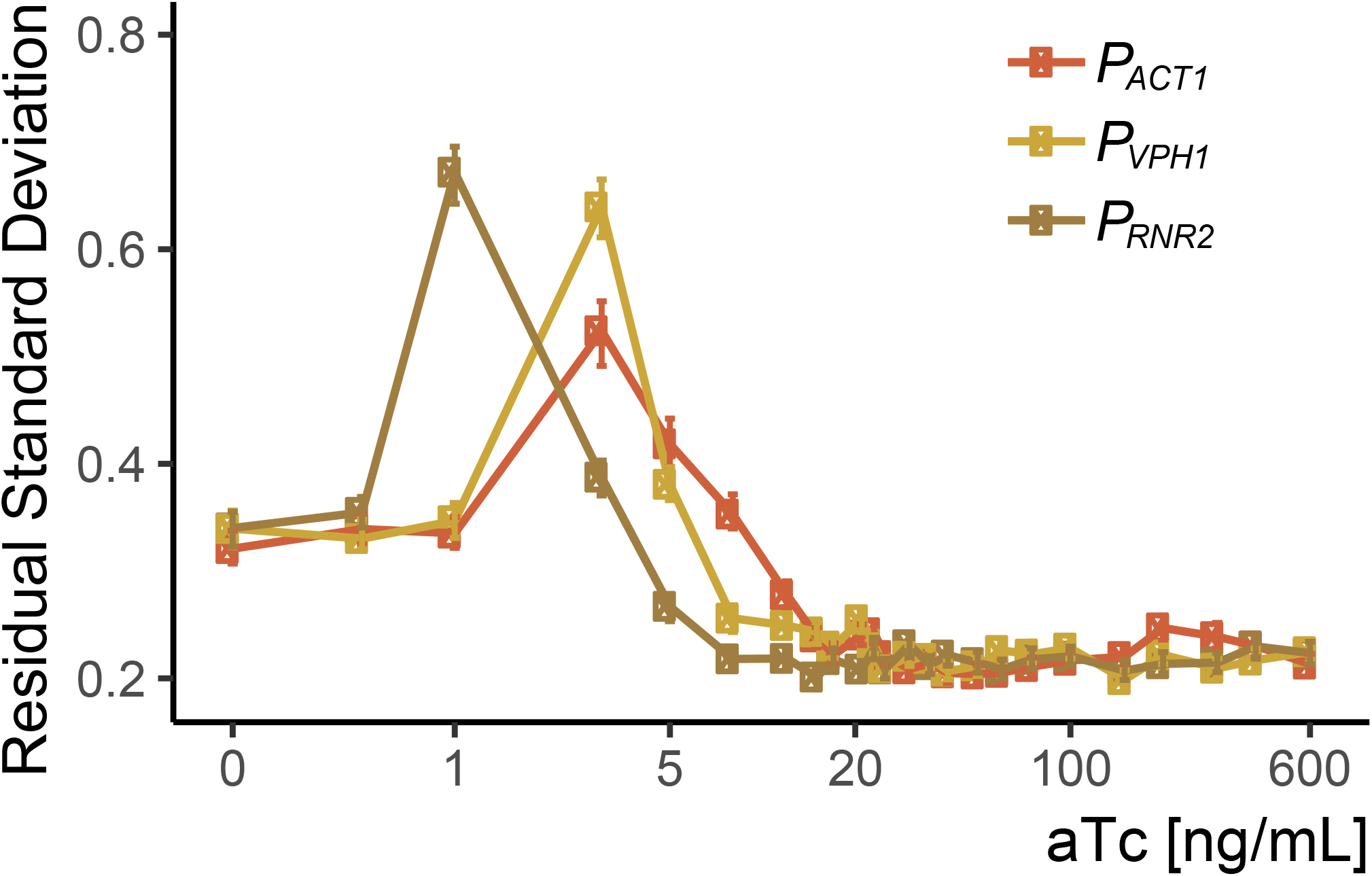
Peak CCV in SR strains corresponds to higher doses at higher expression levels of TetR-nls-Tup1. We used data from the dose response experiment shown in Figure 3C to calculate single-reporter RSD CCV as described in the Supplementary Text. We induced three strains (FRY2669, FRY2676, and FRY2717) which carried a *P_7tet.1_*-Citrine construct integrated at LEU2, and which carried constructs integrated at HIS3 in which the *P_ACT1_*, *P_VPH1_* and *P_RNR2_* promoters drove TetR-nls-Tup1 expression. Induction was done with different concentrations of aTc and Citrine fluorescence signal was measured after seven hours using flow cytometry. Error bars indicate the 95% confidence interval as calculated by bootstrapping (See Materials&Methods). For each strain, peak cell-to-cell variability corresponds to the steepest part of each dose response curve in Figure 3C, and occurs at higher aTc concentrations at higher TetR-nls-Tup1 expression levels (TetR-nls-Tup1 expression level inferred from the endogenous protein abundance of Act1, Vph1 and Rnr2^11^).

**Figure 14:**
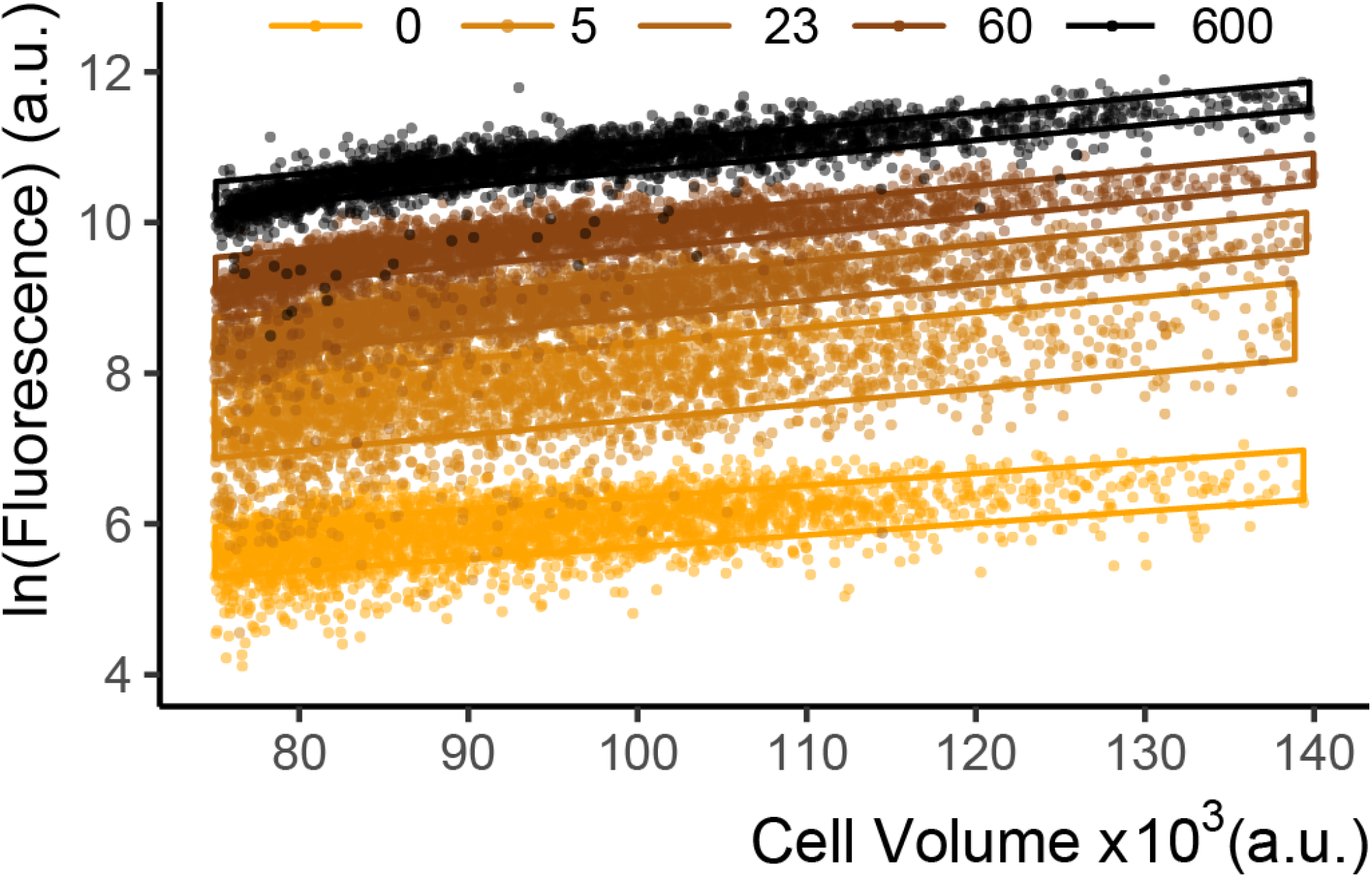
Fluorescence and volume of the *WTC_846_::Citrine* strain induced with different aTc concentrations. We grew cells of *WTC_846_::Citrine* strain (FRY2759) to exponential phase, and measured Citrine fluorescence with flow cytometry 7h after induction with aTc. The complete dose response is presented in main text Figure 4B. For this plot, we calculated the cell volume proxy as the vector of SSC-H and FSC-W signals as described (Materials&Methods and Supplementary Text). Each dot represents a single cell. The boxes were drawn as explained in Figure S3. The height of these boxes spans +/− one RSD and indicate the magnitude of CCV in the population. As explained in main text Figure 4D, at lower aTc concentrations the CCV is slightly higher due to the effect of the SR regime of the cAR architecture, however for most of the input dynamic range, variation in fluorescence at a given volume is low, as was observed for the AR architecture in main Figure 2D.

**Figure 15:**
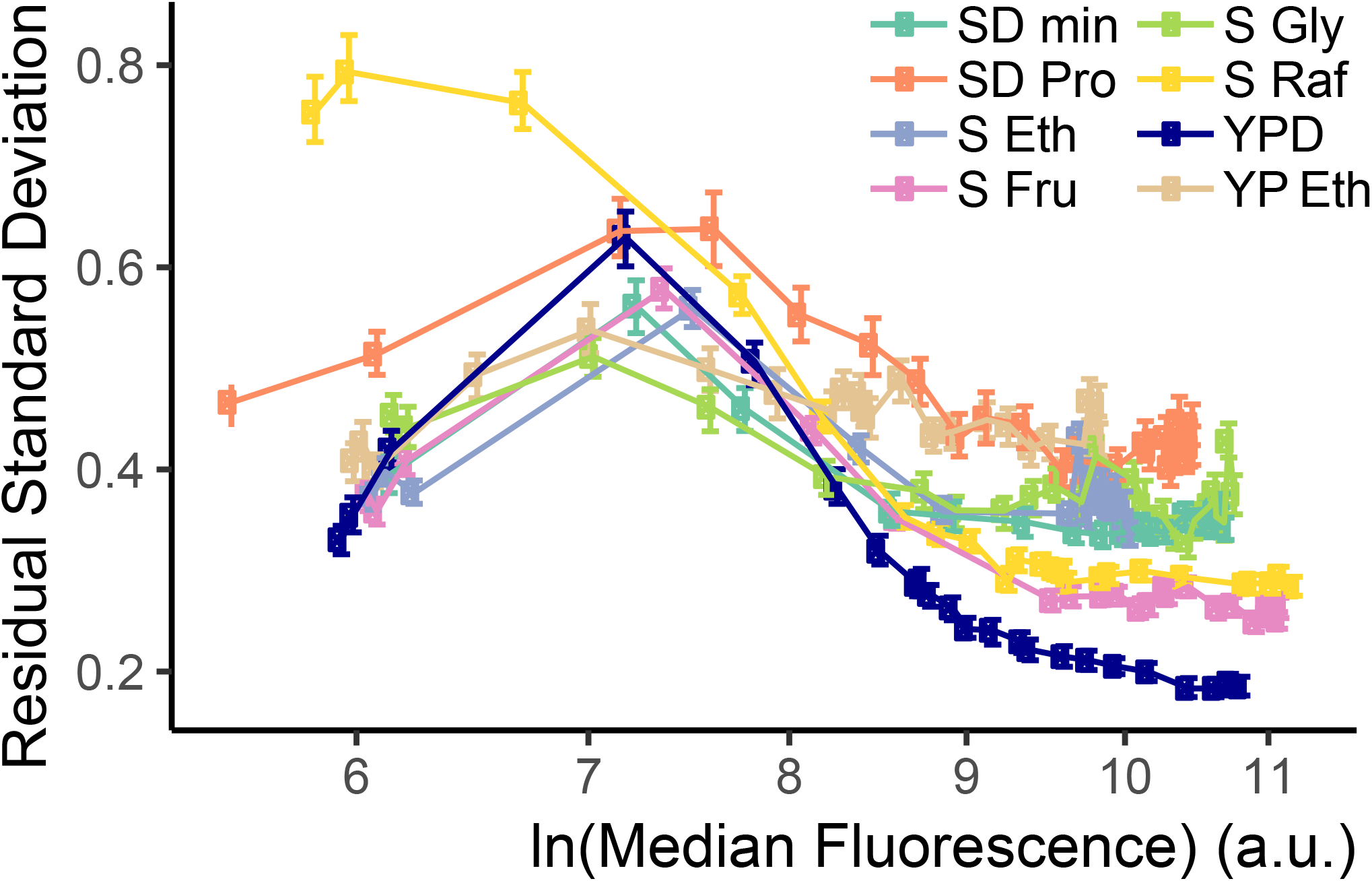
CCV in WTC_846_-controlled expression in cells grown in different media. CCV of WTC_846_ was measured in different media conditions using the *WTC_846_::Citrine* strain (FRY2759). Cells were grown in different media at different aTc concentrations, and Citrine fluorescence was recorded with flow cytometry at steady state. Time to reach steady state depends on the growth rate of the cells in the media condition, and is longer the slower the growth rate is. We calculated the single reporter RSD measure of CCV as explained in the Supplementary Text. Error bars indicate 95% confidence interval calculated using bootstrapping (n=1000) as explained in Materials&Methods. In all media, CCV peaked at around the same level of expression (same fluorescence signal value), and was lower at higher expression levels. For all media, variation in signal at higher expression levels was comparable to that for a *P_TDH3_*-Citrine construct integrated at the LEU2 locus (FRY2683, shown in Figure S10D).

**Figure 16:**
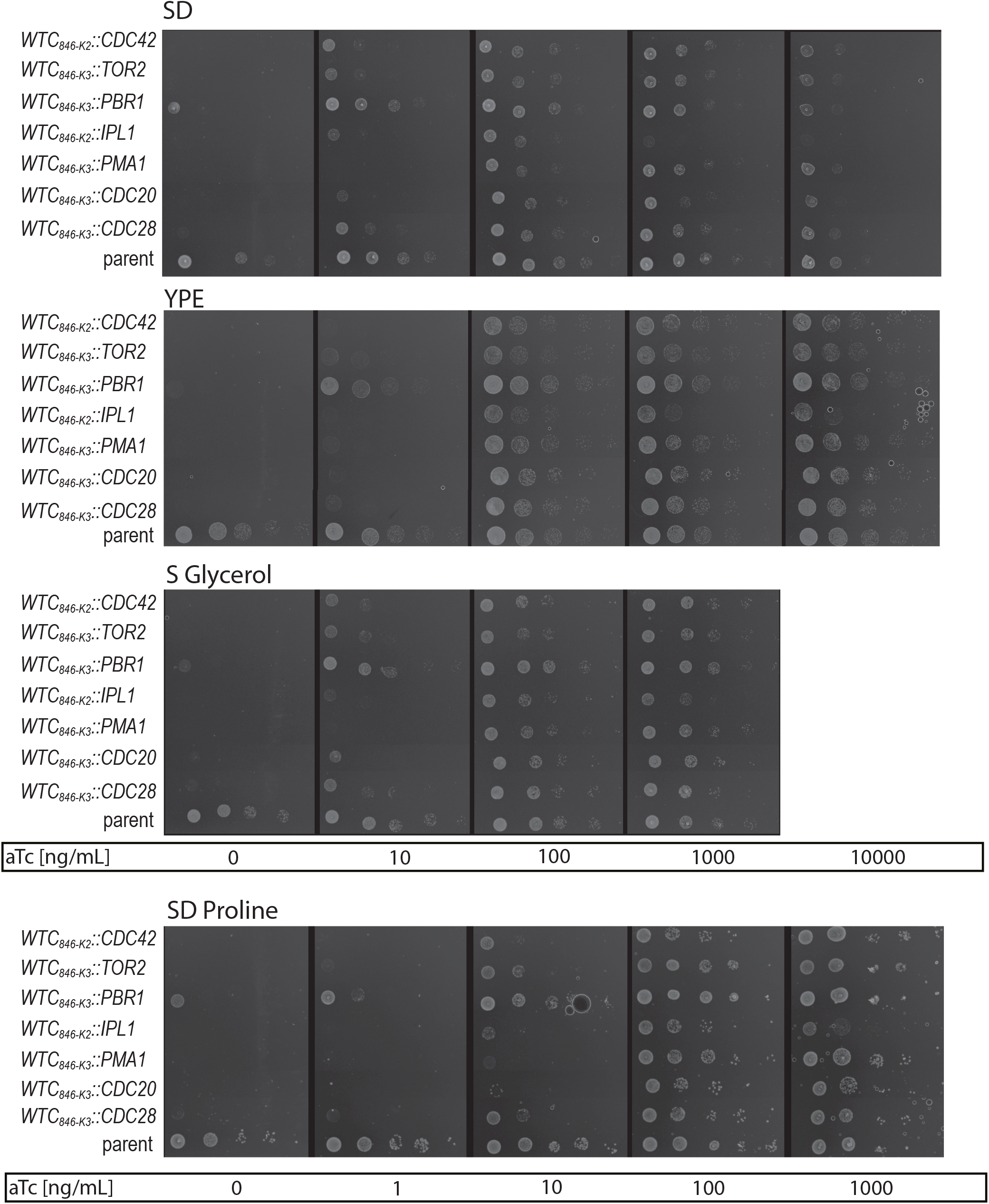
Regulated protein dosage from WTC_846_ alleles controls growth on different solid media. Cells of MATa haploid strains bearing genes whose expression was controlled by WTC_846_ were spotted onto solid media. Names of the genes are given on the left, which correspond to the “Name” column in Table S1. Spotting protocol is explained in Materials&Methods. Briefly, cells were grown in liquid medium that supported growth (YPD with aTc), followed by media allowing WTC_846_ shutoff (YPD without aTc). Cells were grown in YPD without aTc for 6 hours to allow WTC_846_ to fully shut off and any residual gene products to be degraded or diluted. Cells were then spotted onto different plates, such that the leftmost spot on each plate had 2.25×10^6^ cells and each subsequent spot was a 1:10 dilution. The aTc concentration in each plate is indicated below each image. “Parent” refers to strain FRY2769, which only has the *P_7tet.1_* directed TetR and *P_RNR2_* directed TetR-nls-Tup1 expression from the LEU2 locus. Plates were imaged after 24h for SD, and 42h for YPE, S Glycerol and SD Proline. For all but one strain, no growth (as assayed both by increase in optical density of cells in spots and formation of single colonies) was observed without induction. That exception was the WTC_846-K3_::PBR1 strain, which showed slight increase in optical density of the most concentrated spot, indicating very slow growth or residual protein activity. This could be either due to slow degradation kinetics of the protein upon aTc removal, or very low, residual expression even without induction. No single colonies were observed. The fact that strains bearing WTC_846_ controlled essential genes did not grow without aTc confirms that in the absence of the inducer, these WTC_846_ alleles are operationally nulls. In all media tested, growth as assayed by increase in density of the spot and increased frequency of formation of single colonies increased with increasing aTc concentration, indicating titratability of the protein dosage in these alleles. In one strain (e.g. *WTC_846-K2_::IPL1*) decreased growth at high protein dosage was observed as indicated by lighter spots and reduced single colony formation.

**Figure 17:**
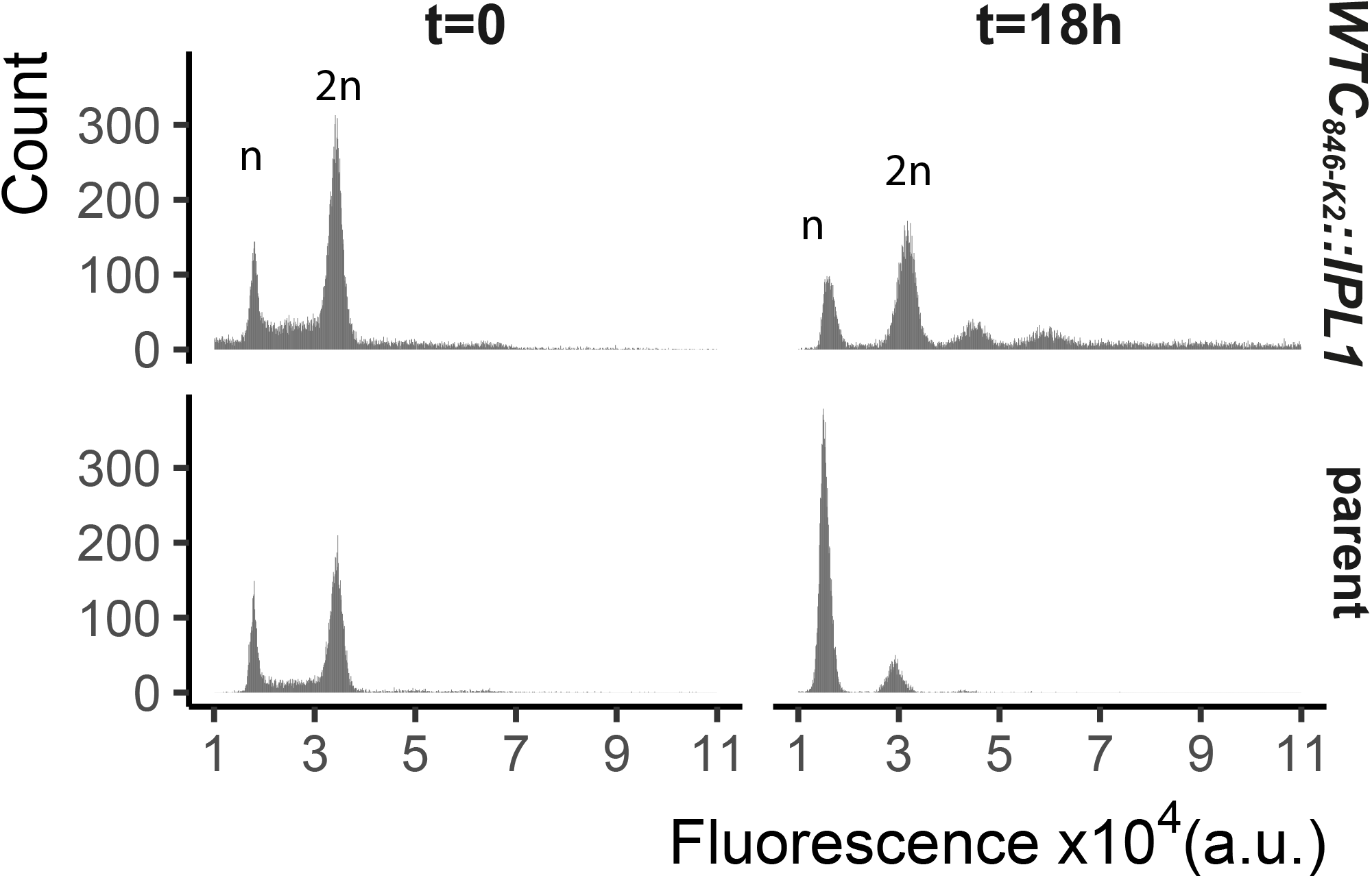
WTC_846_-driven overexpression of Ipl1 prolongs G2/M and produces cells with ¿2n ploidy. We grew FRY2789 *WTC_846-K2_::IPL1* and an otherwise isogenic parent strain (FRY2769), where Ipl1 was under the control of its native promoter, in YPD with 400 ng/mL aTc for 18 hours. At each time point cells were fixed with 70% ethanol, stained for DNA content with Sytox and measured using flow cytometry. We recorded 20000 *WTC_846-K2_::IPL1* cells and 10000 parent cells at each time point. Plots show counts of cells for each fluorescence value. Peaks corresponding to 1 set and 2 sets of chromosomes, indicating cells in G1 and in G2/M are labelled n and 2n. Results indicate that overexpression of Ipl1 leads to a prolonged G2/M phase and cells with aberrant chromosome numbers above 2n.

**Figure 18:**
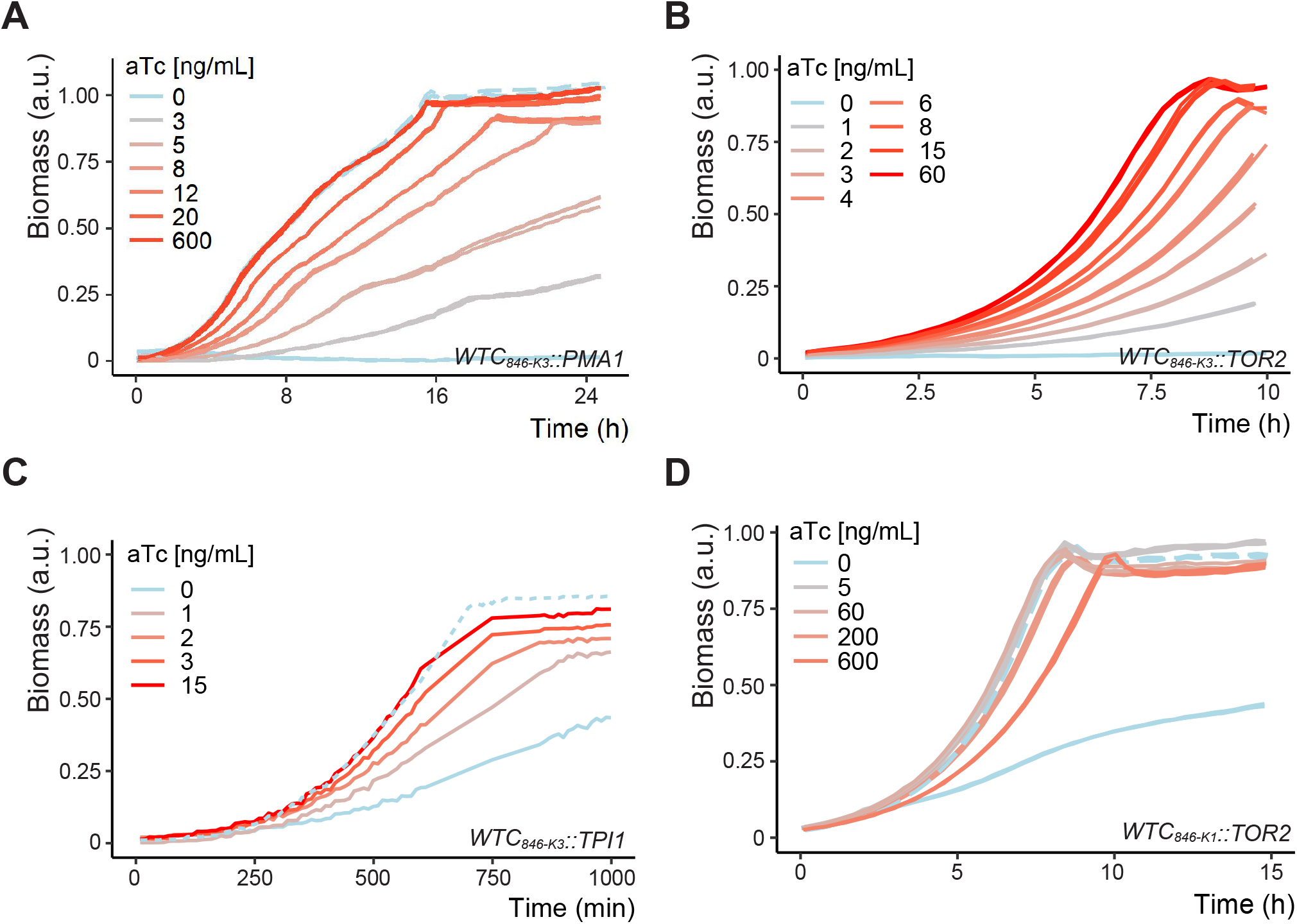
Adjustable protein dosage from WTC_846_ alleles of essential and metabolic genes controls growth rates in different liquid media. Plots show growth curves of (A) FRY2828/ *WTC_846-K3_::PMA1* in SD Full, pH 4.5, (B) FRY2773/*WTC_846-K3_::TOR2* in YPD medium (in triplicate), (C) FRY2849/*WTC_846-K3_::TPI1* in YPD, and (D) FRY2772/*WTC_846-K1_::TOR2* strain in YPD (in duplicates). Solid lines indicate the WTC_846_ alleles, dashed lines, where present, indicate the otherwise isogenic comparison strain in which the gene of interest was under endogenous control (FRY2769). Data in (A,B and D) was acquired with a Biolector device, and 1 mL total culture volume. Data in (C) was acquired with a Growth Profiler and 250uL total culture volume as described in Materials&Methods. Both devices were seeded with 50,000 cells per mL at the start of the experiment. y axis indicates the culture density measured by the device, normalized such that the highest recorded value equals 1. Growth at different aTc concentrations and different dosages of these essential (Pma1,Tor2) and non-essential metabolic (Tpi1) protein products leads to different growth rates. Comparison of panel D FRY2772/*WTC_846-K1_::TOR2* and panel B FRY2773/*WTC_846-K3_::TOR2* demonstrates that the range of adjustable protein dosages achievable by WTC_846_ alleles can be further enlarged by choice of Kozak sequences (translation initiation sequences), and that, in the absence of inducer, the reduced translation *WTC_846-K3_::TOR2* allele is a null.

**Figure 19:**
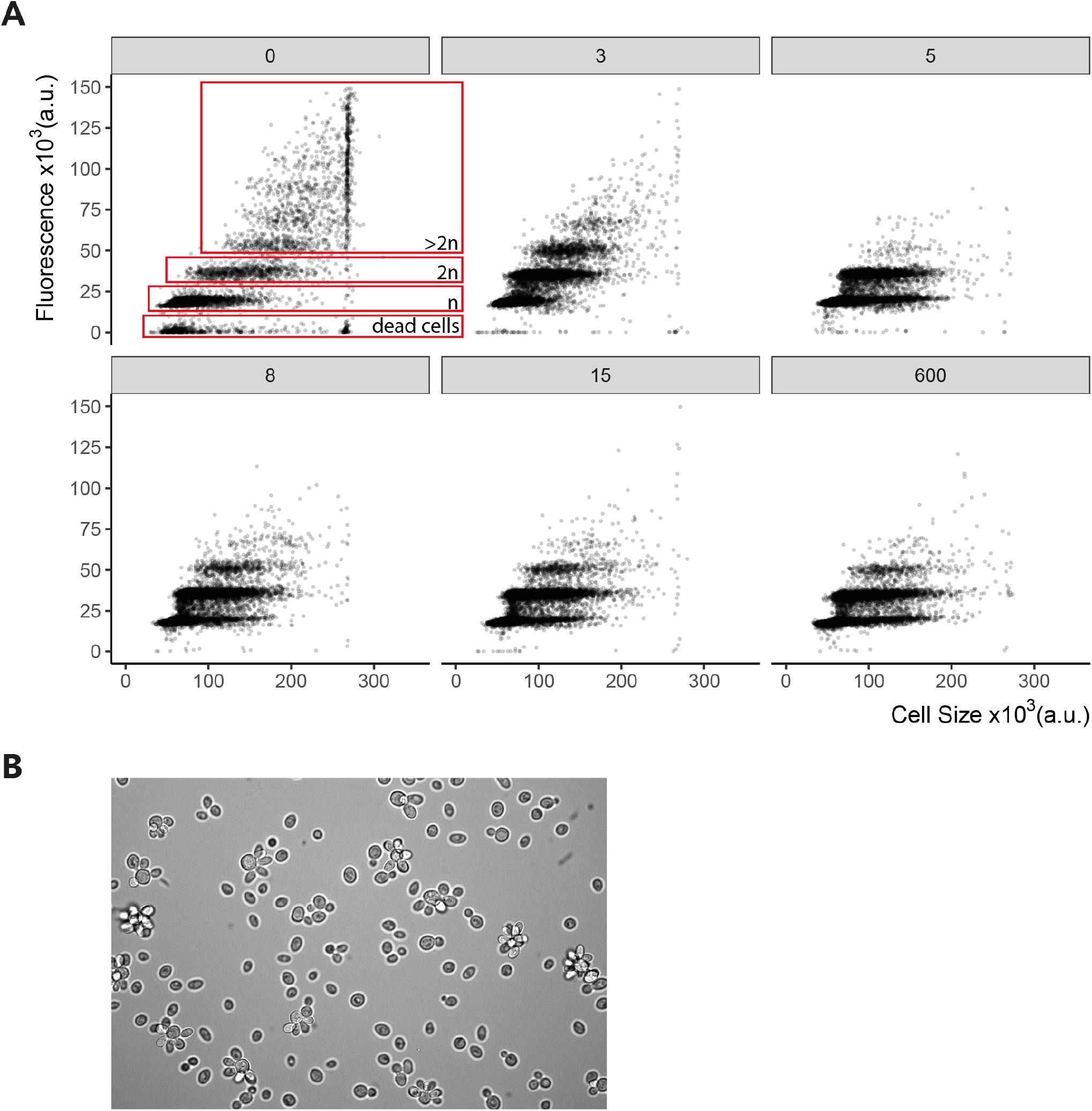
Regulated clamped hypomorphic expression of *WTC_846_::PMA1* allele causes cell separation defect. We grew FRY2828/*WTC_846-K3_::PMA1* in SD Full media with the aTc concentrations indicated as ng/mL in the grey boxes above the panels. We fixed the samples with 70% ethanol for Sytox staining. We used flow cytometry to analyze the cells and fluorescence was used as a proxy for DNA content. Figure shows peaks in fluorescence corresponding to 1 (n) and 2 (2n) sets of chromosomes indicating cells in G1 and G2/M. Figure 23 also shows peaks corresponding to cells with ¿2 sets of chromosomes, and dead cells/debris. We calculated cell volume as the vector of SSC-H and FSC-W signals as explained in Materials&Methods. Each dot corresponds to a single cell measurement. Values above 280000 saturated the measurement device, and this fact resulted in an apparent increase in the number of cells around this value. At no or low aTc concentrations and thus low Pma1 abundance, daughter cells fail to separate from the mother, leading to an apparent increase in cell volume and ploidy as shown in panel (B) using microscopic observation of a representative sample of *WTC_846-K3_::PMA1* cells grown without aTc. Image was acquired using 40x magnification.

**Figure 20:**
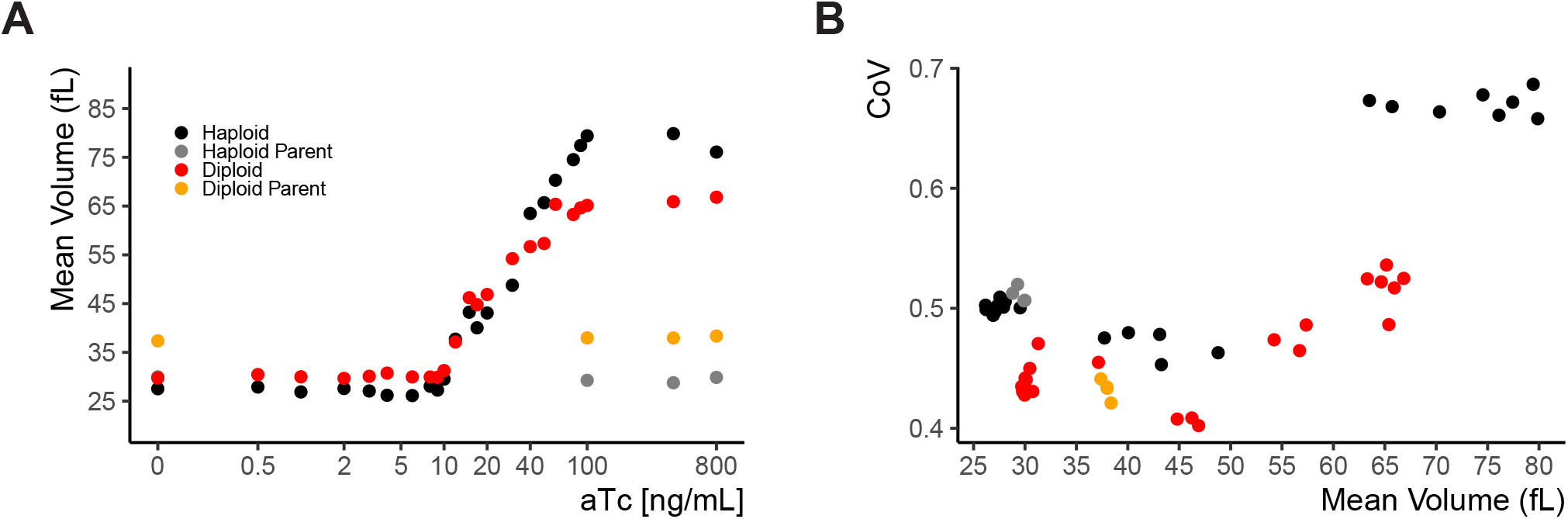
Whi5 titration leads to increased cell volume without an increase in cell-to-cell variation. A) Expression of Whi5 was clamped at different levels by growth of FRY2791, a haploid *WTC_846-K1_::WHI5* strain, of FRY2929, a *WTC_846-K1_::WHI5/WTC_846-K1_::WHI5* diploid strain, and otherwise-isogenic control strains in which Whi5 was expressed from its endogenous promoter (FRY2769). Cells were grown in S Ethanol medium at different concentrations of aTc to yield different cell volumes, measured by a Coulter counter. B) We calculated and plotted the CoV of the mean cell volume at each aTc dose as a measure of CCV. Except when Whi5 is overexpressed in haploid cells, CoV of this WTC_846_ controlled phenotype is at or around the same level as WT variation.

**Figure 21:**
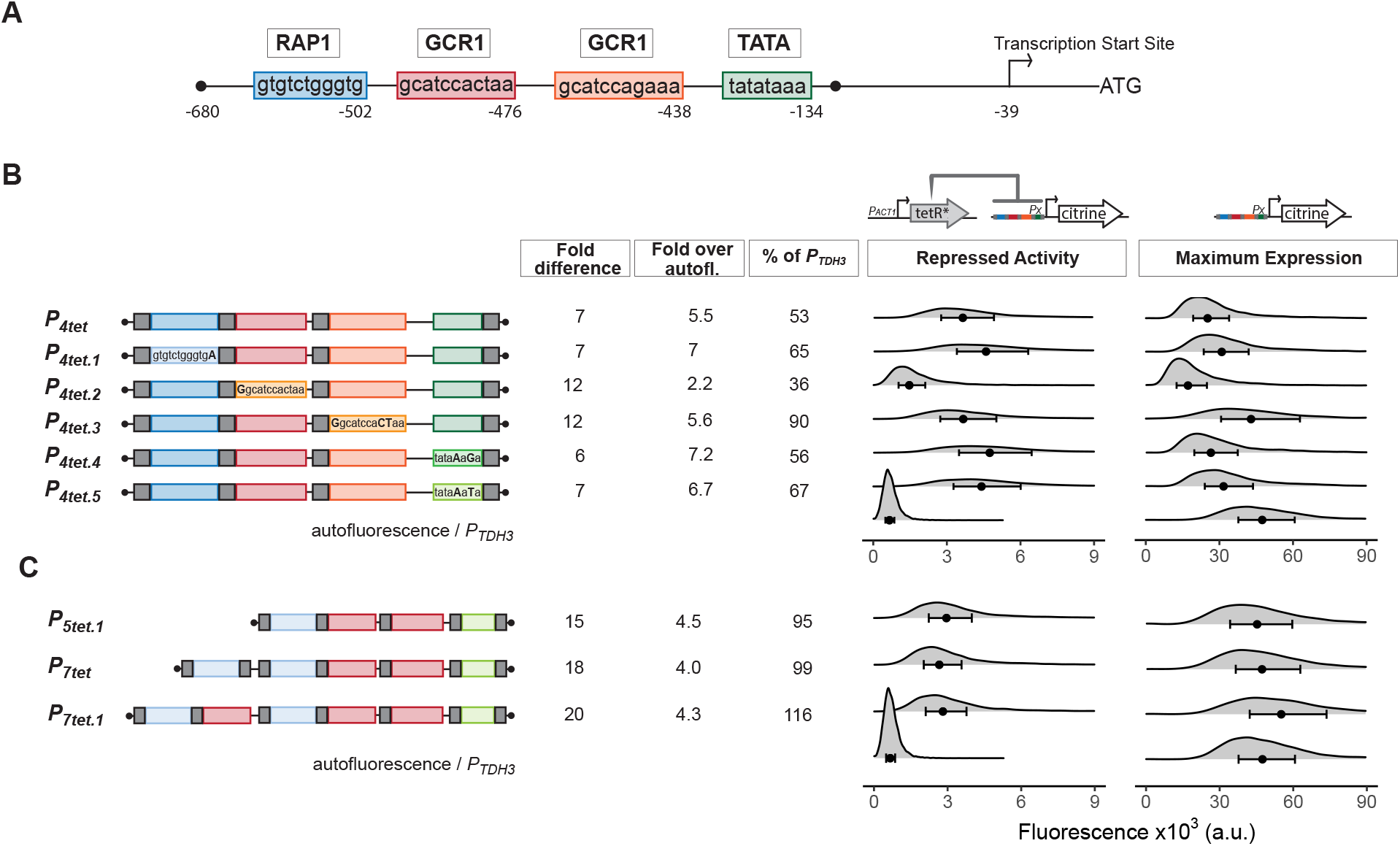
Optimization of *tetO_1_* placements and endogenous transcription factor binding sites to increase maximum activity of a TetR-repressible derivative of *P_TDH3_*. (A) Diagram of *P_TDH3_* shows the nucleotide positions of the binding sites for the endogenous transcription factors Rap1 and Gcr1, the TATA-sequence, and the transcription start site relative to the TDH3 start codon. (B) Repression and maximum activity of the *P_TDH3_* derivatives tested for optimization. Diagrams above the plots display the genetic elements of strains used (FRY2565, 2575,2598,2647,2599,2648,2601,2649,2602,2650,2603,2656,70,2683). Left diagram depicts strains used to test repressed activity, right diagram maximum activity. Px denotes any tetR repressible promoter. The * in TetR indicates a SV40 Nuclear Localization Sequence. In all strains, the *P_TDH3_* derivative promoters diagrammed on the left directed the synthesis of Citrine integrated into the LEU2 locus. Grey boxes inside the diagrams denote *tetO_1_* TetR binding sites. For measurement of repressed activity, otherwise-isogenic strains carried a *P_ACT1_*-TetR construct integrated in the HIS3 locus. Citrine fluorescent signal was detected by flow cytometry. For the measurements, “fold difference” measures the median of the maximum activity signal divided by the median of the repressed activity. “Fold over autofluorescence” refers to median repressed activity signal divided by the median autofluorescence background signal. Maximum promoter activity is quantified as median fluorescence signal expressed as percentage of signal from otherwise-isogenic *P_TDH3_*-Citrine strain. For the plots, x axis shows intensity of fluorescence signal. Plots are density distributions of the whole population, such that the area under the curve equals 1 and the y axis indicates the proportion of cells at each fluorescence value. The circles inside each density plot show the median and the upper and lower bounds of the bar show the first and third quartiles of the distribution. C) Repression and maximum activity of optimized *P_5tet_* derivatives. Diagrams and plots as in (B). These promoter variants contained additional binding sites for Rap1 and Gcr1 selected for higher activity, as well as an alternative TATA sequence as described.

**Figure 22:**
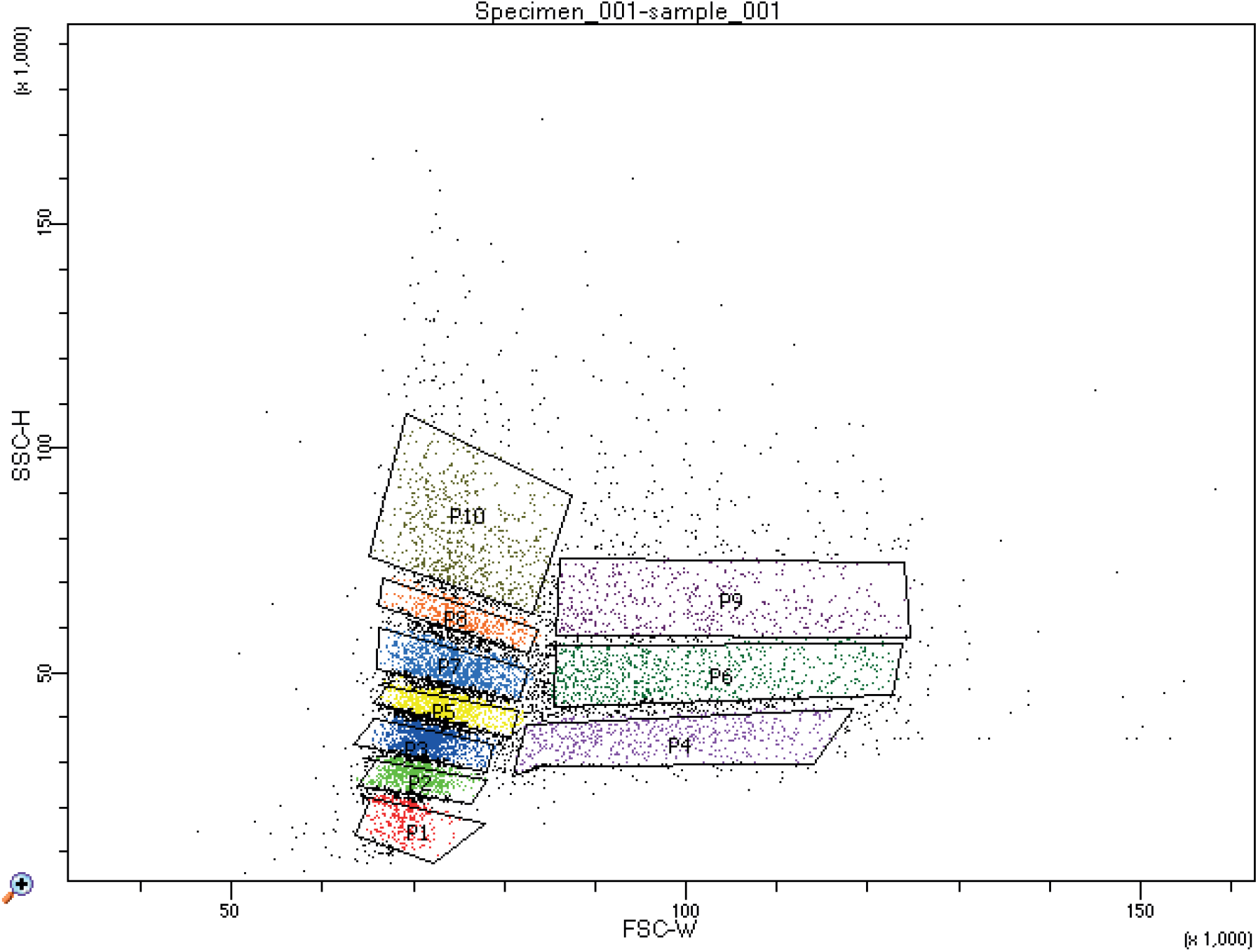
Sub-populations collected by FACS for validating the cell volume proxy measure. Strain FRY2683 was grown to exponential phase in YPD and was run through the sorter at a concentration of 2 million cells per mL. 10 separate gates were set on the FSC-W and SSC-H signals for collecting sub-populations as depicted in the figure.

**Figure 23:**
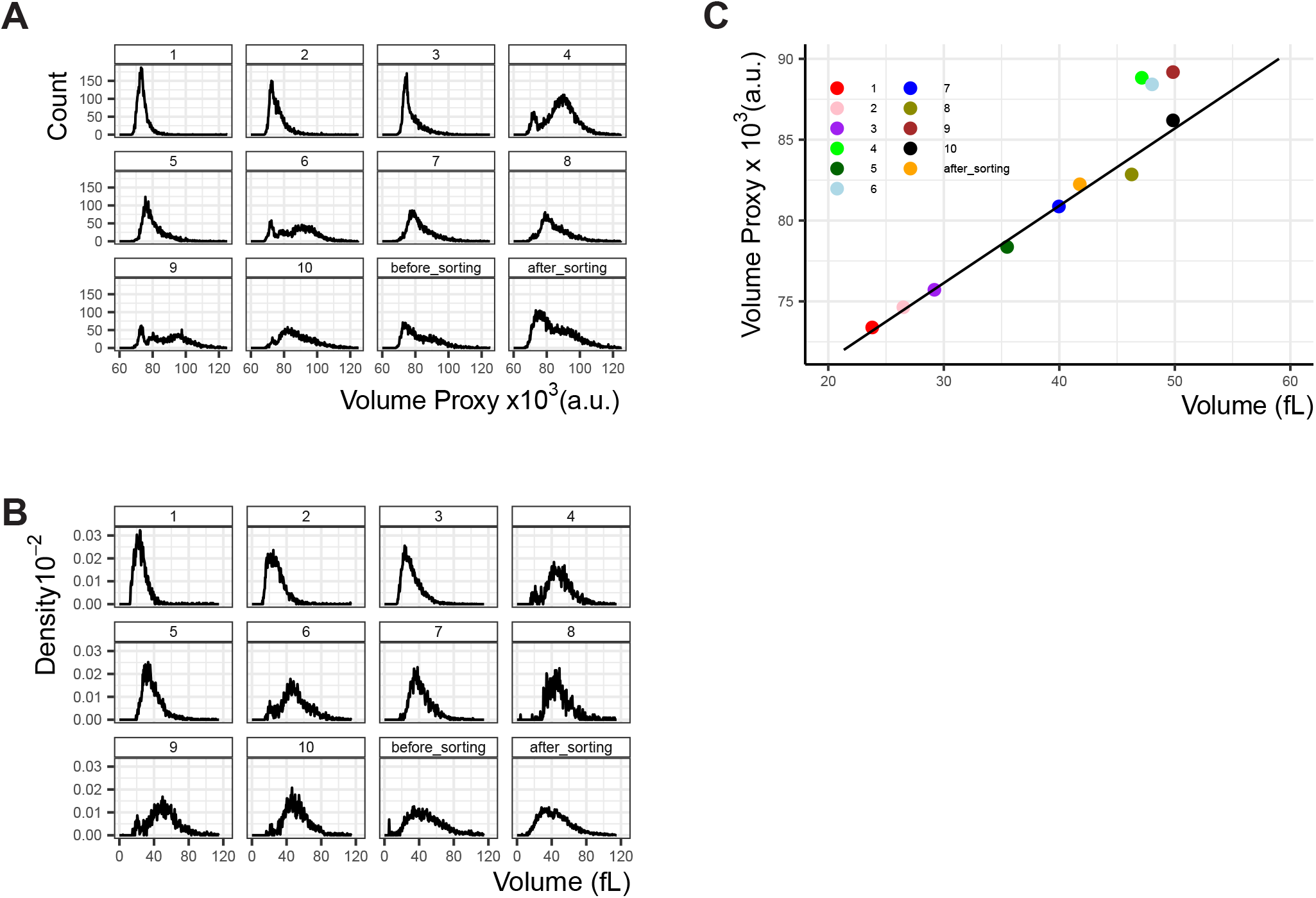
Comparison of the flow cytometry based cell volume proxy to cell volume measured by Coulter counter. 10 sub-populations were collected from an exponentially growing culture used as constitutive Citrine expression control in other experiments (FRY2683), using FACS as described in Figure S22. Each of the sub-populations were (A) measured in the LSRII Fortessa LSR flow cytometer used for all other experiments and the proxy for cell volume calculated and (B) measured in a Coulter counter. The sub-population numbers are indicated above the plots and correspond to the gates seen in Figure S22. The culture was kept on ice throughout the sorting process, and the full population was measured twice, once before the sorting process began (before sorting) and once after (after sorting) to ensure the volume distribution in the population did not change over the course of the experiment. No significant difference was observed. (C) The median volume of each sub-population was plotted, as measured by flow cytometry (y axis) or Coulter counter (x axis). The linear fit was generated as explained in Materials&Methods (R^2^=0.98, p=7.37×10^−7^), without taking into account gates 4,6, and 9 where the distribution is bimodal and the median is not a good descriptor of the population.(A) displays cell counts per volume proxy, and B displays density plots where the area under the curve is 1.

**Figure 24:**
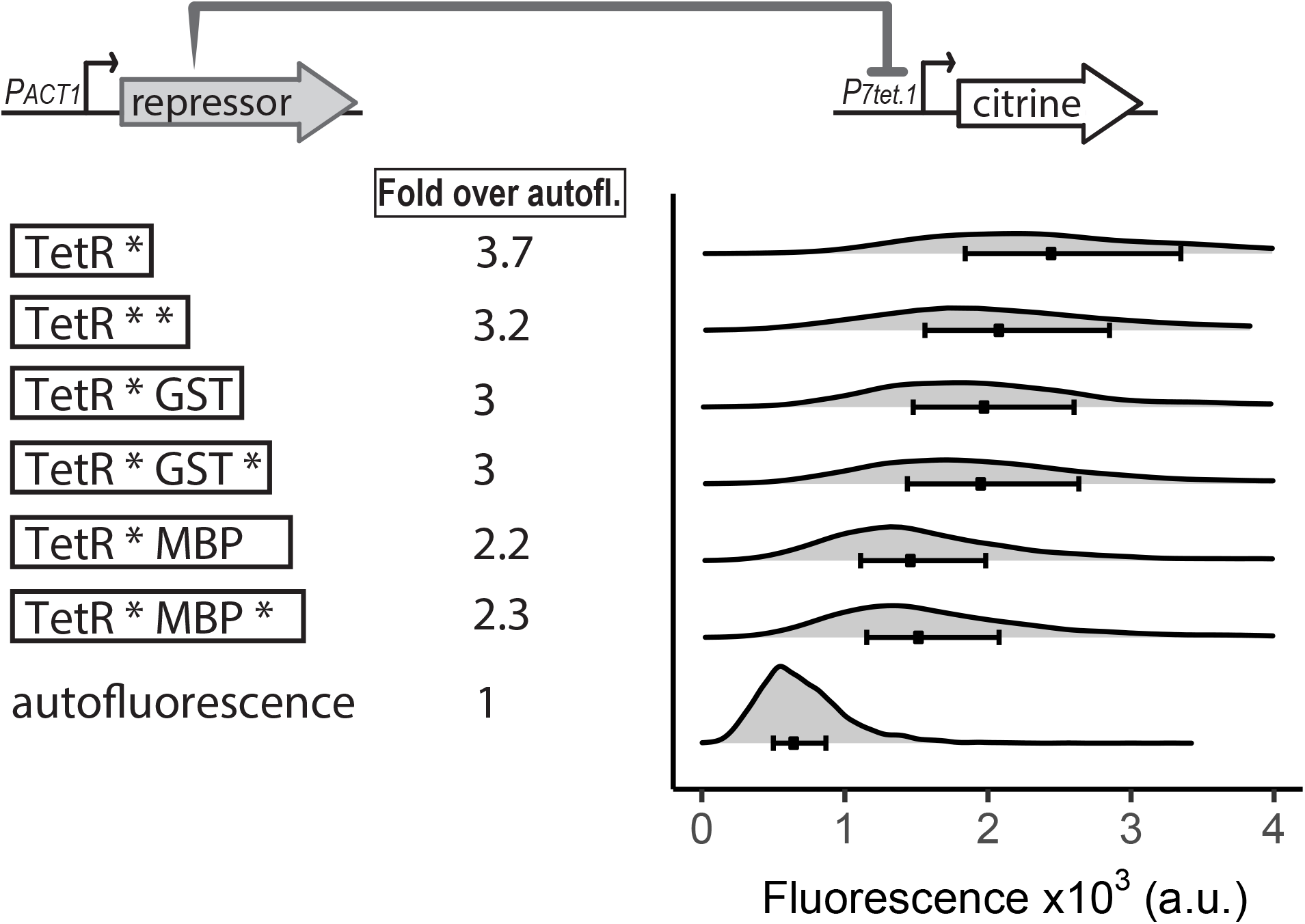
Effect of increased nuclear concentration and size of TetR on repression. The top diagram indicates the genetic elements of the SR architecture used to test the ability of various TetR derivatives to abolish basal activity of *P_7tet.1_*. Diagrams to the left of the plots show the different repressors used. Each * indicates one SV40 Nuclear Localization Sequence. GST refers to Glutathione S-transferase, and MBP to Maltose Binding Protein, both of *E.coli*. Citrine fluorescence from *P_7tet.1_* repressed by the repressors indicated was measured using flow cytometry. Plots are density distributions of the whole population, such that the area under the curve equals 1 and the y axis indicates the proportion of cells at each fluorescence value. The circles inside each density plot show the median and the upper and lower bounds of the bar correspond to the first and third quartiles of the distribution. Numbers to the left of the plot indicate fold expression over autofluorescence, i.e. the median of the Citrine fluorescence detected divided by the median of the autofluorescence signal. Although increased nuclear concentration and size of TetR increase repression efficiency, these strategies are not enough to fully abolish basal expression from *P_7tet.1_*.

---

**Table S1:**
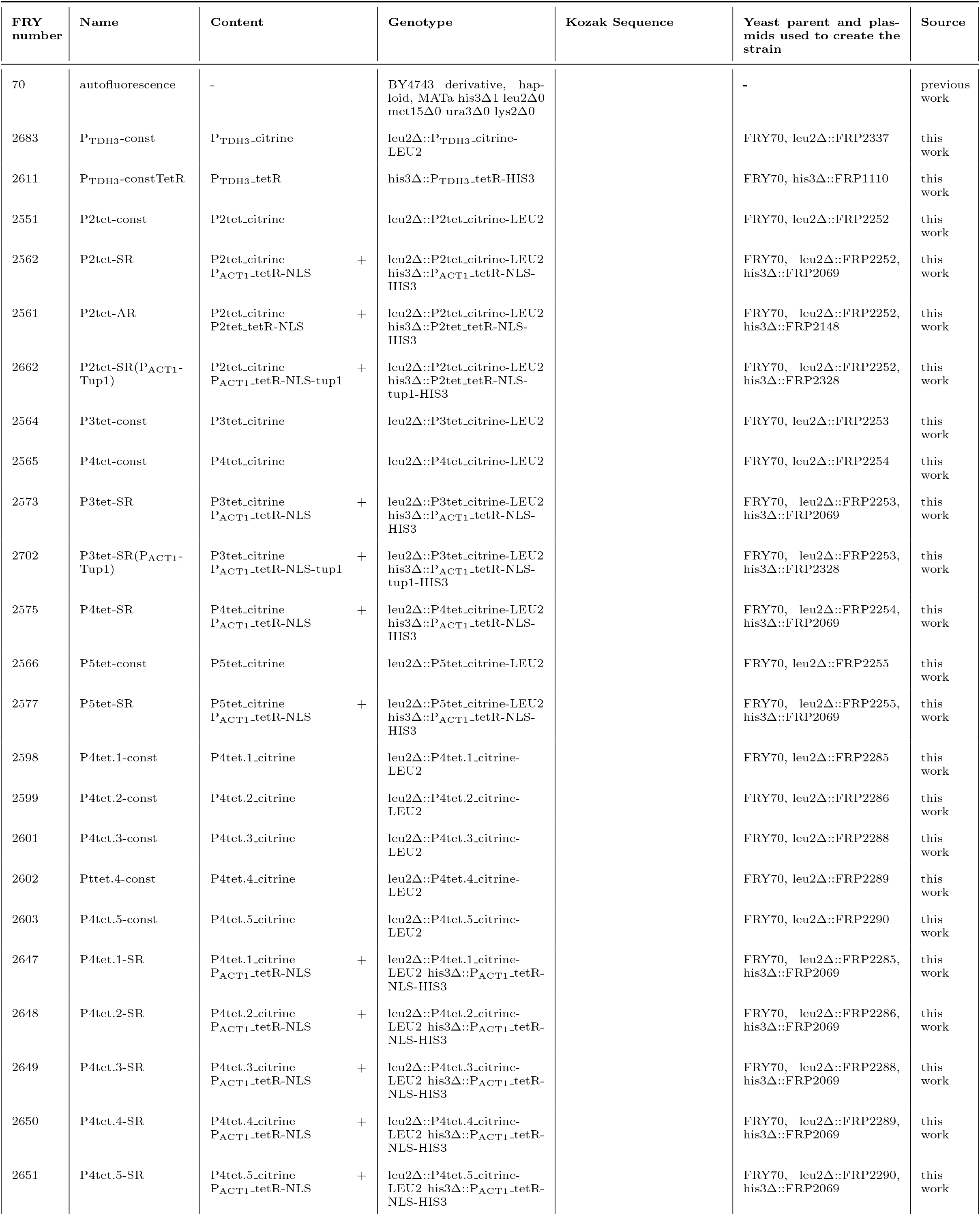
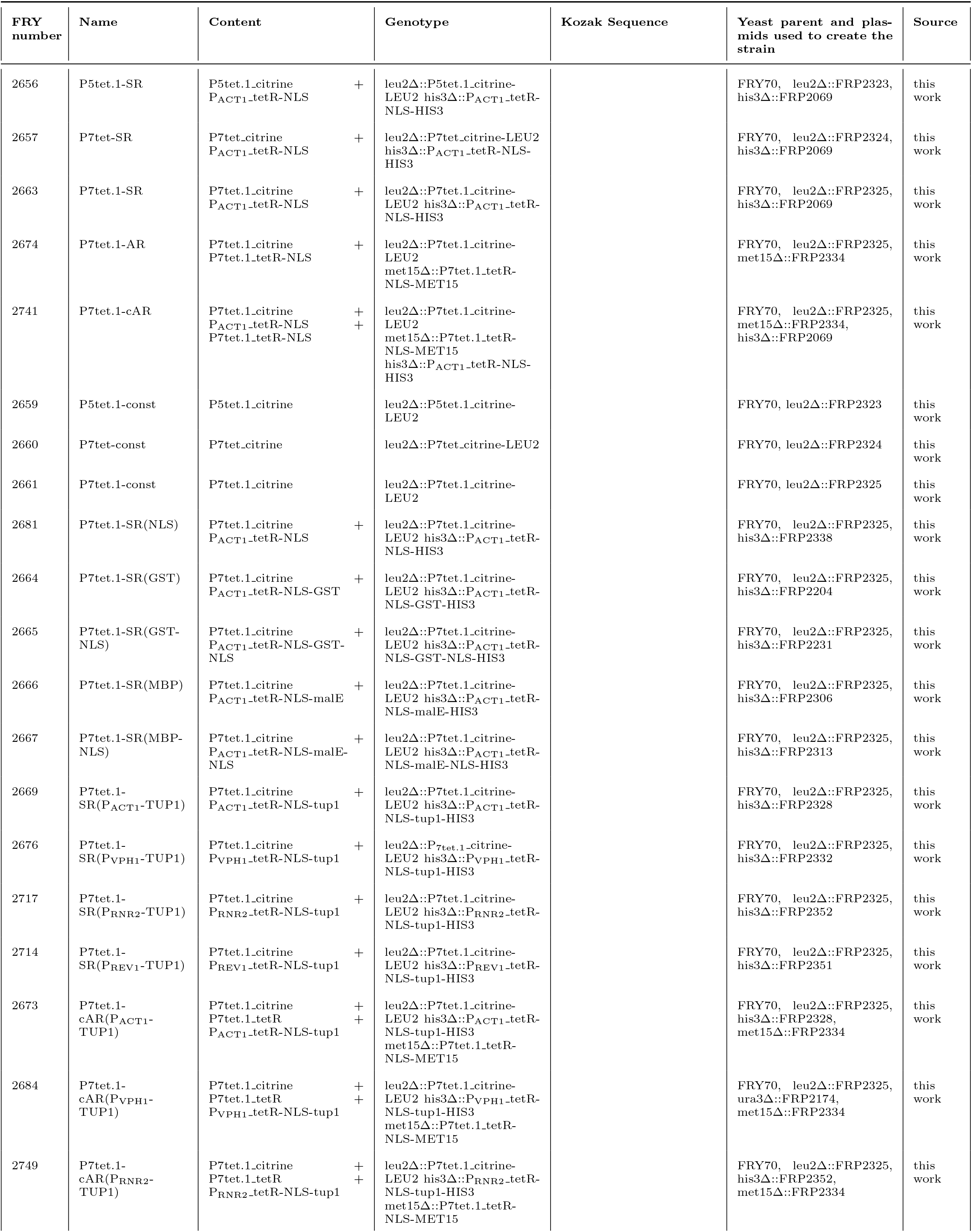
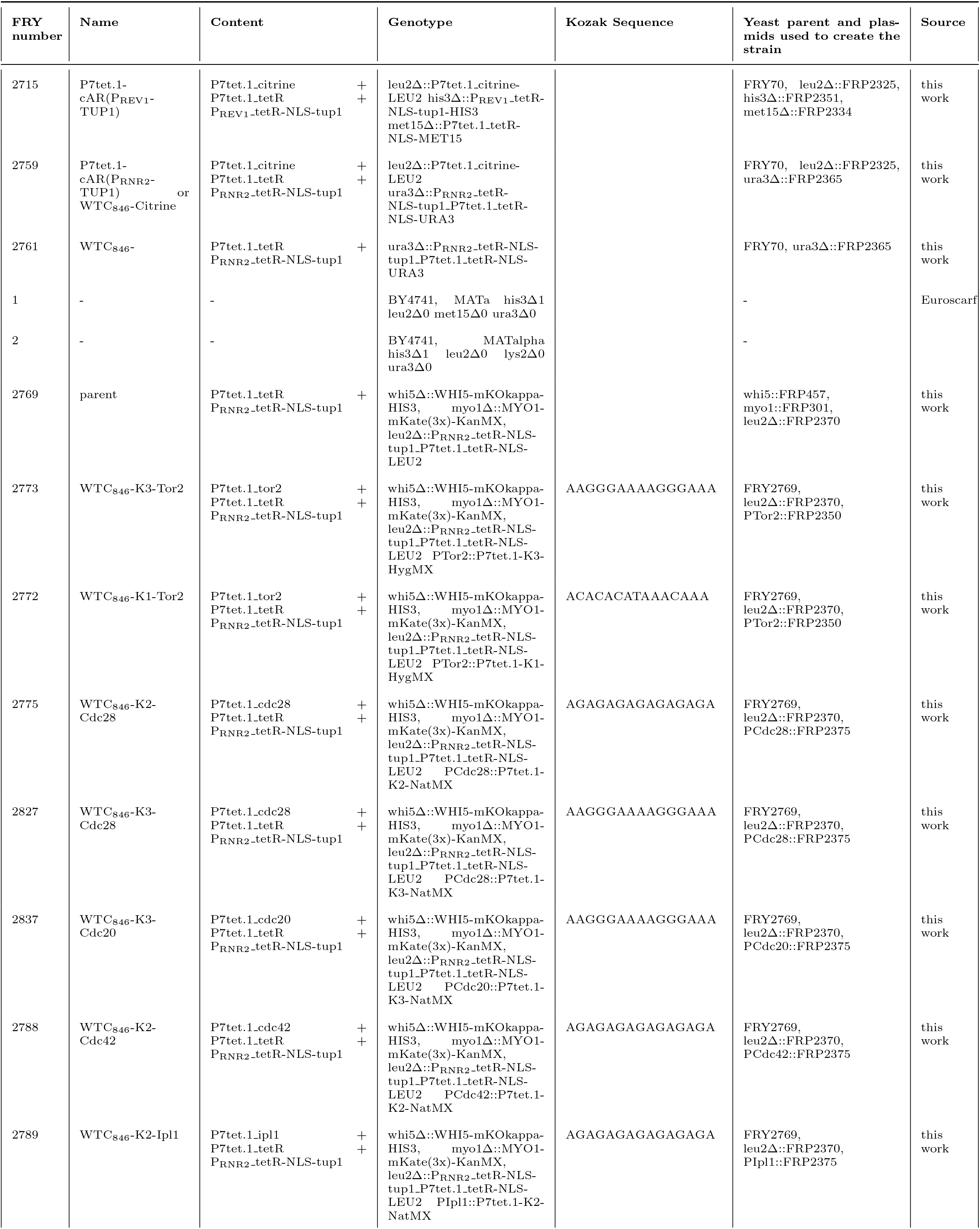
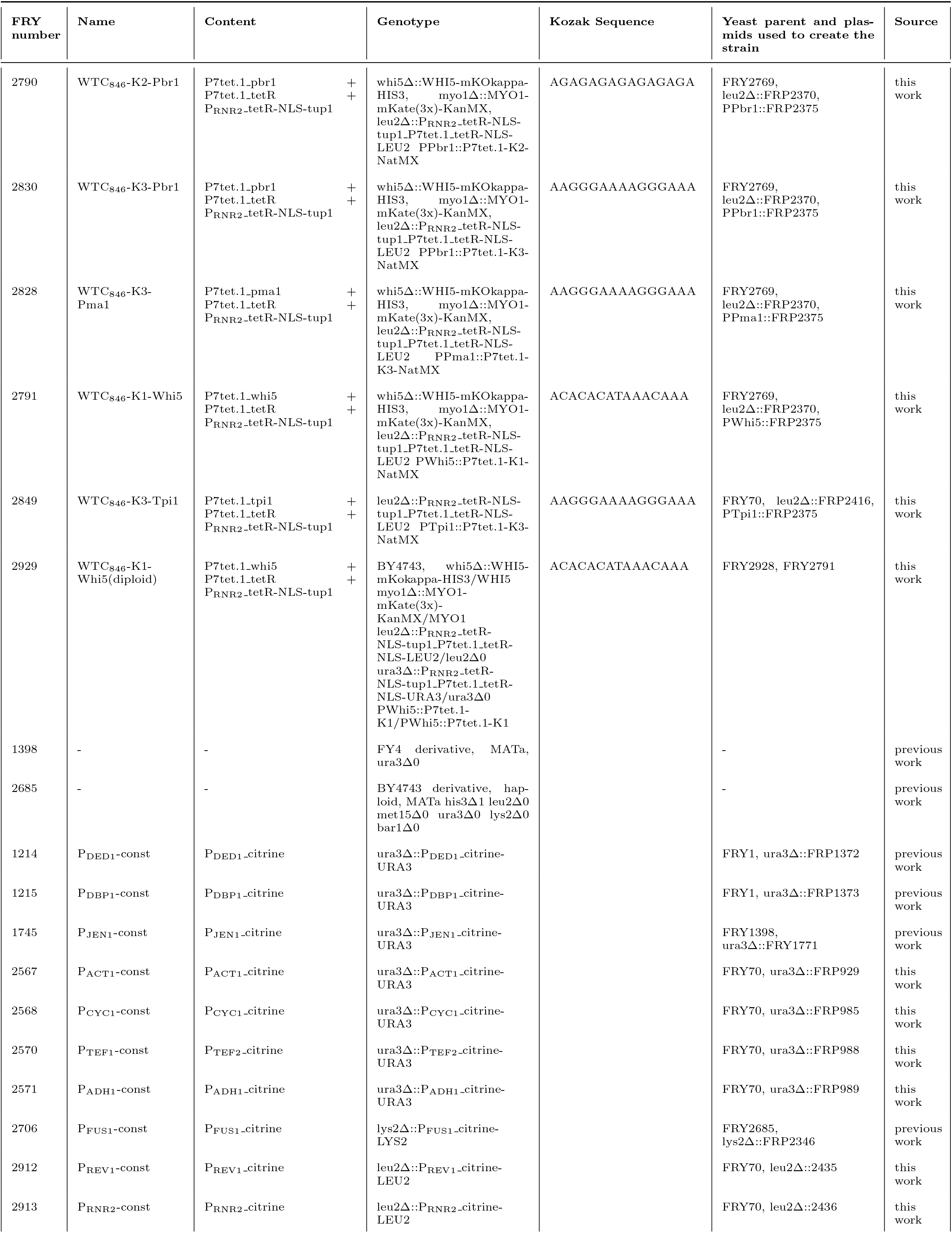
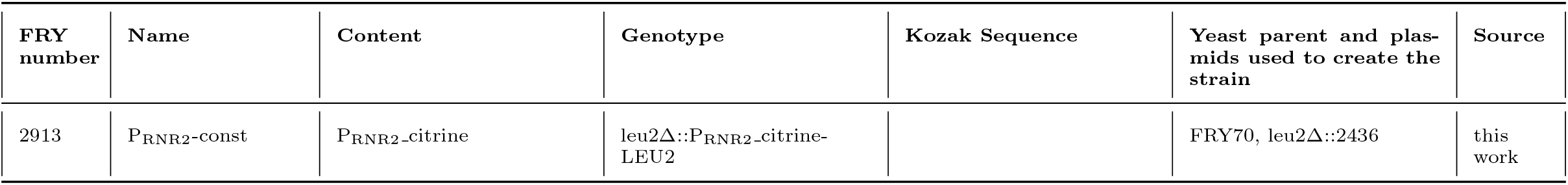
Strains used in this study. Kozak sequence is the last 15bp before the start codon.

**Table S2:** Plasmids used in this study. * indicates plasmids available through Addgene. † indicates that the plasmid has the insert in the reverse orientation compared to all the other plasmids containing Citrine (See Materials&Methods).

| FRP number | Content | Yeast Marker | Source |
| --- | --- | --- | --- |
| 2337 | P<sub>TDH3</sub>-citrine.t<sub>CYC1</sub> | Leu2MX | this work |
| 2088† | p2tet.citrine.t<sub>CYC1</sub> | Leu2MX | this work |
| 2252 | P2tet.citrine.t<sub>CYC1</sub> | Leu2MX | this work |
| 2069 | P<sub>ACT1</sub>-tetR-nls.t<sub>CYC1</sub> | His3MX | this work |
| 2174 | P<sub>VPH1</sub>-tetR-nls.t<sub>CYC1</sub> | Ura3 | this work |
| 2148 | P2tet.tetR-nls.t<sub>CYC1</sub> | His3MX | this work |
| 2334 | P7tet.tetR-nls.t<sub>CYC1</sub> | Met15 | this work |
| 2253 | P3tet.citrine.t<sub>CYC1</sub> | Leu2MX | this work |
| 2254 | P4tet.citrine.t<sub>CYC1</sub> | Leu2MX | this work |
| 2255 | P5tet.citrine.t<sub>CYC1</sub> | Leu2MX | this work |
| 2285 | P4tet.1.citrine.t<sub>CYC1</sub> | Leu2MX | this work |
| 2286 | P4tet.2.citrine.t<sub>CYC1</sub> | Leu2MX | this work |
| 2288 | P4tet.3.citrine.t<sub>CYC1</sub> | Leu2MX | this work |
| 2289 | P4tet.4.citrine.t<sub>CYC1</sub> | Leu2MX | this work |
| 2290 | P4tet.5.citrine.t<sub>CYC1</sub> | Leu2MX | this work |
| 2323 | P5tet.1.citrine.t<sub>CYC1</sub> | Leu2MX | this work |
| 2324 | P7tet.citrine.t<sub>CYC1</sub> | Leu2MX | this work |
| 2325 | P7tet.1.citrine.t<sub>CYC1</sub> | Leu2MX | this work |
| 2338 | P<sub>ACT1</sub>-tetR-nls-nls.t<sub>CYC1</sub> | His3MX | this work |
| 2204 | P<sub>ACT1</sub>-tetR-nls-gst.t<sub>CYC1</sub> | His3MX | this work |
| 2231 | P<sub>ACT1</sub>-tetR-nls-gst-nls.t<sub>CYC1</sub> | His3MX | this work |
| 2306 | P<sub>ACT1</sub>-tetR-nls-malE.t<sub>CYC1</sub> | His3MX | this work |
| 2313 | P<sub>ACT1</sub>-tetR-nls-malE-nls.t<sub>CYC1</sub> | His3MX | this work |
| 2328 | P<sub>ACT1</sub>-tetR-nls-tup1.t<sub>CYC1</sub> | His3MX | this work |
| 2332 | P<sub>VPH1</sub>-tetR-nls-tup1.t<sub>CYC1</sub> | His3MX | this work |
| 2352 | P<sub>RNR2</sub>-tetR-nls-tup1.t<sub>CYC1</sub> | His3MX | this work |
| 2351 | P<sub>REV1</sub>-tetR-nls-tup1.t<sub>CYC1</sub> | His3MX | this work |
| 457 | mKOKappa | His3MX | previous work |
| 301 | mKate | KanMX | previous work |
| 2416 | P<sub>RNR2</sub>-tetR-nls-tup1.t<sub>CYC1</sub>-P7tet.1.tetR-nls.t<sub>CYC1</sub> | Ura3MX | this work |
| 1372 | P<sub>DED1</sub>-citrine.t<sub>CYC1</sub> | Ura3 | previous work |
| 1373 | P<sub>DBP1</sub>-citrine.t<sub>CYC1</sub> | Ura3 | previous work |
| 1771 | P<sub>JEN1</sub>-citrine.t<sub>CYC1</sub> | Ura3 | previous work |
| 929 | P<sub>ACT1</sub>-citrine.t<sub>CYC1</sub> | Ura3 | previous work |
| 985 | P<sub>CYC1</sub>-citrine.t<sub>CYC1</sub> | Ura3 | previous work |
| 988 | P<sub>TEF2</sub>-citrine.t<sub>CYC1</sub> | Ura3 | previous work |
| 989 | P<sub>ADH1</sub>-citrine.t<sub>CYC1</sub> | Ura3 | previous work |
| 2346 | P<sub>FUS1</sub>-citrine.t<sub>CYC1</sub> | Lys2 | previous work |
| 2435 | P<sub>REV1</sub>-citrine.t<sub>CYC1</sub> | Leu2MX | this work |
| 2436 | P<sub>RNR2</sub>-citrine.t<sub>CYC1</sub> | Leu2MX | this work |
| 2365* | P<sub>RNR2</sub>-tetR-nls-tup1.t<sub>CYC1</sub>-P7tet.1.tetR-nls.t<sub>CYC1</sub> | Ura3MX | this work |
| 2370* | P<sub>RNR2</sub>-tetR-nls-tup1.t<sub>CYC1</sub>-P7tet.1.tetR-nls.t<sub>CYC1</sub> | Leu2MX | this work |
| 2350* | P7tet.1 | HygMX | this work |
| 2375* | P7tet.1 | NatMX | this work |
| 2371* | P<sub>RNR2</sub>-tetR-nls-tup1-t<sub>CYC1</sub>-P7tet.1-tetR-nls-t<sub>CYC1</sub> | His3MX | this work |
| 2372* | P<sub>RNR2</sub>-tetR-nls-tup1-t<sub>CYC1</sub>-P7tet.1-tetR-nls-t<sub>CYC1</sub> | Lys2 | this work |
| 2374* | P<sub>RNR2</sub>-tetR-nls-tup1-t<sub>CYC1</sub>-P7tet.1-tetR-nls-t<sub>CYC1</sub> | Met15 | this work |

**Table S3:**
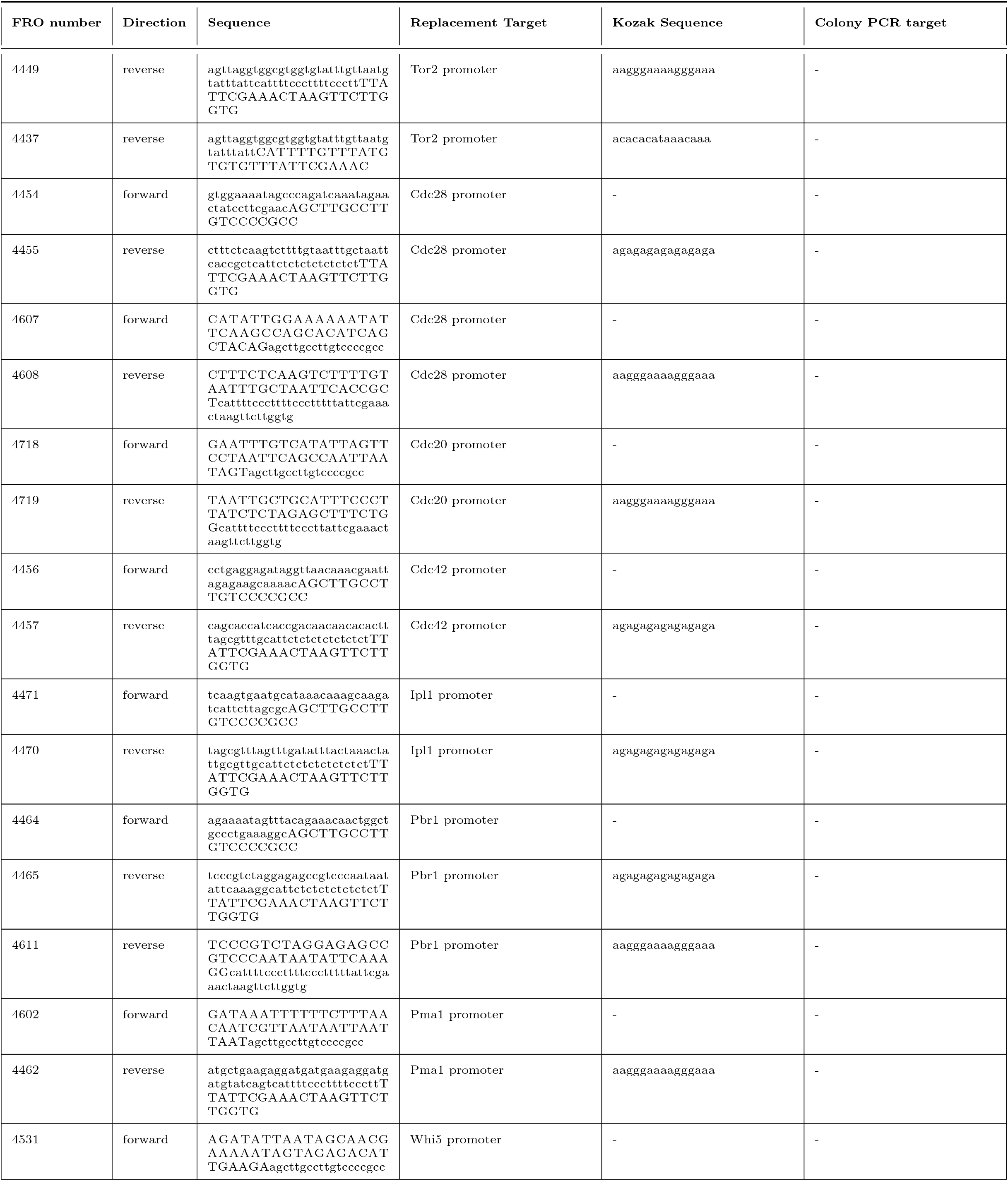
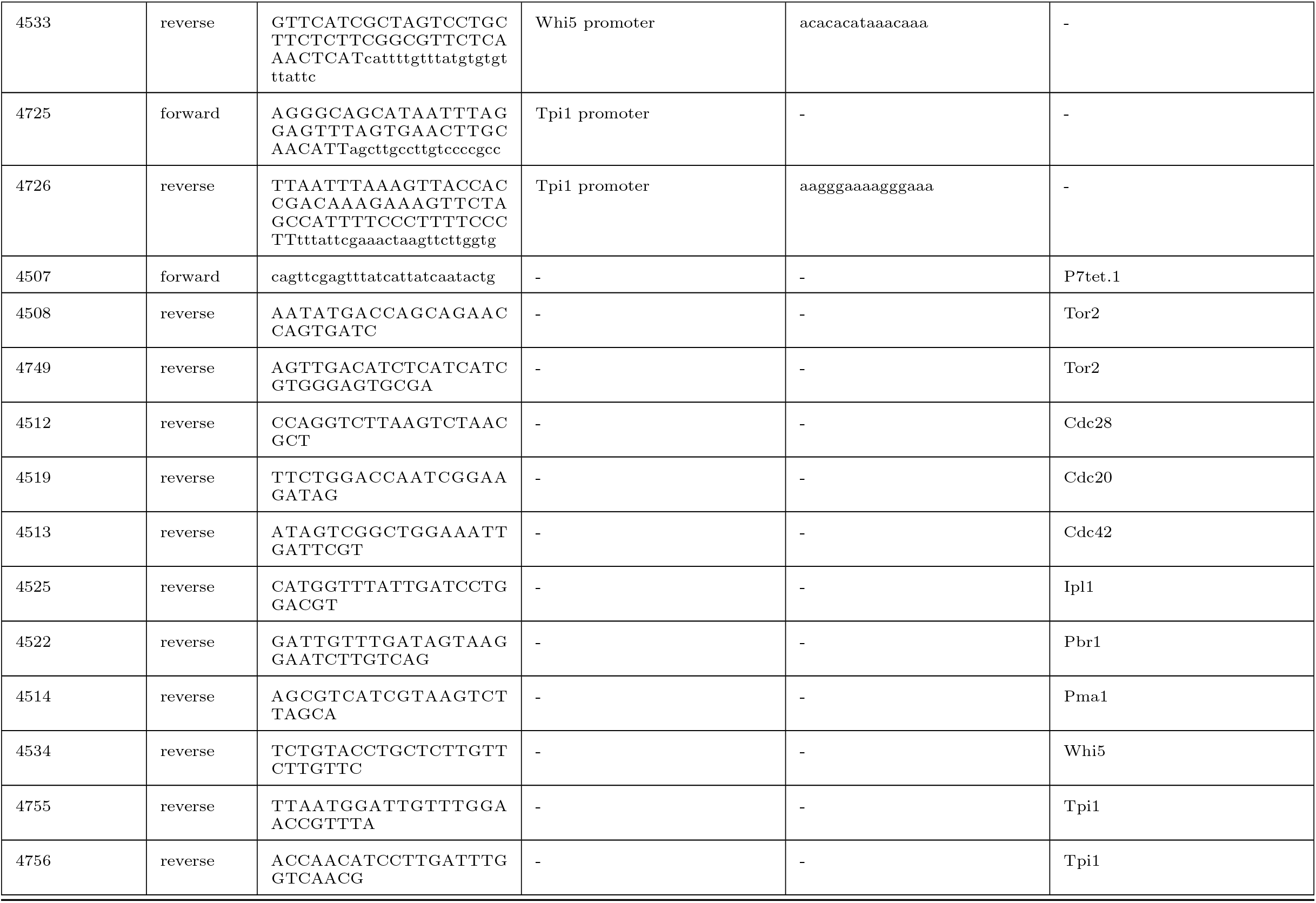
Oligos used in this study to create strains where WTC_846_ controls endogenous gene expression. See Supplementary Information for the protocol used for endogenous gene promoter replacement through homology directed repair. These oligos were used in conjunction with FRP2375 (NatMX) or FRP2350 (HygMX) to create the linear PCR fragment necessary for promoter replacement, or as colony PCR oligos to confirm correct promoter replacement.

**Table S4:**
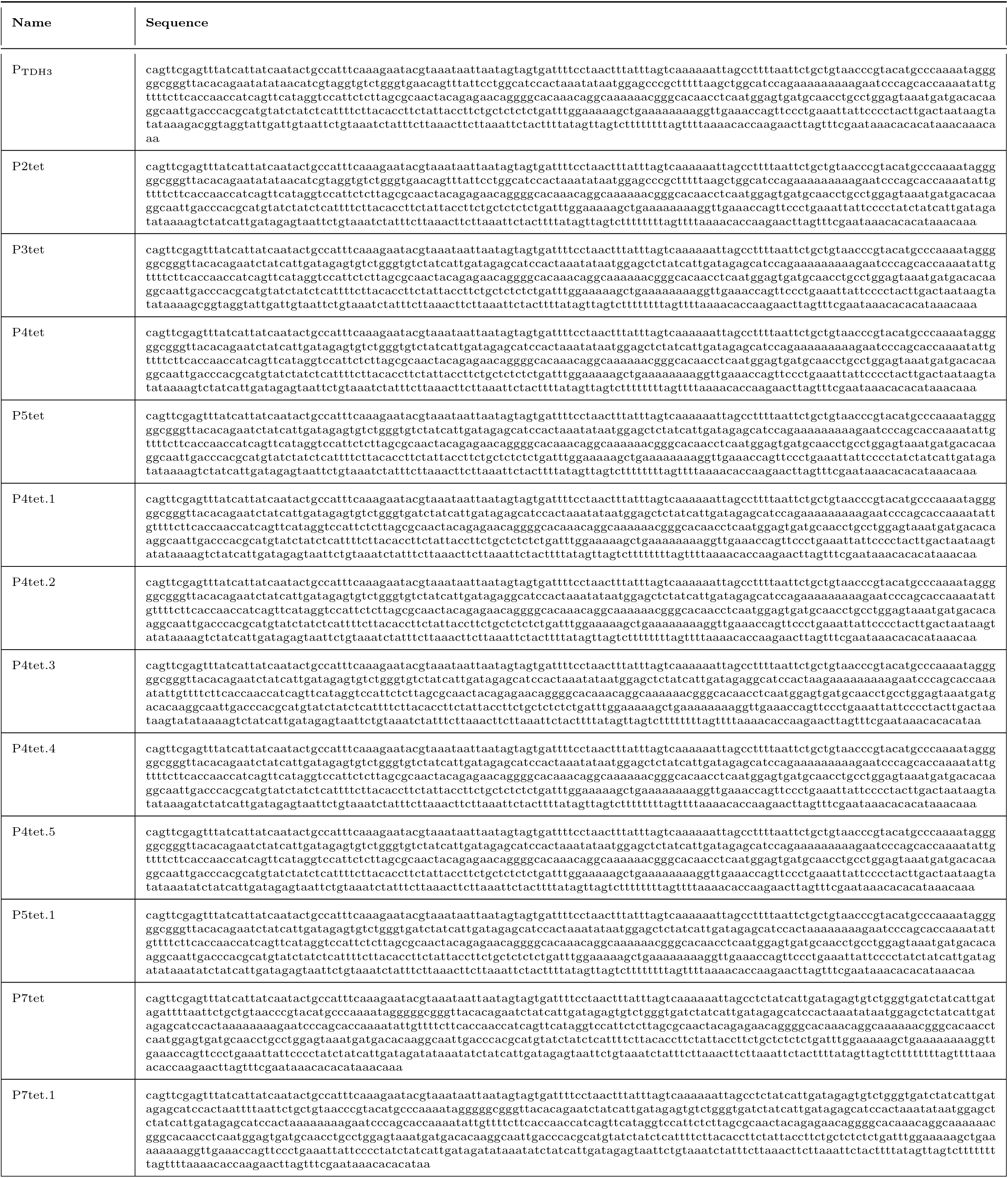
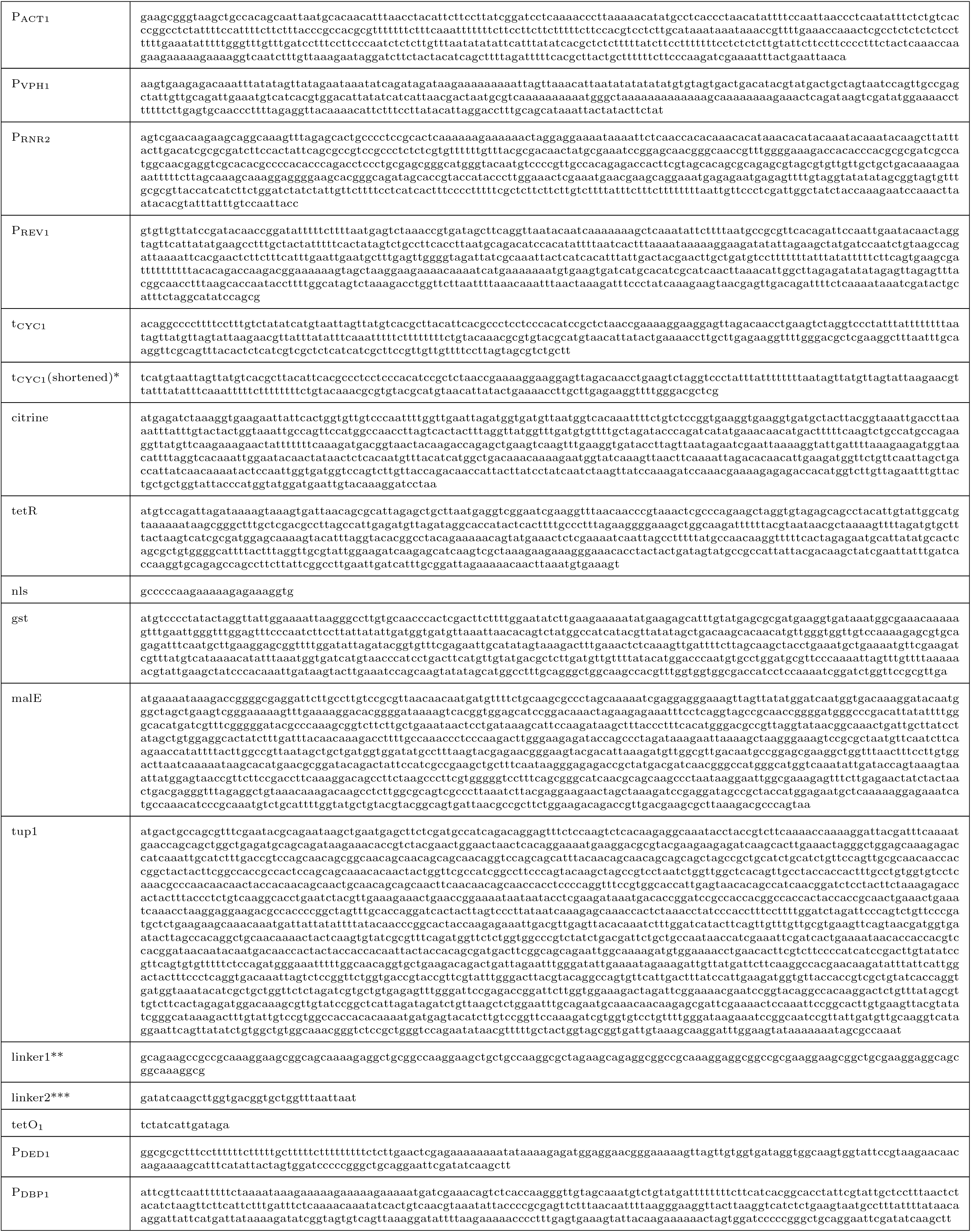
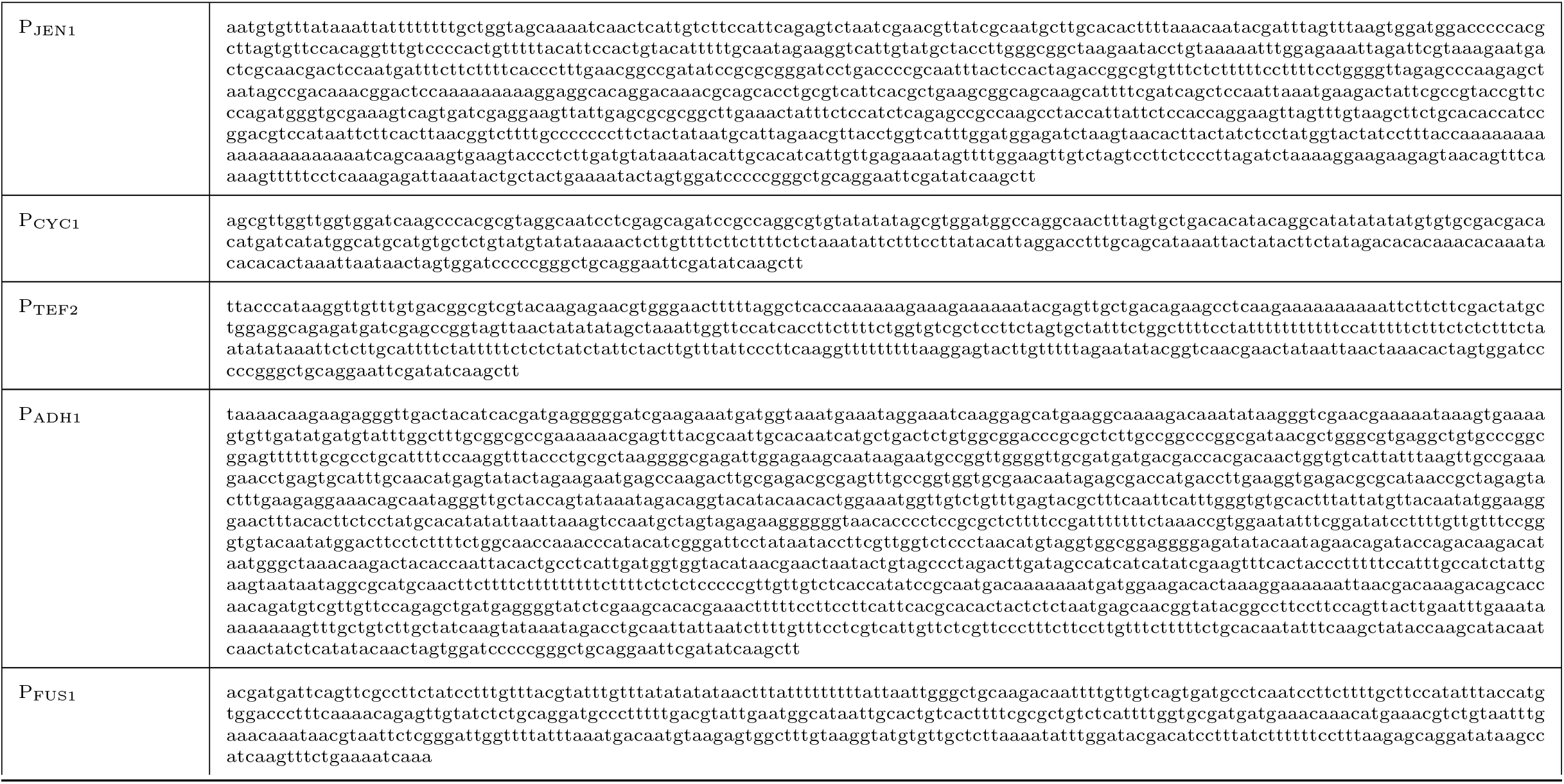
Sequences used in this study. (*) indicates a shortened tCYC1 used to avoid homology in plasmids where there are more than one tCYC1 sequences. (**) indicates the linker sequence used between TetR-nls and the fusion partners MBP and Tup1. (***) indicates the linker sequence used between TetR-nls and the fusion partner GST.

**Table S5:** Parameters used to fit 5-parameter sigmoid curves to experimental data. See Materials&Methods for the 5-parameter log logistic forumula.

| Figure | Curve | p1 | p2 | b | c | d | e |
| --- | --- | --- | --- | --- | --- | --- | --- |
| 2B | SR | -3 | 3 | 38.34324 | 4243.43 | 78857.94 | -0.04581 |
|  | AR | -0.8 | -0.5 | 27.12798 | 7863.577 | 202723.7 | -14.6903 |
|  | cAR | -3 | -0.8 | 84.16559 | 5818.268 | 402084.3 | 3.999615 |
| 3C | P<sub>ACT1</sub> | -3 | 3 | 23.82371 | 256.7728 | 37644.58 | -0.08909 |
|  | P<sub>VPH1</sub> | -3 | 3 | 8.739237 | 177.5421 | 33594.13 | -0.10418 |
|  | P<sub>RNR2</sub> | -3 | 3 | 2.669754 | 471.4979 | 35332.46 | -0.1217 |
| 4B | 30 | -0.8 | 3 | 5.042134 | 450 | 3635.025 | -0.02048 |
|  | 60 | -0.9 | 3 | 5.647304 | 450.0008 | 13159.54 | -0.0119 |
|  | 90 | -0.8 | 3 | 5.749193 | 450.001 | 19744.46 | -0.01477 |
|  | 120 | -0.8 | 2.7 | 6.259459 | 450.0114 | 27467.93 | -0.02503 |
|  | 150 | -0.8 | 3 | 6.054607 | 450.0018 | 29866.02 | -0.01622 |
|  | 180 | -0.8 | 3 | 6.101549 | 450.0153 | 34213.67 | -0.01615 |
|  | 240 | -0.8 | 3 | 6.034959 | 450.0343 | 40824.04 | -0.01712 |
|  | 300 | -0.8 | 3 | 5.995046 | 450.0013 | 41711.19 | -0.01808 |
|  | 360 | -0.8 | 3 | 6.017621 | 450.0001 | 45759.42 | -0.01791 |
|  | 420 | -0.8 | 3 | 6.084224 | 450.0012 | 51293.96 | -0.01874 |
| S10A | SD min | -0.8 | 3 | 3.115107 | 398.4286 | 42718.77 | -0.05582 |
|  | SD Pro | -0.8 | 3 | 1.710265 | 215.6653 | 33137.52 | -0.10014 |
|  | S Eth | -0.8 | 3 | 3.825844 | 356.4444 | 19760.93 | -0.15743 |
|  | S Fru | -0.8 | 3 | 4.547801 | 340.8464 | 63715.13 | -0.03714 |
|  | S Gly | -0.8 | 3 | 3.295805 | 942.3264 | 43800.35 | -0.06344 |
|  | YP Eth | -0.8 | 3 | 5.195388 | 445.1206 | 18735.67 | -0.02659 |
|  | S Raf | -0.8 | 3 | 5.841324 | 247.2277 | 77302.33 | -0.01523 |

## Protocol for WTC_846_ strain generation

## Introduction

The Well-tempered Controller_846_ (WTC_846_) is a two unit transcriptional control system for *S. cerevisiae* (Figure 1A). An inducible promoter (*P_7tet.1_*) is placed in front of the Gene of Interest (GOI). The promoter is based on an engineered version of the strong constitutive promoter of *TDH3*. It was made repressible by placing TetR binding sites next to the binding sites for the transcriptional machinery. As a result, binding of the TetR protein can prevent binding of the endogenous proteins which normally drive transcription. The repressors TetR and TetR-Tup1 are found on one integrative repressor plasmid (Figure 2A). TetR is expressed under the above described promoter (*P_7tet.1_*) creating an autorepression loop. TetR-nls-Tup1 abolishes the basal activity of *P_7tet.1_* and is expressed under the control of the weak, constitutive *RNR2* promoter.

**Figure 1:**
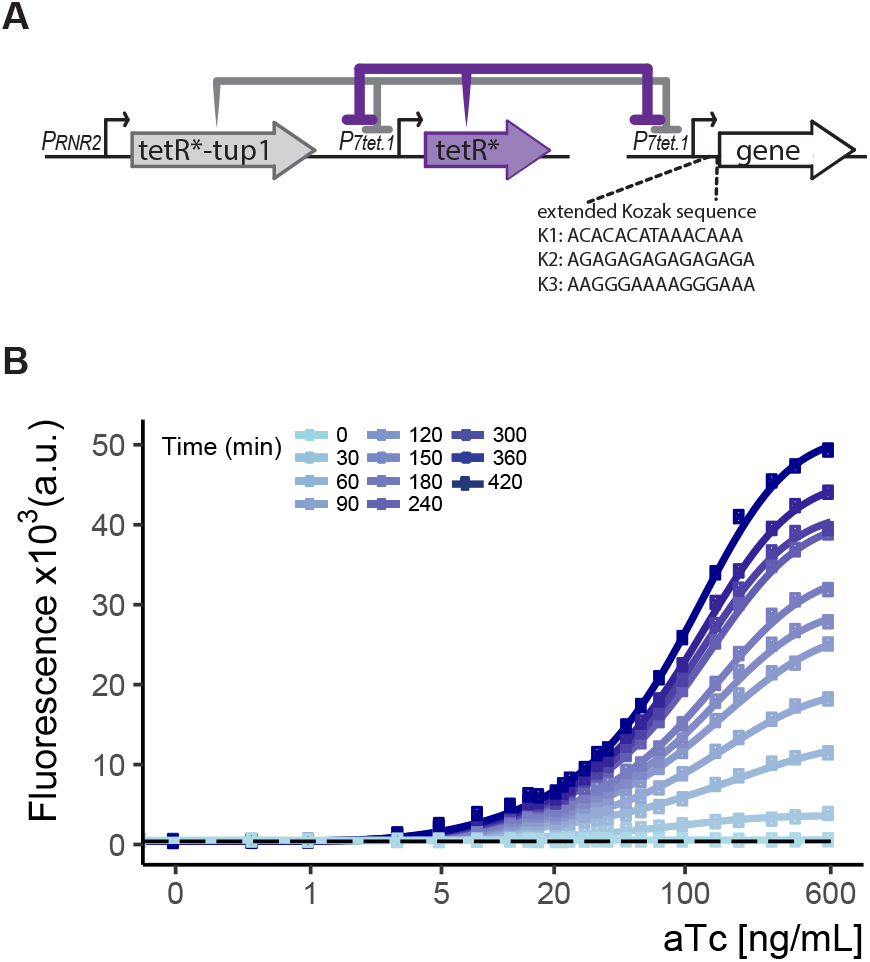
The configuration of and an example of gene expression control by WTC_846_. A) Genetic elements of the WTC_846_ controller. On the integrative plasmid, TetR is driven by the *P_7tet.1_*, TetR-nls-Tup1 is driven by the *RNR2* promoter. The promoter of the gene of interest is replaced with *P_7tet.1_* in the genome. B) WTC_846_ controlled Citrine expression. Flow cytometry measurements from a strain where WTC_846_ regulates expression of Citrine. Anhydrotetracycline (aTc) was added to exponentially growing cells, and samples were taken every 30 minutes for flow cytometry analysis. Circles represent the median of the fluorescence signal, lines were fitted. The dashed line indicates autofluorescence control, i.e. the parent strain without any Citrine integrated.

**Figure 2:**
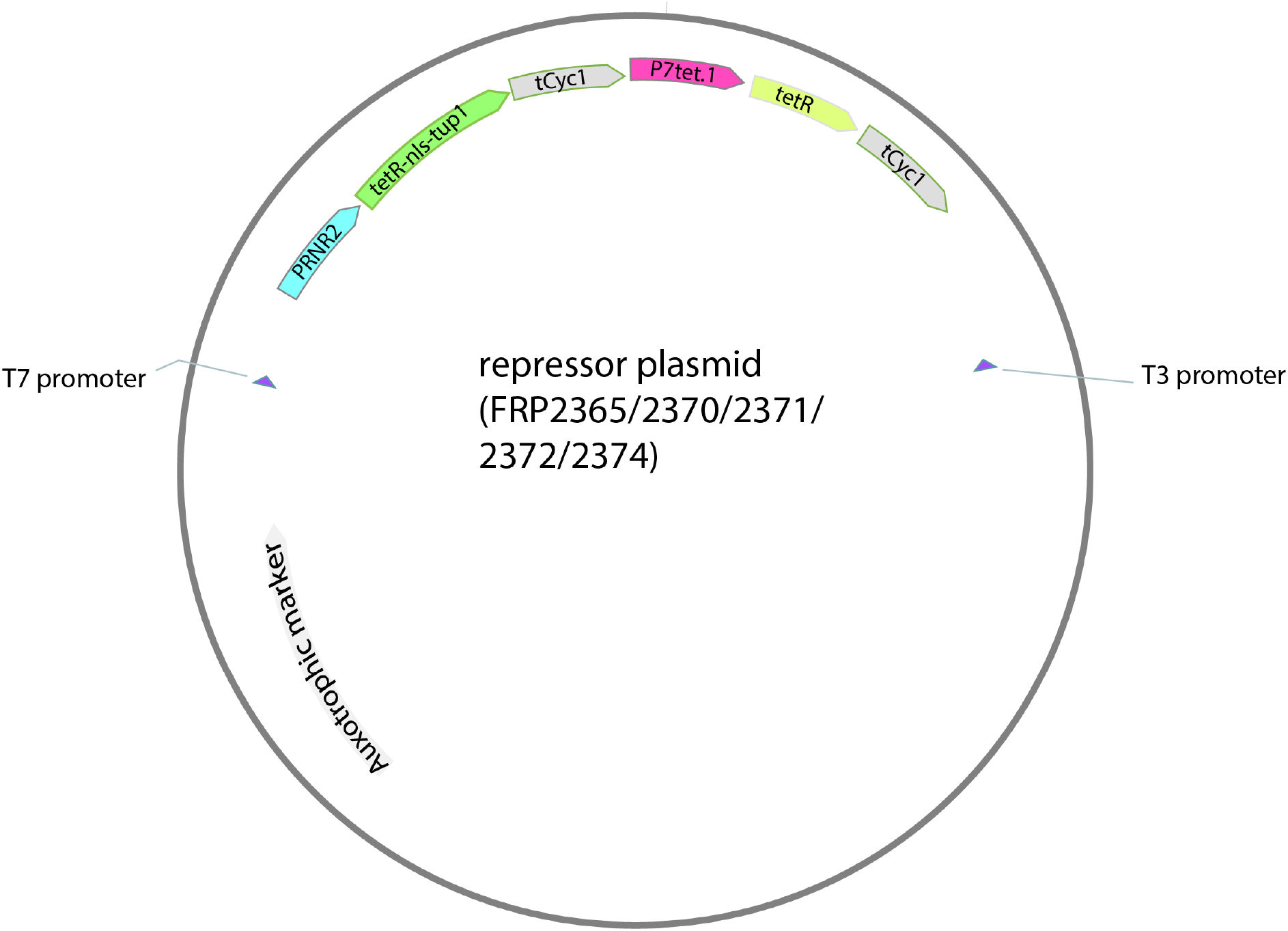
Map of the repressor plasmid. Auxotrophic marker is different depending on the plasmid backbone.

To create a functional system, we advise to first integrate the repressor plasmid. The *P_7tet.1_* can then be placed in front of any gene in the genome using PCR tagging[1]. The tagging plasmid (based on [1]) is used as a template (Figure 3). We provide two versions of the *P_7tet.1_* followed by a flag tag followed by a linker composed of 8 glycine residues; either cloned in a plasmid providing a HygR marker (FRP2350), or a NAT marker (FRP2375). For PCR based tagging, the 5’ and 3’ ends of the PCR fragment need to be complementary to a sequence upstream of the GOI and to the beginning of the GOI, respectively. This is ensured by using primers with tails complementary to these regions. We tested the plasmid for use with and without the flag tag.

**Figure 3:**
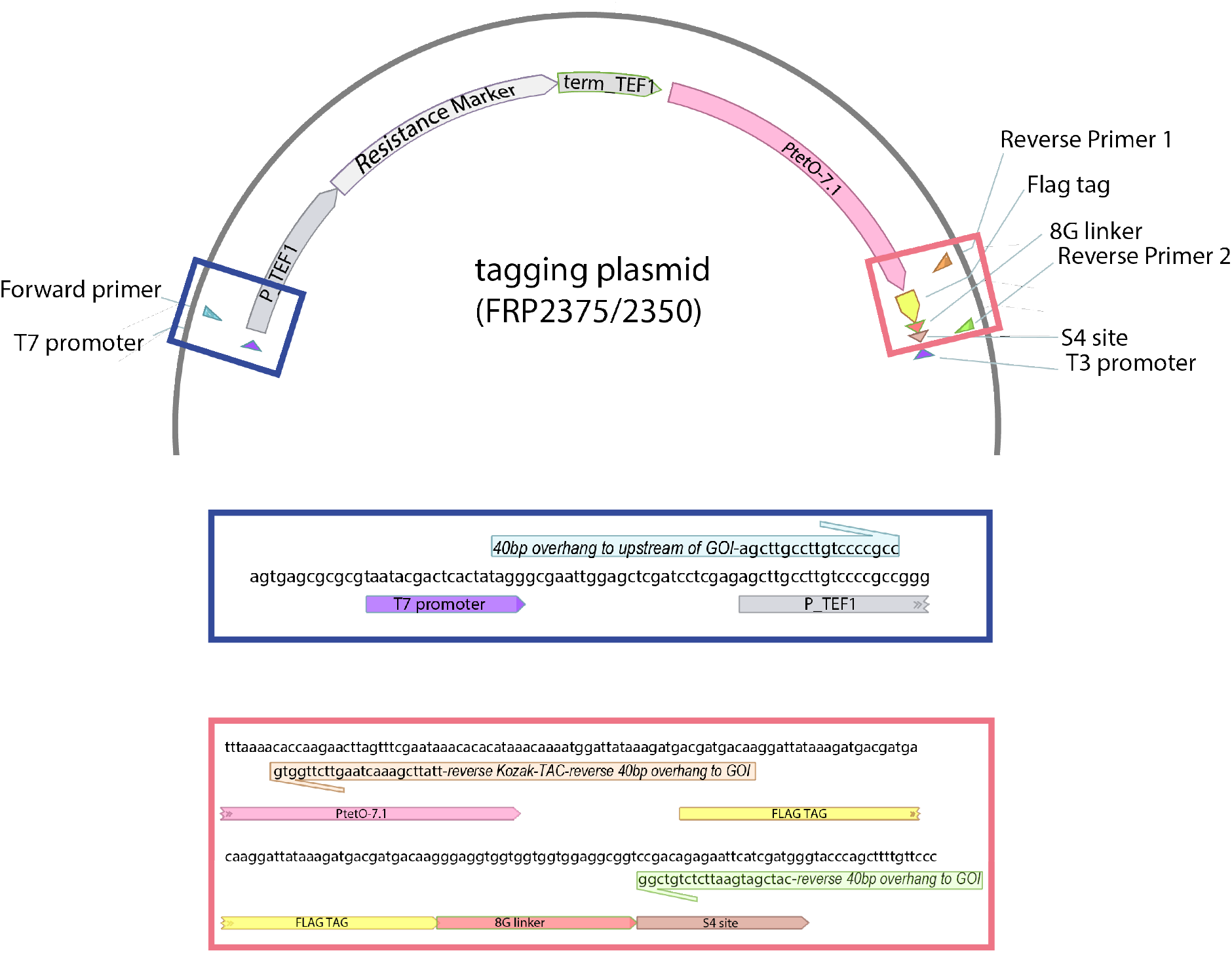
Map of the tagging plasmid. Resistance marker is NAT or HygR depending on the plasmid number. The coloured boxes zoom in to the marked regions to demonstrate how the primers anneal to the plasmid.

Induction of the tagged gene can then be controlled by aTc, a small molecule that causes TetR to dissociate from its binding sites on *P_7tet.1_*. An example is seen in Figure 1B, where Citrine expression was controlled across a large expression range using aTc. At this point we advise to use aTc and not Tetracycline or Doxycycline. While these can be used, they will likely require different concentrations compared to aTc.

The basal activity of *P_7tet.1_* can be controlled by the Kozak sequence (last 15 bp before the start codon of the gene of interest). The provided sequence in the FRP2350 and FRP2375 plasmids shows no detectable basal Citrine expression. However, even a small basal expression level can become an issue if the GOI encodes a protein that is required in very small numbers. We encountered this problem with Tor2, Cdc28 and similarly low abundance, stable proteins. In this case changing the translation efficiency by modifying the Kozak sequence allowed us to abolish all basal expression. The protocol below also explains how to achieve this.

## Tagging Protocol

1. Transform the repressor plasmid in a strain that has the correct auxotrophic marker deletion. The plasmid should be linearized using AscI digestion for integrative transformation [2]. The repressor plasmids are given in Table 1:
2. Design primers to create the tagging fragment from the tagging plasmid (FRP2350 or FRP2375).

- *Forward primer:* Use the sequence agcttgccttgtccccgcc as the annealing part of the forward primer. Select 40 base pairs anywhere upstream of the GOI, and use this sequence as the 5’ tail of your forward primer. Remember that the region between these 40 base pairs and the start codon of the gene will be deleted during the transformation. You can thus remove the entire natural promoter of the gene, but this is not mandatory.
- *Reverse primer option 1 - without flag tag:* Take tttattcgaaactaagttcttggtg as the annealing portion of your reverse primer, which will anneal to the sequence caccaagaacttagtttcgaataaa on the plasmid. Then use the reverse complement of the first 40 base pairs (including ATG) of the GOI, followed by the reverse complement of the desired Kozak sequence as your 5’ tail, such that the primer reads: 5’-reverse GOI sequence-CAT-reverse Kozak sequence-annealing portion-3’.
- *Kozak sequence to modulate expression:* The Kozak sequences that we have tested, in decreasing order of translation efficiency are (reverse complement is given in parentheses):

- ACACACATAAACAAA (TTTGTTTATGTGTGT)
- AGAGAGAGAGAGAGA (TCTCTCTCTCTCTCT)
- AAGGGAAAAGGGAAA (TTTCCCTTTTCCCTT)
- *Reverse primer option 2 - including flag tag:* If you would like to include the flag tag at the start of the gene, use the sequence catcgatgaattctctgtcgg as the annealing portion of your reverse primer, which will anneal to the standard S4 primer binding site on the plasmid (ccgacagagaattcatcgatg). In this case the Kozak sequence cannot be altered, and the one already on the tagging plasmid has to be used (this is the first one in the list above). Use the reverse complement of the first 40 bases (after ATG) of the gene as the 5’ tail of your reverse primer such that the primer reads: 5’-reverse GOI sequence-annealing portion-3’.
3. Perform the tagging PCR to generate the tagging fragment. Use the primers designed in the previous step and the PCR protocol detailed below (adapted from [1]):

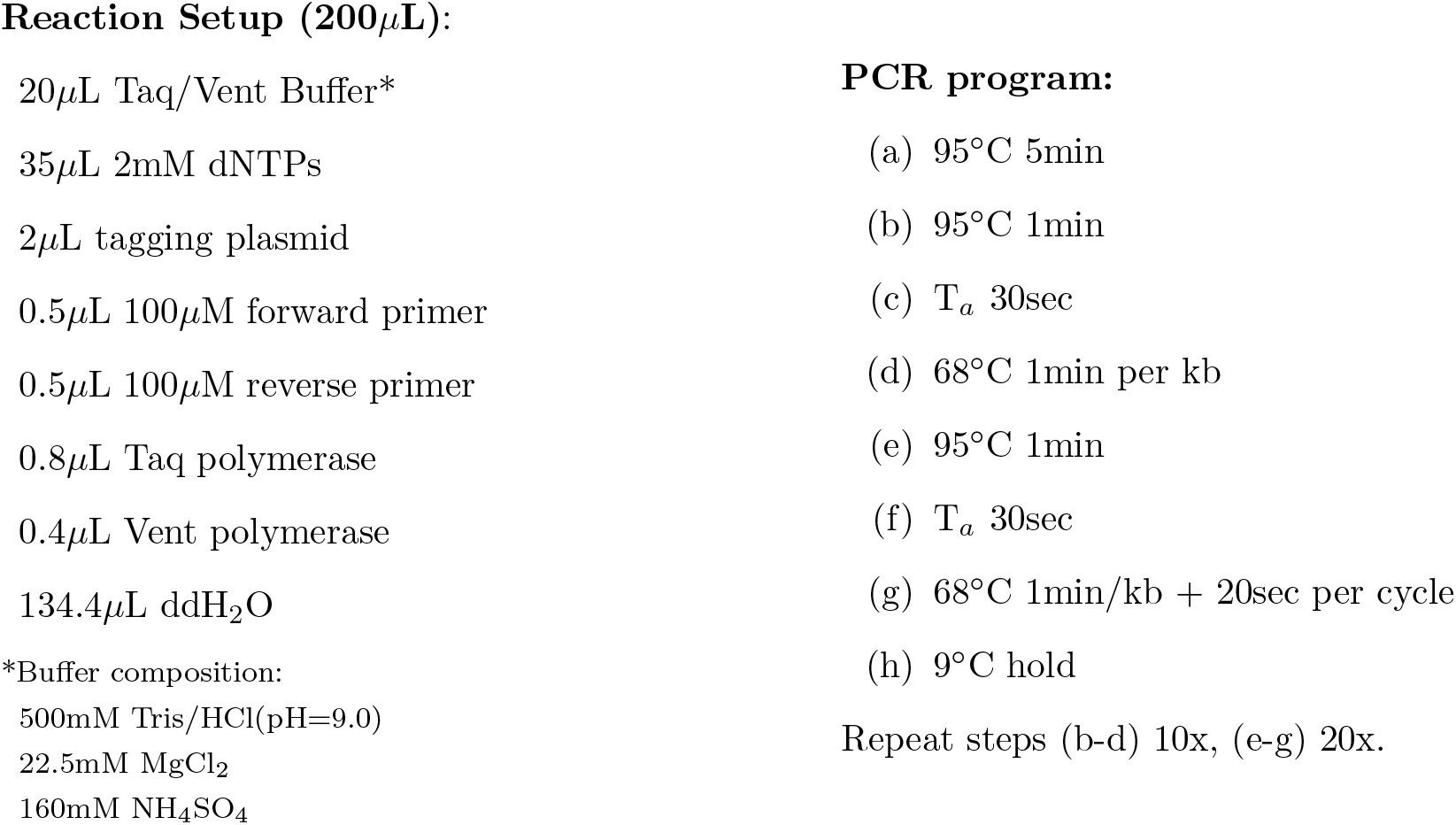
4. Gel isolate and transform the tagging fragment into the strain created in step 1. Select on solid medium with the appropriate antibiotic and aTc. If the GOI is an essential gene, the transformation efficiency will be low. In order to increase transformation efficiency, pre-culture, recovery media for the cells and selection plate should all contain aTc.
5. Correct integration can be confirmed using colony PCR. Use sequence cagttcgagtttatcattatcaatactg as the forward primer (binds at the start of *P_7tet.1_*), and a reverse primer that anneals within the GOI. The fragment length will depend on where in the GOI the reverse primer anneals. (This forward primer will work for all cases except when the *TDH3* promoter is being replaced. Since *P_7tet.1_* is based on the *TDH3* promoter, this primer will anneal to the promoter whether or not the replacement successful.) Integration efficiency is low when tagging essential genes (about 10% of colonies screened), but a positive PCR result generally is enough to indicate correct integration. However it is best to isolate the PCR fragment and sequence the entire promoter to confirm correct integration.

**Table 1:** Repressor plasmids

| FRP Number | Marker | pRG Number[2] |
| --- | --- | --- |
| FRP2365 | URA3 | pRG206 |
| FRP2370 | LEU2MX | pRG205MX |
| FRP2371 | HIS3MX | pRG203MX |
| FRP2372 | LYS2 | pRG207 |
| FRP2374 | MET15 | pRG201 |

## Footnotes

1 Bach, Johann Sebastian, 1685-1750. The Well-Tempered Clavier. Book I: 24 Preludes and Fugues, BWV 846, C Maj

